# Structural Insight into the Function of Human Peptidyl Arginine Deiminase 6

**DOI:** 10.1101/2024.06.10.598250

**Authors:** Jack P. C. Williams, Stephane Mouilleron, Rolando Hernandez Trapero, M. Teresa Bertran, Joseph A. Marsh, Louise J. Walport

## Abstract

Peptidyl arginine deiminase 6 (PADI6) is vital for early embryonic development in mice and humans, yet its function remains elusive. PADI6 is less conserved than other PADIs and it is currently unknown whether it has a catalytic function. Here we have shown that human PADI6 dimerises like hPADIs 2-4, however, does not bind Ca^2+^ and is inactive in *in vitro* assays against standard PADI substrates. By determining the crystal structure of hPADI6, we show that hPADI6 is structured in the absence of Ca^2+^ where hPADI2 and hPADI4 are not, and the Ca-binding sites are not conserved. Moreover, we show that whilst the key catalytic aspartic acid and histidine residues are structurally conserved, the cysteine is displaced far from the active site centre and the hPADI6 active site pocket appears closed through a unique evolved mechanism in hPADI6, not present in the other PADIs. Taken together, these findings provide insight into how the function of hPADI6 may differ from the other PADIs based on its structure and provides a resource for characterising the damaging effect of clinically significant *PADI6* variants.

## Introduction

Peptidyl arginine deiminase 6 (PADI6) is a poorly understood member of the PADI family that is crucial for early embryo development in mice and humans. 32 *PADI6* variants have been reported in 26 infertile women, with embryos from 24 of the women found to arrest at the 4- to 8-cell stage [1–10]. A further 14 *PADI6* variants have been reported in 9 fertile women whose children are often born with multi-locus imprinting disorders (MLID) [11–15]. In mice, *Padi6* knock-out females are infertile, with their embryos arresting at the 2-cell stage [16]. The molecular mechanisms of PADI6 that contribute to these observed phenotypes are poorly understood, however.

Canonical members of the PADI family catalyse the post-translational conversion of peptidyl arginine residues to citrulline in a process known as citrullination (Fig 1A). However, PADI6 does not yet have a confirmed catalytic function, with its classification as an arginine deiminase based on sequence conservation and genomic co-localisation with the other PADI family members [17,18]. Evidence for citrullination by PADI6 in the mouse embryo is inconclusive. In one work, *Padi6* knock-out ovaries showed a decrease in immunohistochemical staining compared to wild-type ovaries when probed with an antibody for citrullinated histone 4 peptide (H4Cit3) [16]. In a later work, however, when wild-type and *Padi6* knock-out ovaries were stained with antibodies to a range of citrullinated histone sequences, including H4Cit3, no difference was observed [19]. Recombinantly expressed mouse PADI6 was enzymatically inactive against the standard PADI substrate benzoyl-L-arginine ethyl ester in the citrulline detecting COlor DEvelopment Reagent (COLDER) assay [20,21]. Additionally, it has recently been reported that recombinant hPADI6 is not active against histone H3 or cytokeratin 5, known substrates of hPADI4 [22].

**Fig 1.**
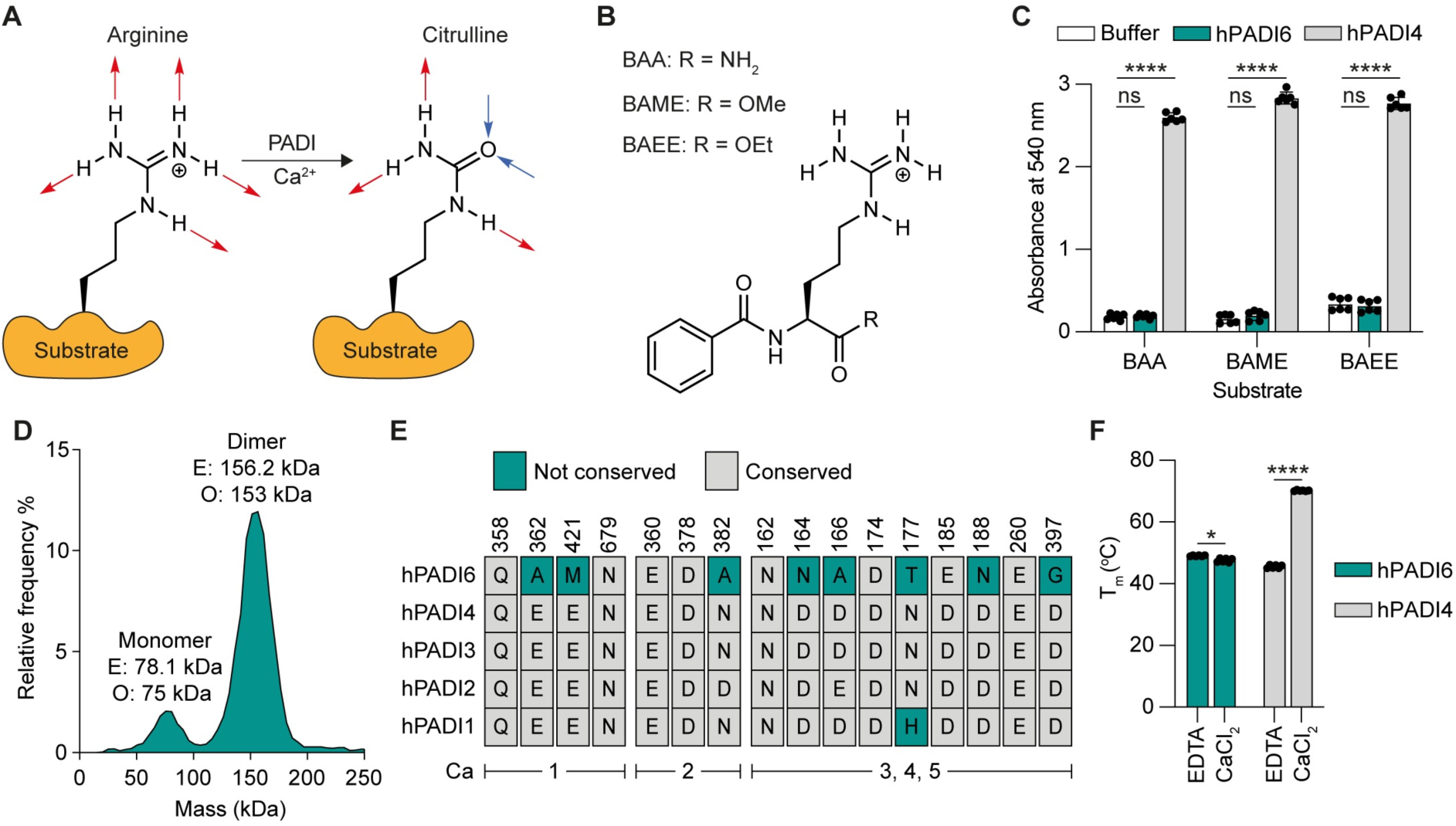
Biophysical and enzymatic characterisation of recombinant hPADI6. (A) Scheme of the PADI-catalysed post-translational conversion of the positively charged arginine to the neutral citrulline. Red arrows = hydrogen bond donors, blue arrows = hydrogen bond acceptors. (B) Structure of standard PADI substrates Nα-benzoyl-l-arginine amide (BAA), Nα-benzoyl-l-arginine methyl ester (BAME) and Nα-benzoyl-l-arginine ethyl ester (BAEE). (C) Activity of hPADI6 or hPADI4 with standard PADI substrates depicted in (B) measured using COLDER assays. Reactions performed in 10 mM CaCl_2_ and quenched after 1 h incubation at RT. [hPADI6] = 500 nM, [hPADI4] = 50 nM, [substrate] = 10 mM. Unpaired parametric t-test, **** = p < 0.0001. 2 independent replicates of 3 technical replicates performed. (D) Mass photometry histogram of hPADI6 expressed from Expi293 cells showing that hPADI6 mainly exists as a dimer *in vitro*. Bin size = 5 kDa. (E) Conservation of Ca^2+^ binding residues in human PADI enzymes, grouped by Ca site. Residue number in hPADI6 highlighted above. Not-conserved = teal, conserved = grey. (F) NanoDSF determined melting temperature (T_m_) of hPADI6 and hPADI4 in either 10 mM EDTA or 10 mM CaCl_2_. Unpaired parametric t-test, **** = p < 0.0001. 2 independent replicates of 3 technical replicates performed.

Despite overall sequence homology with the other PADIs, PADI6 is the least conserved and possesses some key sequence differences to the other PADIs. Human PADIs 1 to 4 bind between 4 and 6 Ca^2+^ ions at defined binding sites (Ca1-6) [23–26]. Sequential Ca^2+^ coordination induces structural changes that form the active site cleft and form catalytically competent enzyme. Thus, high concentrations of calcium are required for *in vitro* enzymatic activity [24,26]. The residues involved in Ca^2+^ binding are highly conserved across the human PADIs, excluding hPADI6. This has led to speculation that hPADI6 does not bind and is not activated by Ca^2+^ [27].

The molecular mechanisms surrounding the function of PADI6 are poorly understood, however a growing body of evidence supports a structural role for PADI6 in early embryo development. It was first reported in 2007 that PADI6 was critical for the formation and/or maintenance of an oocyte and embryo specific cytoskeletal structure known as cytoplasmic lattices (CPLs) [16]. Since then, several studies have reported that the CPLs were potentially composed of, and act as storage sites for, maternal ribosomes and mRNA, as well as being involved in organelle localisation and symmetric division [28–31]. Jentoft et al. recently confirmed that the CPLs were composed of protein fibres containing PADI6, along with members of an oocyte and embryo specific protein complex, the sub-cortical maternal complex (SCMC) [32]. Given the CPLs are absent in *Padi6* knock-out oocytes, it is therefore highly likely that PADI6 is a key structural component of the CPLs in mouse oocytes. In the same work, CPL-like structures were also observed in human oocytes suggesting that this function of PADI6 in the formation and structural organisation of the CPLs is conserved between mice and humans.

Here we recombinantly expressed and purified hPADI6 and confirmed that it is not catalytically active under standard PADI citrullination conditions, it forms a dimer and is not stabilised by Ca^2+^ *in vitro*. To gain insight into how the lack of conservation in hPADI6 affects its structure, and what effect clinically significant variants are having on its structure, we have determined the crystal structure of hPADI6. Analysis of this structure highlighted key differences to the other PADIs which have significant implications on a possible catalytic function of hPADI6. Finally, we use our structure to predict the damaging effect of clinically significant variants on the structure and highlight potentially interesting variants which could be used for further study of the function of hPADI6.

## Results and discussion

### Expression and biophysical characterisation of hPADI6

We first expressed human PADI6 (hPADI6) using the mammalian Expi293 system and purified the recombinant protein to homogeneity by Strep-Tactin® affinity chromatography followed by size exclusion chromatography (S1A-S1E Figs). A single protein of approximately 80 kDa was isolated and confirmed to be hPADI6 by intact mass spectrometry (S1F Fig). Using an established *in vitro* PADI activity assay, the COlor DEvelopment Reagent (COLDER) assay [20], we probed possible enzymatic activity of the recombinant hPADI6 using three standard PADI substrates, Nα-benzoyl-l-arginine ethyl ester (BAEE), Nα-benzoyl-l-arginine methyl ester (BAME) and Nα-benzoyl-l-arginine amide (BAA) (Fig 1B). hPADI6 showed no citrullination activity (Fig 1C). hPADI2 to 4 have been shown to form stable dimers in solution, with dimerisation enhancing citrullination activity of hPADI4 [24–26,33]. Furthermore, it has been reported that recombinant mouse PADI6 forms oligomers up to hexamers in a chemical cross-linking experiment [21]. We therefore characterised possible hPADI6 dimerisation by mass photometry showing that hPADI6 formed a dimer in solution at an approximate ratio of 8:1 dimer:monomer at a concentration of 4 nM (Fig 1D).

### hPADI6 is not stabilised by Ca^2+^

We next characterised the Ca^2+^ binding capacity of hPADI6. The Ca^2+^-free structures of hPADI2 and hPADI4 contain many disordered regions, in particular loops surrounding the Ca^2+^ binding sites that become structured upon Ca^2+^ binding. The structural rearrangements induced by Ca^2+^ binding in hPADI4 have been shown to dramatically improve thermal stability [34]. 8 of the 16 residues directly involved in Ca^2+^ coordination in hPADI4 however, are not conserved in hPADI6, and none of the calcium binding sites (Ca1-5) are fully intact in hPADI6 (Fig 1E). Using differential scanning fluorimetry, the melting temperature (T_m_) of hPADI6 was measured with either EDTA or CaCl_2_. hPADI6 showed no increase in T_m_ in the presence of either CaCl_2_ or EDTA where hPADI4 was stabilised by 25.1 °C (Fig 1F), suggesting that hPADI6 does not bind calcium ions.

### Structure determination

To investigate how the lack of conservation in the PADI6 sequence impacts its structure, and to gain insight into the molecular function of PADI6, we determined the crystal structure of hPADI6 at a resolution of 2.44 Å (Fig 2A, Table 1). Like hPADI2 and hPADI4, hPADI6 crystallised as a head-to-tail homodimer (Fig 2B).

**Fig 2.**
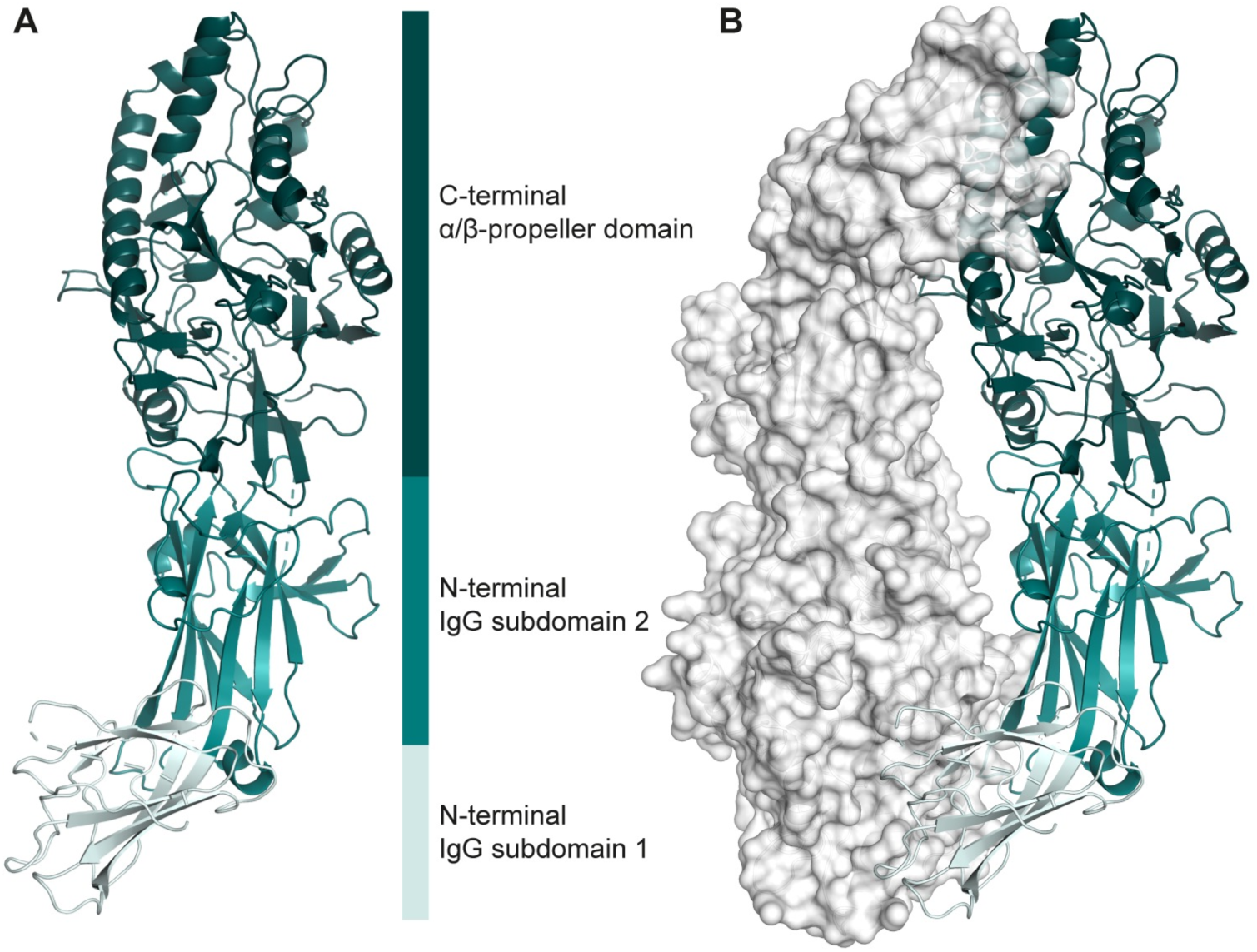
Overall structure of hPADI6. (A) Ribbon representation of monomeric hPADI6. N-terminal IgG subdomains 1 and 2 as well as the C-terminal α/β-propeller domain are light teal, teal and deep teal respectively. Flexible loops that could not be modelled are represented by dashed lines. (B) Structure of the hPADI6 dimer with chain A represented with a ribbon and the surface of chain B displayed. Colouring the same as in (A).

**Table 1.** Crystallographic data and refinement statistics.

|  | **hPADI6** |
| --- | --- |
| Resolution range | 54.06 - 2.44<br>(2.52 - 2.44) |
| Space group | P2<sub>1</sub>2<sub>1</sub>2<sub>1</sub> |
| Unit cell a, b, c | 104.04 123.33 126.54 |
| $\alpha, \beta, \gamma$ | 90 90 90 |
| Total reflections | 846 213 (75 082) |
| Unique reflections | 65 221 (6 051) |
| Multiplicity | 13.8 (12.4) |
| Completeness (%) | 99.20 (96.02) |
| Mean I/sigma(I) | 4.98 (0.24) |
| Wilson B-factor | 58.14 |
| R-merge | 0.36 (5.62) |
| R-meas | 0.37 (5.86) |
| R-pim | 0.10 (1.654) |
| CC1/2 | 0.99 (0.40) |
| Reflections used in refinement | 60 731 (5 810) |
| Reflections used for R-free | 3 027 (322) |
| R<sub>work</sub> | 0.25 (0.47) |
| R<sub>free</sub> | 0.30 (0.49) |
| Number of non-hydrogen atoms | 10 224 |
| macromolecules | 10 224 |
| ligands | 0 |
| solvent | 0 |
|  | 1 329 |
| Protein residues |  |
| RMS(bonds) | 0.004 |
| RMS(angles) | 0.61 |
| Ramachandran favoured (%) | 96.65 |
| Ramachandran allowed (%) | 3.27 |
| Ramachandran outliers (%) | 0.08 |
| Clash score | 8.09 |
| Average B-factor | 78.28 |

The dimer interface area was calculated using PDBe PISA [35] to be 2035 Å^2^, slightly lower than apo-hPADI2 (2551 Å^2^, PDB: 4N20) [24] and apo-hPADI4 (2315 Å^2^, PDB: 1WD8) [26]. hPADI6 possessed a similar domain architecture to hPADI2 and hPADI4, with two N-terminal IgG sub-domains from residues 1-121 and 122-303 respectively, and a C-terminal α/β-propeller domain from residues 304-694. Within the hPADI6 C-terminal domain, the five ββaβ modules were structurally intact (S2 Fig). As for hPADI2 and hPADI4, the hPADI6 dimer is formed through contacts between the C-terminal α/β-propeller domain of one monomer and the N-terminal IgG subdomain 1 of the second.

### hPADI6 is structurally ordered around Ca1 and Ca2

Having determined experimentally that hPADI6 does not bind calcium (Fig 1F) we wanted to investigate the structures of these sites. Four loops are highly disordered in the Ca^2+^ free apo-hPADI4 structure and become structured in the catalytically active holoenzyme upon Ca1 and Ca2 coordination, and substrate binding (S312-L320, C337-Q349, S370-D388, and G395-G403, PDB: 1WDA) [26]. These loops are already structured in hPADI6 in the absence of Ca^2+^ (Fig 3A, S3 Fig). Ca1 is coordinated in hPADI4 by two glutamic acid residues initially, followed by a glutamine residue upon Ca2 binding (E353, E411, Q349) [26]. These Ca1 binding glutamic acid residues are not conserved in hPADI6 where they are replaced by a methionine and alanine (A362, M420), rendering Ca1 binding highly unlikely (Fig 3B). Ca2 is coordinated by an aspartic acid, a glutamic acid and an asparagine (E351, D369, N373) in hPADI4 (Fig 3C). Only the asparagine is not conserved in hPADI6 where it is replaced by an alanine (A382). In hPADI4 the equivalent substitution, N373A produces a catalytically dead protein suggesting it is critical for Ca2 coordination and as such it is unlikely hPADI6 would be able to bind Ca2 either [26].

**Fig 3.**
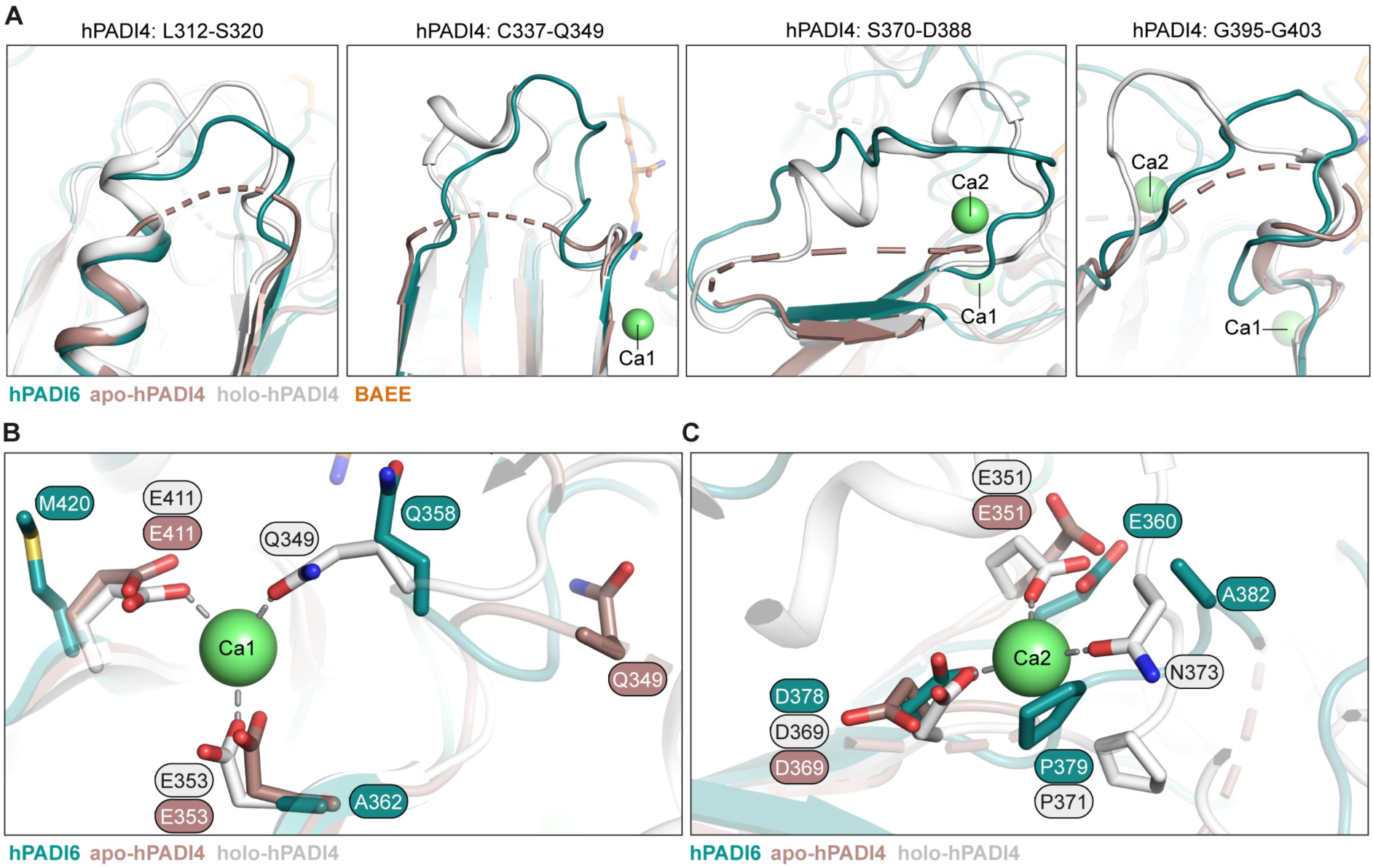
hPADI6 is structured around Ca1 and Ca2 in the absence of Ca^2+^ coordination. (A) Close-up views of unstructured loops in apo-hPADI4 (brown, PDB: 1WD8) that are structured in holo-hPADI4 (white, PDB: 1WDA) aligned with corresponding loops in hPADI6 that are already structured in the absence of Ca^2+^ (teal). Unstructured loops represented by dashed lines and residue ranges of the unstructured loops in apo-hPADI4 are highlighted. (B) hPADI4 Ca1 site in the presence (white, PDB: 1WDA) and absence (brown, PDB: 1WD8) of Ca^2+^, aligned with hPADI6 (teal). hPADI4 Ca1 coordinating residues, and corresponding residues in hPADI6 shown and labelled. Amino acid interactions with Ca1 represented by grey dashed lines. (C) Same as (B) but Ca2 instead of Ca1.

Only one of the disordered regions in hPADI4 that becomes structured upon Ca^2+^ binding was disordered in hPADI6, A166-K178 (S4 Fig). The corresponding region of hPADI4 becomes ordered upon Ca3,4,5 coordination [26]. 5 of the 9 residues involved in coordinating Ca3,4,5 are not conserved in hPADI6 and thus it seems unlikely that hPADI6 could coordinate Ca^2+^ in these sites (Fig 1E).

Given hPADI6 is already structured in the majority of the flexible regions of hPADI4 in the absence of Ca^2+^ and substrate binding, we wondered whether hPADI6 structurally aligns more closely with the inactive Ca^2+^-free PADI2/4 structures, or the active Ca^2+^-bound structures. Using PyMOL, we calculated Root Mean Square Deviation (RMSD) values of our hPADI6 monomer structure compared with various published apo- and holo-hPADI2 and hPADI4 structures (Table 2) (S5 Fig) [24,26]. Interestingly, hPADI6 is similarly different to the holo-hPADI2 and hPADI4 structures compared to the apoenzymes.

**Table 2.** Cα RMSD values of hPADI6 aligned with various hPADI2 and hPADI4 structures.

| PDB | Protein | Crystallisation Complex | hPADI6 RMSD (Å) | Cα |
| --- | --- | --- | --- | --- |
| 4N20 [24] | hPADI2 | Ca free | 1.134 | 441 |
| 4N2B [24] | hPADI2 | Ca1, 3, 4, 5, 6 bound | 1.079 | 434 |
| 4N2C [24] | hPADI2<sup>F211/222A</sup> | Ca bound | 1.018 | 423 |
| 1WD8 [26] | hPADI4 | Ca free | 1.027 | 416 |
| 1WD9 [26] | hPADI4 | Ca bound | 1.002 | 430 |
| 1WDA [26] | hPADI4<sup>C645A</sup> | Ca and BAEE bound | 1.060 | 449 |

RMSD values calculated using PyMOL only counting Cα position. Cα = number of Cα used to calculated RMSD.

### The catalytic aspartic acid and histidine residues of hPADI6 align with Ca^2+^-bound hPADI4

The calcium binding of hPADI2 and hPADI4 occurs sequentially, with first Ca1 binding, followed by Ca3,4,5 and finally Ca2 [24]. It is the coordination of Ca1 and Ca2, and the subsequent structural ordering of the loops surrounding these sites that creates the active site, producing a catalytically competent form of the protein. As hPADI6 is already structurally ordered in the absence of Ca1 and Ca2 coordination, we explored the positioning and structure of key residues in the active site. The PADI catalytic site residues include a key nucleophilic cysteine, two aspartic acids and a histidine. The aspartic acids (D359, D482) and histidine (H480) are conserved in hPADI6, and structurally align with the equivalent residues in both Ca^2+^-bound hPADI2 and hPADI4 (S6 Fig, Fig 4A). This is most clear for D359 where the Cα of the equivalent D350 in hPADI4 moves 3.1 Å upon Ca^2+^ coordination, with Cα distances of hPADI6 to the Ca^2+^-free and bound hPADI4 D350 being 3.6 Å and 0.9 Å respectively.

**Fig 4.**
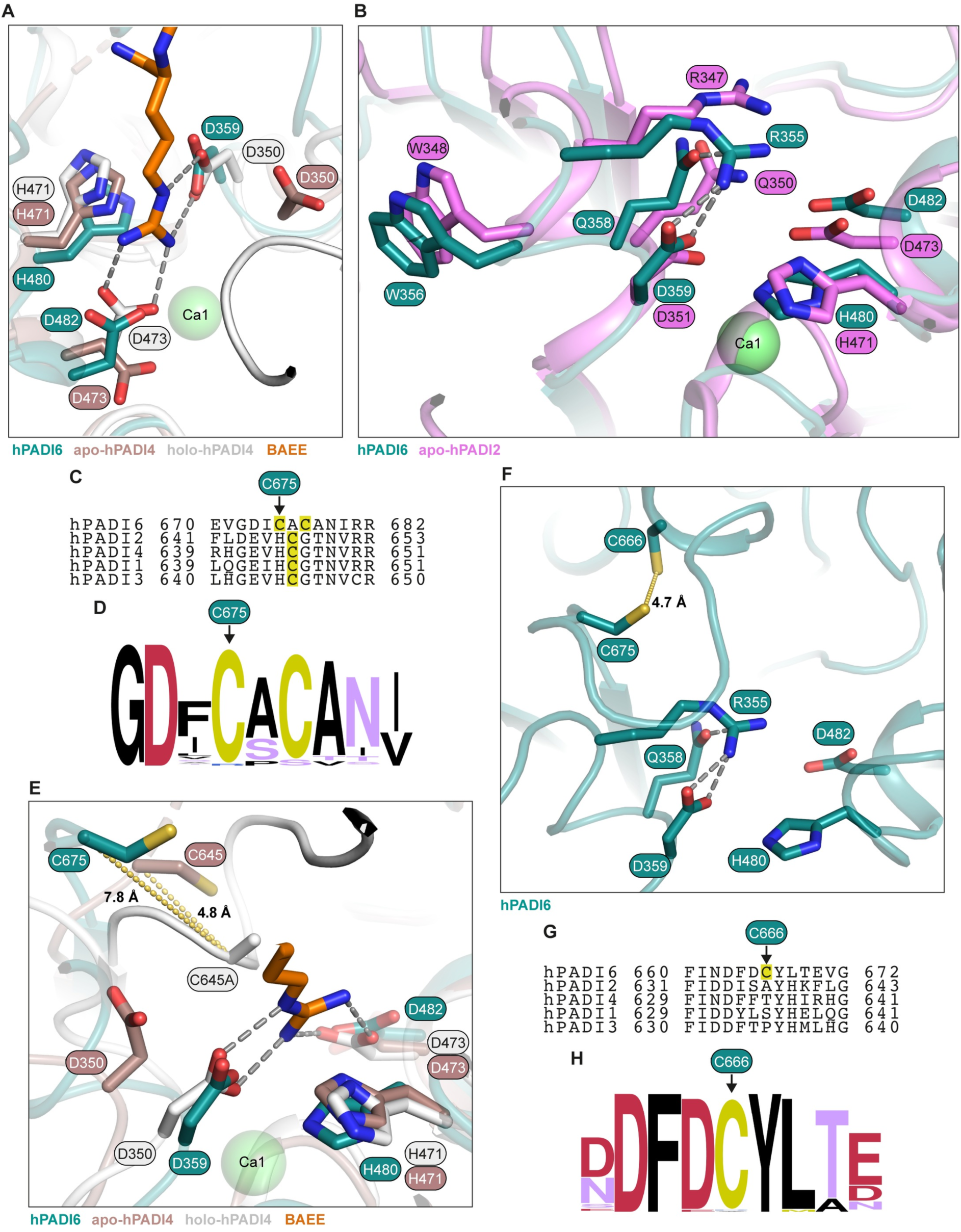
The hPADI6 active site partially aligns with hPADI2 and hPADI4. (A) Close up view of hPADI6 active site residues (teal), aligned with apo-hPADI4 (brown, PDB: 1WD8), and holo-hPADI4 (white, PDB: 1WDA). Substrate BAEE as part of the holo-hPADI4 displayed in orange. (B) Close up view of hPADI6 active site residues (teal), aligned with apo-hPADI2 (pink, PDB: 4N20). Key arginine, glutamine and tryptophan residues, as well as catalytic tetrad residues highlighted. (C) Sequence alignment of the human PADIs centred on the key catalytic cysteine residue. Yellow shading = potential hPADI6 key catalytic cysteine residues and confirmed hPADI1-4 catalytic cysteines. Predicted hPADI6 catalytic cysteine C675 highlighted. (D) Logo plot of the hPADI6 sequence surrounding potential catalytic cysteine residues C675 and C677 aligned with the sequences of PADI6 in 79 other species. For a list of species used see S1 File. Logo plot produced using WebLogo.berkeley.edu [36,37]. (E) Close up view of hPADI6 active site residues (teal), aligned with apo-hPADI4 (brown, PDB: 1WD8), and holo-hPADI4 (white, PDB: 1WDA). Predicted hPADI6 catalytic cysteine shown with distances to confirmed hPADI4 catalytic cysteine C645 (C645A in holo-hPADI4 structure) in the apo and holoenzyme structures. Substrate BAEE as part of the holo-hPADI4 structure displayed in orange. (F) Close-up view of hPADI6 C675 and C666 positioned above the catalytic aspartic acids and histidine. Distance between sulphur atoms of each cysteine = 4.7 Å. (G) Sequence alignment of the human PADIs centred on the hPADI6 C666, highlighted by yellow shading. C666 is not conserved in hPADIs 1 to 4. (H) Logo plot of the PADI6 sequence surrounding hPADI6 C666 aligned with the PADI6 sequences of 79 other species. C666 is conserved in 78 out of 80 aligned sequences. For a list of species used see S1 File. Logo plot produced using WebLogo.berkeley.edu [36,37].

### Arg355 gatekeeps the hPADI6 active site pocket

In the Ca^2+^-free hPADI2 structure, an arginine shields the active site, functioning as an important regulatory element in PADI activity [24]. In Ca^2+^-free hPADI2, R347 points into the active site pocket, stabilised through a hydrogen bond with Q350 (PDB: 4N20). Upon Ca2 binding, R347 flips out of the active site completely and W348 moves in and forms a wall in the active site pocket with Q350 rotating to coordinate Ca1. In hPADI6, R355 is similarly gatekeeping the active site, held in place by a hydrogen bond to Q358 (Fig 4B) and the W356 of hPADI6 mimics the positioning of hPADI2 W348 in the Ca^2+^-free state. The movement of hPADI2 W348 has been shown to be critical for catalytic function as its substitution to alanine dramatically reduces the catalytic activity of hPADI2 [24].

### The predicted catalytic cysteine of hPADI6 is displaced away from the active site center

The catalytic cysteine of hPADIs 1 to 4 is not conserved in hPADI6 but is replaced by an alanine flanked by two cysteine residues (Fig 4C). Both these residues are highly conserved in the PADI6 sequence; the first is conserved in 78 of the 80 complete available PADI6 sequences with the second cysteine being replaced by serine in rodents (Fig 4D, S1 File). Given the first of these cysteines, C675 in hPADI6, is more strongly conserved, we hypothesised that this would be the catalytic cysteine of hPADI6. This is reinforced by the structure of hPADI6, as C677 is likely involved in a disulfide bond with C320, important in stabilising one of the five ββaβ modules of the C-terminal domain (S7 Fig). C675 is still far from the active site centre however and would need to move approximately 9.1 Å to be in a similar position to the catalytic cysteine of Ca^2+^-bound holo-hPADI4 (Fig 4E). C645 of hPADI4 does move 4.5 Å upon Ca2 binding, as such it is conceivable that hPADI6-C675 could move into a similar position above the other active site residues although it is highly unlikely this movement is induced by Ca^2+^ binding. In close proximity to C675 is another cysteine, C666 (distance between sulphur atoms ∼ 4.7 Å) (Fig 4F). Whilst these two cysteines do not appear to be disulfide bonded in the refined structure, hPADI6 was purified and crystallised in the presence of reducing agent TCEP and as such it is conceivable that C675 and C666 could form a disulfide bond *in vivo*. C666 is not conserved in any of the other hPADIs yet is highly conserved in the PADI6 sequence across all mammals (Fig 4G, H). The positioning of C666 and its uniqueness to PADI6 is intriguing and could indicate a possible redox mechanism regulating the accessibility and flexibility of C675.

### The hPADI6 active site cleft is blocked

In the absence of Ca^2+^ binding, the hPADI4 and hPADI2 active site clefts are highly disordered [24,26]. Binding of Ca1 and Ca2 induces structural ordering of this region, creating the active site cleft and allowing the substrate to enter. By contrast, hPADI6 is well ordered in this region (Fig 5A) and a loop (E670–D673, blocking loop) blocks the entrance to the active site cleft, occupying the site bound by BAEE in substrate-bound hPADI4 (Fig 5B, C). The ordered structure and positioning of this loop over the active site cleft is likely maintained by hydrogen bonds between the δ^2^-NH_2_ group of N598 on a neighbouring loop (holding loop) and D673, and the δ^1^-O of N598 with the backbone NH of G672 (Fig 5B). Both D673 and G672 of the blocking loop are fully conserved in the PADI6 sequence from 80 species (Fig 5D). The holding loop is an insertion in the PADI6 sequence, not present in PADIs 1 to 4 (Fig 5E). Whilst there is some variation in the sequence of this loop between the PADI6 sequence of different species, N598 is highly conserved, and present in this position in 79 out of the 80 sequences analysed (Fig 5F). Together this indicates a unique evolved mechanism by which the PADI6 active site is blocked. Interestingly, a compound heterozygous variant in p.N598 (p.N598S/p.R682Q) has been reported in a patient with female infertility [1]. Additionally, a patient carrying homozygous p.I599A has also been reported to be infertile, again with high miscarriage incidence and hydatidiform mole formation [7]. Together these clinical instances suggest this structural feature may have functional relevance.

**Fig 5.**
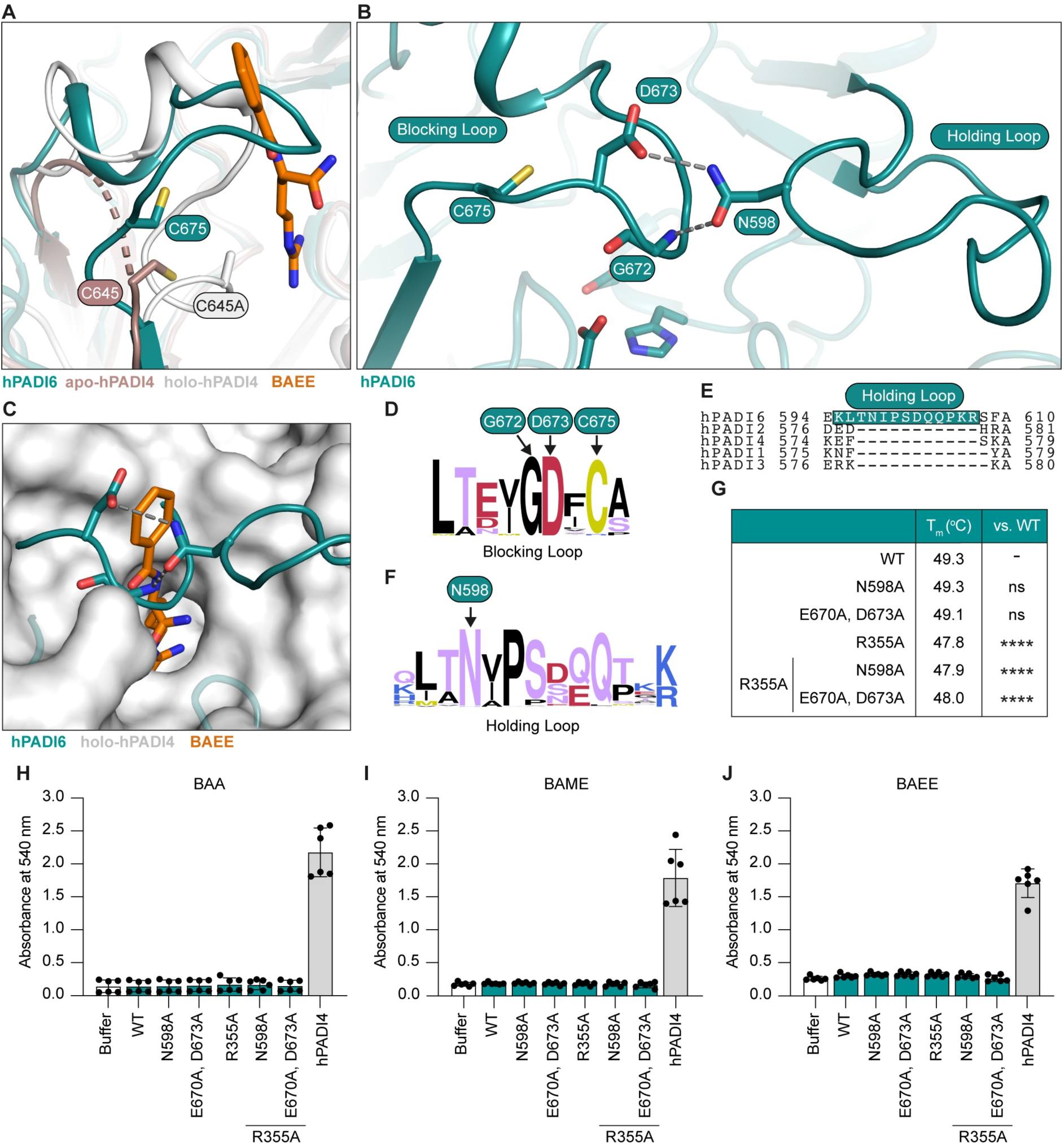
The hPADI6 active site pocket is blocked. (A) Close-up view of unstructured loop D632-C645 in hPADI4 in the absence of Ca^2+^ (brown, PDB: 1WD8) that becomes structured in holo-hPADI4 (white, PDB: 1WDA) aligned with corresponding loop in hPADI6 that is already structured in the absence of Ca^2+^ (teal). Unstructured loops displayed by dashed lines. Substrate BAEE as part of the holo-hPADI4 structure displayed in orange. hPADI4 catalytic cysteine C645 (C645A in holo-hPADI4 structure) and predicted hPADI6 catalytic C675 highlighted. (B) hPADI6 blocking and holding loops showing hydrogen bonds between D673 of the blocking loop and backbone NH of G672, with N598 of the holding loop. Predicted catalytic cysteine C675 is adjacent to the blocking loop. (C) Surface representation of the holo-hPADI4 active site cleft (white, PDB: 1WDA), superimposed with ribbon representation of hPADI6 (teal). Substrate BAEE as part of the holo-hPADI4 structure displayed in orange. hPADI6 D673, G672, and N598 shown with predicted hydrogen bonds represented by grey dashed lines. (D) Logo plot of the hPADI6 blocking loop sequence aligned with the sequences of PADI6 in 79 other species. For a list of species used see S1 File. D673 is conserved in all 80 of the aligned sequences. Logo plot produced using WebLogo.berkeley.edu [36,37]. (E) Sequence alignment of the human PADIs covering the hPADI6 holding loop sequence showing it is an insertion in the PADI6 sequence. hPADI6 holding loop highlighted with grey dashed box. (F) Logo plot of the hPADI6 holding loop sequence aligned with the sequences of PADI6 in 79 other species. For a list of species used see S1 File. N598 is conserved in 79 of the 80 aligned sequences. Logo plot produced using WebLogo.berkeley.edu [36,37]. (G) NanoDSF determined melting temperatures (T_m_) of wild-type (WT) hPADI6 and variants. Unpaired parametric t-test, **** = p < 0.0001. 2 independent replicates of 3 technical replicates performed (H) Activity of WT hPADI6, hPADI6 variants or hPADI4 with PADI substrate BAA determined by COLDER assay. Reactions performed in 10 mM CaCl_2_ and quenched after 1 h incubation at RT. [hPADI6/hPADI6 variant] = 500 nM, [hPADI4] = 50 nM, [substrate] = 10 mM. Unpaired parametric t-test, **** = p < 0.0001. 2 independent replicates of 3 technical replicates performed. (I) As (H) with BAME instead of BAA. (J) As (H) with BAEE instead of BAA.

Given the sequence proximity of the hPADI6 predicted catalytic cysteine (C675) to the blocking loop, we hypothesised that a structural rearrangement of the blocking loop would also move C675. As such, disruption of the hydrogen bonds between the blocking and holding loops could theoretically open the active site pocket and simultaneously move C675 into the active site. To test this hypothesis, five hPADI6 variants were prepared recombinantly. The first two variants aimed to directly disrupt the hydrogen bonding between the loops: one with N598 substituted with alanine (hPADI6^N598A^), and a second with D673 as well as the proximal E670 substituted with alanine residues (hPADI6^E670A, D673A^) (S8 Fig). E670 was substituted as well as D673 due to its proximity to N598 and potential capacity to form a hydrogen bond in the absence of D673. These substitutions would likely not result in the movement of gatekeeper R355 out of the active site pocket, however. To counter this, the hPADI6 variants containing blocking or holding loop substitutions were also recombinantly prepared with R355 substituted to alanine (hPADI6^R355A, N598A^ and hPADI6^R355A, E670A, D673A^), as well as a fifth variant with only R355 substituted to alanine for comparison (hPADI6^R355A^) (S8 Fig). hPADI6^N598A^ and hPADI6^E670A, D673A^ both melted at a similar temperature to the unsubstituted hPADI6 protein (Fig 5G), suggesting disruption of the loop hydrogen bonds do not destabilise the overall folded structure of hPADI6. The three variants containing the R355A substitution were destabilised by an average of 1.3 °C suggesting R355 had some importance in folding interactions, however, the destabilising effect of its substitution is minor. None of the variants displayed any citrullination activity against PADI substrates BAEE, BAME or BAA, however, demonstrating that disruption of this loop region is not sufficient to induce catalytic activity (Fig 5H-J). Without structural information on the loop in each of the variants, we cannot determine whether these substitutions successfully disrupted the interactions and opened the active site pocket or whether the active site pocket was opened but the cysteine was not moved into position. Additionally, it is likely that other structural rearrangements are required such as the movement of W356 to form a wall in the active site pocket, which has been shown to be critical for the catalytic activity of hPADI2 [24].

### Predicting damage score of clinically significant *PADI6* variants

Finally, with a high-resolution X-ray crystal structure of hPADI6 in hand, we investigated the structural consequences of the reported clinically significant variants. A total of 25 variants were mapped onto the hPADI6 structure, 12 of which are associated with early embryonic developmental arrest, 3 of which are associated with extended pregnancies with recurring hydatidiform moles, and 8 that are present in fertile women whose offspring have imprinting disorders (Fig 6A). 2 variants were reported to result in both the early embryonic development arrest and imprinting phenotypes. 19 out of the 25 mapped substitutions were present in the C-terminal domain of hPADI6.

**Fig 6.**
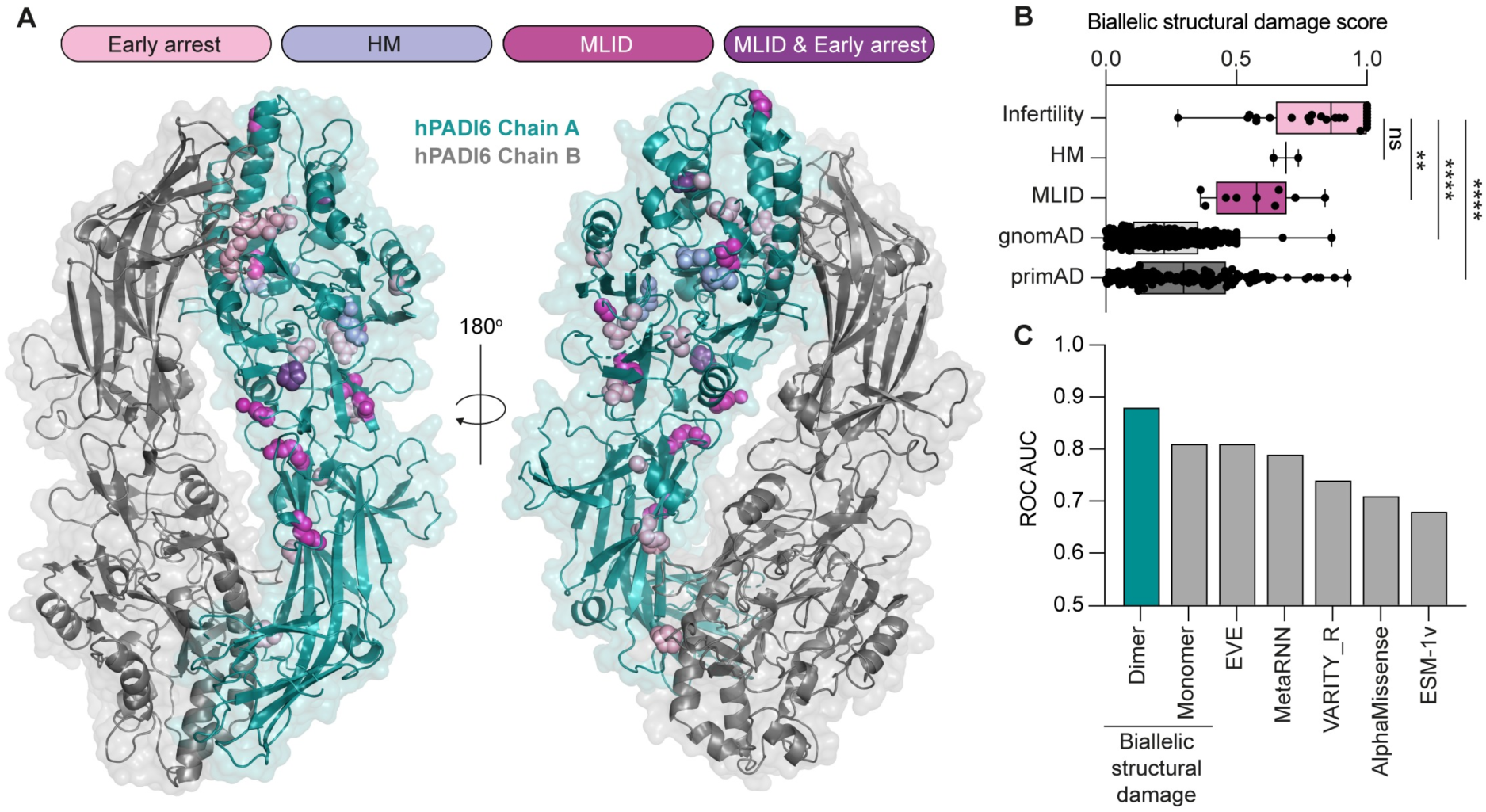
FoldX damage score prediction of clinically significant hPADI6 variants. (A) hPADI6 dimer structure with clinically significant hPADI6 variants highlighted on Chain A as spheres. Variants that result in early embryonic developmental arrest shown in light pink, hydatidiform moles in lilac, multi-locus imprinting disorders (MLID) in offspring in magenta, and variants that have been reported to result in both MLID and early embryonic developmental arrest in purple. (B) Distribution of biallelic structural damage scores, calculated from the full dimer structure, for women with variants associated with infertility, HM and MLID, as well as putatively unaffected controls from gnomAD and primAD. (C) Performance of biallelic damage scores, calculated from the dimeric and monomeric structures, as well as from five sequence-based variant effect predictors, at discriminating between infertility and MLID patients, as measured by ROC AUC, whereby a value of 1 indicates perfect discrimination, and 0.5 would be expected for random chance.

We modelled the effects of all possible hPADI6 missense variants on the stability of the full homodimer and used this to calculate the recently introduced ΔΔG_rank_ metric for each variant, whereby a value of 0 represents the least damaging possible mutations and 1 represents the most damaging variant [38]. Wild-type alleles were given a value of 0 and protein null (*i.e.* truncating) variants were given a value of 1. Next, we scored each patient in terms of their predicted biallelic structural damage to hPADI6 by averaging the two values for each allele. In Fig 6B, we compare the biallelic structural damage scores for the infertility, HM and MLID patients. We also compared heterozygous and homozygous variants from gnomAD v2.1, and homozygous variants from primAD as putatively unaffected controls. Notably, the infertile women tend to have significantly higher structural damage scores than the MLID cases or the controls, suggesting that structural damage to hPADI6 is the primary molecular mechanism underlying infertility-associated missense variants.

Finally, we compared the dimer-based biallelic structural damage scores to equivalent scores calculated from the monomeric subunit structure, and from several state-of-the-art variant effect predictors [39], testing them for their ability to distinguish between infertility and MLID phenotypes as measured by the receiver operating characteristic (ROC) area under the curve (AUC) (Fig 6C). Remarkably, we find that the dimer-based scores show considerably better discrimination, demonstrating the power of considering protein structural context for clinical interpretation of hPADI6 variants. Interestingly, in the infertility group, four of the biallelic structural damage scores were markedly higher for the dimeric structure than the monomeric structure, where on average the other infertility-related variants had comparable biallelic structural damage scores for the monomer and dimer (S9A Fig). Each case possessed either a H211Q, P289L, or E586K causing variant, all of which are localised at the hPADI6 dimer interface (S9B Fig). This highlights the importance of hPADI6 dimerisation to its function in early embryo development.

## Conclusion

Together, this work highlights key differences in the structure of hPADI6 compared with the other hPADIs and provides insight into how its regulation and function differs from the rest of the family. Our structure also provides a useful resource for characterising the effect that clinically significant *PADI6* variants have on its structure and function. The lack of sequence homology between hPADI6 and the rest of the PADIs does not impact its dimerization ability, although it does eliminate the capacity of hPADI6 to bind calcium. The function of dimerization in hPADI6, and also the other PADIs, has yet to be fully elucidated [23–26,33]. Understanding the cellular function of hPADI6 dimerisation could provide insight into the functions of dimerisation in the other PADIs too. In this regard the three potential dimerisation disrupting PADI6 variants identified in this work offer a unique system to studying this further. Perhaps the most intriguing observation from this work is the structure of the hPADI6 active site. The active site pocket of hPADI6 appears blocked through hydrogen bonding between two loops, one of which is an insertion in the PADI6 sequence, not present in the other PADIs. Furthermore, the predicted catalytic cysteine of hPADI6 is displaced away from the active site centre. It is conceivable that a structural rearrangement of the blocking loop could both open the pocket and move the cysteine into an enzymatically active position. A move of such distance is plausible considering that the cysteines of hPADI2 and hPADI4 both undergo dramatic movements upon calcium binding, along with significant structural rearrangements in general [24,26]. Given that hPADI6 does not appear to bind calcium however, it is not clear what could cause such a movement if hPADI6 is indeed enzymatically active. One possibility is a post-translational modification (PTM) to PADI6, especially as there have been reports of PADI6 phosphorylation [40,41]. Interestingly, however, whilst this manuscript was in preparation, a structure of a hPADI6 variant containing two phosphomimetic substitutions, V10E and S446E, was disclosed (PDB: 8QL0) [22]. The published phosphomimetic structure is similar to our wild-type structure (Cα-RMSD of 0.436 Å over 504 atoms). In particular, the active site is also blocked by the same loop, suggesting, at least these PTM mimics are not sufficient to open the active site. An alternative possibility could be through a protein-protein interaction (PPI) around the Ca1 and Ca2 sites that induces similar structural changes to that of Ca2 binding in hPADI2 and hPADI4. Examples of peptide or protein binding at this site have recently been reported to have activating effects on hPADI4 at significantly lower Ca^2+^ concentrations [34,42]. Without knowledge of the hPADI6 binding partners, this cannot be tested. As binding partners are identified, it is of upmost importance to characterise their capacity to activate hPADI6. Immediate candidates for this are members of the SCMC and other components of the CPLs.

## Methods

### Cloning and preparation of pcDNA3.1-Strep-Strep-TEV-hPADI6 recombinant DNA

Primers were designed using SnapGene software (www.snapgene.com) for In-Fusion cloning. A full list of oligonucleotides used in this work can be found in S1 Table. The hPADI6 CDS (DNASU, #HsCD00297377) was amplified by PCR. The pcDNA3.1-Strep-Strep-TEV vector (donated by the McDonald lab at the Francis Crick Institute) was also linearised by PCR. Amplified fragment inserts and linearised vectors were resolved by gel electrophoresis and extracted and purified from the gel using the PureLink™ Quick Gel Extraction Kit (Invitrogen, #K210012) following the manufacturers protocol. Fragment (∼50 ng) and linearised vector (∼50 ng) were combined with 2 µL 5X In-Fusion HD Enzyme (Takara Bio, #938910) and mixture made up to 10 µL with Nuclease Free Water (Fisher Bioreagents, #10336503). The mixture was incubated for 15 mins at 50 °C and 5 µL transformed into 50 µL a *E. Coli* NEB® 5-alpha aliquot (NEB®, #C2987H). Plasmids were sanger sequenced by GeneWiz (Azenta Life Sciences) to confirm successful gene insertion and MaxiPrepped using the ZymoPURE II™ Plasmid Maxiprep kit (Zymo, #D4202) before transfection into Expi293™ cells.

### Plasmid mutagenesis

Mutagenesis primers were designed to be ∼35 bp in length, with a melting temperature >78 °C with a GC content of ∼40% terminating in C or G. A full list of oligonucleotides used in this work can be found in S1 Table. Mutagenesis was performed following the Agilent QuickChange Protocol. In brief, PCR mixes were made up as follows in a total volume of 50 µL: 1X Pfu reaction buffer (Agilent, #600250), 0.2 mM dNTP (NEB®, #N0447L), 2.5 ng/µL forward primer, 2.5 ng/µL reverse primer, 25 ng plasmid and 0.5 µL Pfu Polymerase (Agilent, #600250). The PCR program was as follows: hot start at 98 °C for 30 sec, denaturation at 98 °C for 30 sec, annealing at 50 °C for 1 min, extension at 72 °C for 1 min per kb of plasmid. Denaturation, annealing and extension steps repeated for 15 cycles. After PCR amplification, 1 µL Dnp1 restriction enzyme (NEB, #R0176S) was added to the PCR mixture and the sample incubated at 37 °C for 1 h. After 1 h, 5 µL were transformed into a E. Coli NEB® 5-alpha aliquot (NEB®, #C2987H). Plasmids were sanger sequenced by GeneWiz (Azenta Life Sciences) to confirm presence of desired mutation and MaxiPrepped using the ZymoPURE II™ Plasmid Maxiprep kit (Zymo, #D4202) before transfection into Expi293™ cells.

### Protein expression

Expi293™ cells (Gibco™, #A14527) were incubated with 150 rpm agitation, 8% CO_2_, at 37 °C and diluted twice weekly. For transfection, cells were counted using Vi-CELL XR counter (Beckmann Coulter®) and diluted to 2 x 10^6^ cells/mL in 400 mL pre-warmed Expi293™ Expression Medium (Gibco™, #A1435101). 24 hours later, cells were transfected as follows: 400 µg MaxiPrepped recombinant DNA (pcDNA3.1_Strep-Strep-TEV-hPADI6) was diluted in 20 mL Opti-MEM® (Gibco™, #11058021), 1.2 mL of 1 mg/mL PEI 25K™ (Polysciences, #23966-1) diluted in 18.8 mL Opti-MEM®. Both mixtures were gently mixed by inversion and incubated for 5 mins at RT before being combined and incubated for 20 mins at RT. After incubation, DNA-PEI mixture was titrated dropwise into 400 mL Expi293™ cell culture with constant agitation of the culture. Cultures were then for 3 days before harvesting by centrifugation at 2000 rpm for 15 mins at 4 °C. Cell pellets were then washed in cold 50 mL cold PBS cOmplete™ EDTA-free Protease Inhibitor Cocktail tablets (Merck, #73567200) and centrifuged at 2000 rpm for a further 10 minutes. The supernatant was discarded, cell pellet flash frozen in liquid nitrogen and stored at −80 °C.

### Protein purification

Frozen transfected Expi293™ cell pellets were re-suspended in purification buffer (50 mM Tris-HCl, pH 7.5, 200 mM NaCl, 1 mM TCEP) with cOmplete™ EDTA-free Protease Inhibitor Cocktail tablets (Merck, #73567200), 1 mg/mL Lysozyme (Sigma, #L6876-5G) and a micro-spatula full of DNAse (Roche, #10104159001). Cells were lysed on ice by sonication (3 secs on, 5 secs off, 30% intensity, 3 mins total on time). Crude lysate was centrifuged at 21,000 rpm for 45 mins on a JA-25.50 rotor (Beckman Coulter®) and filtered through 0.45 µm PVDF filters (Merck, #SLHV033RS) using a syringe. Cleared cell lysate was added to StrepTactin® XT 4Flow® resin (Iba, #2-5010-025) at a ratio of 2 mL packed beads per 500 mL Expi293 cell culture and incubated at 4 °C for 4 hours with gentle rotation. After 4 h, the unbound fraction was removed by gravity filtration and the resin was washed with purification buffer (5 times, double volume of resin). The resin was then re-suspended in twice its volume of purification buffer and TEV protease added at a ratio of 2 mg protease per 500 mL Expi293 cell culture. The cleavage was then incubated at 4 °C overnight with gentle rocking. The cleaved fraction was then removed by gravity filtration and the resin washed with purification buffer (3 times, double volume of resin). The cleaved and wash fractions were combined and concentrated to 2 mL using a Vivaspin® 20, 30000 MWCO ultrafiltration centrifugal concentrator (Sartorius, #VS2022). The concentrated protein samples were then purified by size exclusion chromatography using a HiLoadTM 16/600 SuperdexTM 200 pg Gel Filtration Column (Cytivia, # 28989335), in purification buffer on an ÄKTApure™ system (Cytivia). Fractions were characterised by SDS-PAGE and pooled. Pooled fractions were concentrated again until no further decrease in volume was observed using a Vivaspin® 20, 30000 MWCO ultrafiltration centrifugal concentrator and stored at −80 °C.

hPADI6 variants were purified following the same protocol as the wild type protein excluding the final SEC step and were pooled and stored after cleavage from StrepTactin® XT 4Flow® resin.

### Intact-MS

Denatured proteins were injected onto a C4 BEH 1.7µm, 1 mm x 100 mm UPLC column (Waters™, #186005590) using an Acquity I class LC (Waters™). Proteins were eluted with a 15 min gradient of acetonitrile (2% v/v to 80% v/v) in 0.1% v/v formic acid using a flow rate of 50 µl/min. The analytical column outlet was directly interfaced via an electrospray ionisation source, with a time-of-flight BioAccord mass spectrometer (Waters™). Data was acquired over a *m/z* range of 300 – 8000, in positive ion mode with a cone voltage of 40v. Scans were summed together manually and deconvoluted using MaxEnt1 (Masslynx, Waters™).

### COLour DEveloping Reagent (COLDER) assay

PADI activity was assessed using the Colour Development Reagent (COLDER) assay [20]. Reactions were carried out in 96-well plates (Thermo Scientific™, #260895). hPADI6 (final conc. = 500 nM) or hPADI4 (final conc. = 50 nM) in COLDER buffer (50 mM HEPES pH 7.5, 150 mM NaCl, 10 mM CaCl_2_, 2 mM DTT) were combined with PADI substrate (BAA (fluorochem., #F242800-1G), BAME (fluorochem., #F329642-1G), or BAEE (Merck, #B4500); final conc. = 10 mM) and incubated for 1 h at RT. A negative control with no protein was included for each experiment. After 1 h, reactions were quenched with 50 mM EDTA and citrulline level in the mixtures were determined as follows. 1 volume of 80 mM Diacetyl monoxime/2,3-butanedione monoxime (Sigma, #31550-25G), 2 mM Thiosemicarbazide (Thermo Scientific Chemicals, #138901000), and 3 volumes of 3 M H_3_PO_4_, 6 M H_2_SO_4_ and 2 mM NH_4_Fe(SO_4_)_2_, was added to each well and the plate heated for 15 mins at 95 °C. After 15 mins, plates were cooled to RT and absorbance measured at a wavelength of 540 nm using a CLARIOstar Plus microplate reader (BMG Labtech). Absorbance bar charts were plotted using Prism (GraphPad Software). All experiments were conducted as two independent replicates of three technical replicates. Statistical significance was calculated using Prism using an unpaired parametric students t-test.

### Mass photometry

Proteins were diluted to an approximate concentration of 20 nM in purification buffer (50 mM Tris-HCl, pH 7.5, 200 mM NaCl, 1 mM TCEP). 2 µL of diluted protein was then added to 18 µL of PBS in the well of a gasket on a TwoMP instrument (Refeyn) and events recorded over the course of 1 min using the AquireMP software (Refeyn). Molecule sizes were then calculated using the AnalyseMP software (Refeyn) using BSA (66 kDa, Thermo Scientific™, #23209), ADH (150 kDa, Sigma-Aldrich, #A7011) and Urease (90/272/544 kDa, Sigma-Aldrich, #94280) as standards. Mass normalised event histograms were then produced using Prism (GraphPad Software) with bin sizes of 5 kDa.

### Nano differential scanning fluorimetry

Proteins were diluted to an approximate concentration of 0.2 mg/mL in purification buffer (50 mM Tris-HCl, pH 7.5, 200 mM NaCl, 1 mM TCEP) with either 10 mM EDTA or 10 mM CaCl_2_. Melting experiments were performed using Prometheus™ NT.48 High Sensitivity Capillaries (NanoTemper, #PR-C006-200) on a Prometheus (NanoTemper) instrument with a 20 to 95 °C melt at 1 °C per minute. Melting temperatures were extracted as the first derivative peak in the melting curve calculated using the Panta Analysis software (NanoTemper). All experiments were conducted as two independent replicates of three technical replicates. Statistical significance was calculated using Prism using an unpaired parametric students t-test.

### Crystallisation, data collection, and structure determination of hPADI6

hPADI6 was concentrated to 5 mg/ml and crystallised at 20°C using sitting-drop vapour diffusion. Sitting drops of 600 nL consisted of a 6:5:1 (vol:vol:vol) mixture of protein, well solution and seed stocks from a previous optimisation tray. Well solutions consisted of 20.9% w/v PEG3350, 0.2 M KSCN, 0.1 M Bis-Tris propane, pH 6.5. Crystals appeared within a few days and reached their maximum size after 21 days. Crystals were cryoprotected in perfluoropolyether cryo-oil (Hampton research, #HR2-814). X-ray data were collected at 100 K at beamlines I03 (mx25587-64) of the Diamond Light Source Synchrotron (Oxford, UK). Data collection and refinement statistics are summarized in Table 1. Data sets were indexed, scaled and merged with xia2 [43]. Molecular replacement used the atomic coordinates of an AlphaFold 2.0 model of PADI6 dimer. Refinement used Phenix [44]. Model building used COOT [45] with validation by PROCHECK [46]. Figures were prepared using the PyMOL Molecular Graphics System, Version 2.0 (Schrödinger, LLC). Atomic coordinates and crystallographic structure factors have been deposited in the Protein Data Bank under accession code PDB: 9FMN.

### Calculation of structural damage and variant effect prediction scores

FoldX 5.0 [47] was used to calculate ΔΔG values for all possible missense variants using both the full structure of the hPADI6 dimer and the monomeric subunit considered in isolation. The ‘RepairPDB’ function was run before modelling, and 10 replicates were used for ΔΔG calculation. Absolute ΔΔG values were then converted to the rank normalised ΔΔG_rank_ metric to improve comparability and interpretability of variants [38]. Variant effect predictor scores were taken from a recent study [39], and rank normalised in the same manner as the ΔΔG values to convert them to a 0-1 scale. For all patients and putatively unaffected controls, a biallelic score could then be calculated in the same way, by averaging the 0-1 score for each allele, considering wild type as a value of 0, and null variants as a value of 1.

Two “putatively unaffected” sets were considered for comparison of the biallelic scores. First, gnomAD v2.1 [48] was used, considering all distinct protein-coding variants as separate cases, regardless of allele frequency. For homozygous variants, the biallelic score was calculated from two copies of the same variant, while for heterozygous variants, the other allele was considered to be wild type, given the absence of information about phasing. Second, homozygous missense variants from primAD [49] were used, based on sequencing of non-human primate species. primAD has the advantage of containing far more homozygous variants than gnomAD, facilitating the calculation of individual-level biallelic scores.

### PADI6 multi-species sequence alignment and logo plot

The aligned PADI6 coding sequences from all 90 species recorded with a *PADI6* orthologue gene were extracted from Ensembl (Release 111)[50] on 25/04/2024 and exported into JalView [51]. For an aligned list of Ensembl protein sequences used see S1 File. 10 sequences were excluded due to missing regions in the sequence or multiple stop codons within the coding sequence, leaving 80 sequences. Aligned sequences corresponding to regions of interest in hPADI6 were exported in FASTA format and logo plots produced using WebLogo.berkeley.edu [36,37].

## Acknowledgements

The authors thank the members of the Structural Biology Science Technology Platform at The Francis Crick Institute for assistance, in particular Roger George, and Chloë Roustan for their aid with mass photometry and mammalian cell culture respectively. This work was supported by the Francis Crick Institute which receives its core funding from Cancer Research UK (CC2030), the UK Medical Research Council (CC2030), and the Wellcome Trust (CC2030). JPCW was supported by a Crick-Imperial College London studentship funded by the Department of Chemistry at Imperial College London and the Francis Crick Institute. This project was also supported by funding to JAM from the European Research Council (ERC) under the European Union’s Horizon 2020 research and innovation programme (grant agreement No. 101001169).

## Author Contributions

**Conceptualisation:** Louise J. Walport, Jack P. C. Williams, M. Teresa Bertran, Joseph A. Marsh

**Funding acquisition:** Louise J. Walport, Joseph A. Marsh

**Investigation:** Jack P. C. Williams, Stephane Mouilleron, Rolando Hernandez Trapero

**Project Administration:** Louise J. Walport, Jack P. C. Williams

**Supervision:** Louise J. Walport, Joseph A. Marsh, M. Teresa Bertran

**Visualization:** Jack P. C. Williams, Stephane Mouilleron

**Writing – Original Draft Preparation:** Jack P. C. Williams

**Writing – Review & Editing:** Jack P. C. Williams, Louise J. Walport, Stephane Mouilleron, Rolando Hernandez Trapero, Joseph A. Marsh

## References

1. Qian J, Nguyen NMP, Rezaei M, Huang B, Tao Y, Zhang X, et al. Biallelic PADI6 variants linking infertility, miscarriages, and hydatidiform moles. Eur J Hum Genet. 2018;26(7):1007–13.

2. Dong J, Fu J, Yan Z, Li L, Qiu Y, Zeng Y, et al. Novel biallelic mutations in PADI6 in patients with early embryonic arrest. J Hum Genet. 2022;(September):1–9.

3. Maddirevula S, Coskun S, Awartani K, Alsaif H, Abdulwahab FM, Alkuraya FS. The human knockout phenotype of PADI6 is female sterility caused by cleavage failure of their fertilized eggs. Clin Genet. 2017;91(2):344–5.

4. Zhang T, Liu P, Yao G, Zhang X, Cao C. A complex heterozygous mutation in PADI6 causes early embryo arrest: A case report. Front Genet. 2023 Jan 10;13:1104085.

5. Xu Y, Shi Y, Fu J, Yu M, Feng R, Sang Q, et al. Mutations in PADI6 Cause Female Infertility Characterized by Early Embryonic Arrest. Am J Hum Genet. 2016;99(3):744–52.

6. Wang X, Song D, Mykytenko D, Kuang Y, Lv Q, Li B, et al. Novel mutations in genes encoding subcortical maternal complex proteins may cause human embryonic developmental arrest. Reprod Biomed Online. 2018;36(6):698–704.

7. Rezaei M, Suresh B, Bereke E, Hadipour Z, Aguinaga M, Qian J, et al. Novel pathogenic variants in NLRP7, NLRP5, and PADI6 in patients with recurrent hydatidiform moles and reproductive failure. Clin Genet. 2021;99(6):823–8.

8. Xu Y, Wang R, Pang Z, Wei Z, Sun L, Li S, et al. Novel Homozygous PADI6 Variants in Infertile Females with Early Embryonic Arrest. Front Cell Dev Biol. 2022;10(April):1–8.

9. Zheng W, Chen L, Dai J, Dai C, Guo J, Lu C, et al. New biallelic mutations in PADI6 cause recurrent preimplantation embryonic arrest characterized by direct cleavage. J Assist Reprod Genet. 2020;37(1):205– 12.

10. Liu J, Tan Z, He J, Jin T, Han Y, Hu L, et al. Two novel mutations in PADI6 and TLE6 genes cause female infertility due to arrest in embryonic development. J Assist Reprod Genet. 2021;38(6):1551–9.

11. Eggermann T, Yapici E, Bliek J, Pereda A, Begemann M, Russo S, et al. Trans-acting genetic variants causing multilocus imprinting disturbance (MLID): common mechanisms and consequences. Clin Epigenetics. 2022;14(1):1–17.

12. Begemann M, Rezwan FI, Beygo J, Docherty LE, Kolarova J, Schroeder C, et al. Maternal variants in NLRP and other maternal effect proteins are associated with multilocus imprinting disturbance in offspring. J Med Genet. 2018;55(7):497–504.

13. Cubellis MV, Pignata L, Verma A, Sparago A, Del Prete R, Monticelli M, et al. Loss-of-function maternal-effect mutations of PADI6 are associated with familial and sporadic Beckwith-Wiedemann syndrome with multi-locus imprinting disturbance. Clin Epigenetics. 2020;12(1):139.

14. Eggermann T, Kadgien G, Begemann M, Elbracht M. Biallelic PADI6 variants cause multilocus imprinting disturbances and miscarriages in the same family. Eur J Hum Genet. 2021;29(4):575–80.

15. Tannorella P, Calzari L, Daolio C, Mainini E, Vimercati A, Gentilini D, et al. Germline variants in genes of the subcortical maternal complex and Multilocus Imprinting Disturbance are associated with miscarriage / infertility or Beckwith – Wiedemann progeny. Clin Epigenetics. 2022;14(43):1–7.

16. Esposito G, Vitale AM, Leijten FPJ, Strik AM, Koonen-Reemst AMCB, Yurttas P, et al. Peptidylarginine deiminase (PAD) 6 is essential for oocyte cytoskeletal sheet formation and female fertility. Mol Cell Endocrinol. 2007;273(1–2):25–31.

17. Wright PW, Bolling LC, Calvert ME, Sarmento OF, Berkeley EV, Shea MC, et al. ePAD, an oocyte and early embryo-abundant peptidylarginine deiminase-like protein that localizes to egg cytoplasmic sheets. Dev Biol. 2003;256(1):74–89.

18. Zhang J, Dai J, Zhao E, Lin Y, Zeng L, Chen J, et al. cDNA cloning, gene organization and expression analysis of human peptidylarginine deiminase type VI. Acta Biochim Pol. 2004;51(4):1051–8.

19. Kan R, Jin M, Subramanian V, Causey CP, Thompson PR, Coonrod SA. Potential role for PADI-mediated histone citrullination in preimplantation development. BMC Dev Biol. 2012;12.

20. Knipp M, Vašák M. A colorimetric 96-well microtiter plate assay for the determination of enzymatically formed citrulline. Anal Biochem. 2000;286(2):257–64.

21. Taki H, Gomi T, Knuckley B, Thompson PR, Vugrek O, Hirata K, et al. Purification of enzymatically inactive peptidylarginine deiminase type 6 from mouse ovary that reveals hexameric structure different from other dimeric isoforms. Adv Biosci Biotechnol. 2011;02(04):304–10.

22. Ranaivoson FM, Bande R, Cardaun I, De Riso A, Gärtner A, Loke P, et al. Crystal structure of human peptidylarginine deiminase type VI (PAD6) provides insights into its inactivity. IUCrJ. 2024 May 1;11(3):395–404.

23. Saijo S, Nagai A, Kinjo S, Mashimo R, Akimoto M, Kizawa K, et al. Monomeric Form of Peptidylarginine Deiminase Type I Revealed by X-ray Crystallography and Small-Angle X-ray Scattering. J Mol Biol. 2016 Jul;428(15):3058–73.

24. Slade DJ, Fang P, Dreyton CJ, Zhang Y, Fuhrmann J, Rempel D, et al. Protein arginine deiminase 2 binds calcium in an ordered fashion: Implications for inhibitor design. ACS Chem Biol. 2015;10(4):1043–53.

25. Funabashi K, Sawata M, Nagai A, Akimoto M, Mashimo R, Takahara H, et al. Structures of human peptidylarginine deiminase type III provide insights into substrate recognition and inhibitor design. Arch Biochem Biophys. 2021 Sep;708:108911.

26. Arita K, Hashimoto H, Shimizu T, Nakashima K, Yamada M, Sato M. Structural basis for Ca2+-induced activation of human PAD4. Nat Struct Mol Biol. 2004;11(8):777–83.

27. Williams JPC, Walport LJ. PADI6: What we know about the elusive fifth member of the peptidyl arginine deiminase family. Philos Trans R Soc B Biol Sci. 2023 Nov 20;378(1890):20220242.

28. Yurttas P, Vitale AM, Fitzhenry RJ, Cohen-Gould L, Wu W, Gossen JA, et al. Role for PADI6 and the cytoplasmic lattices in ribosomal storage in oocytes and translational control in the early mouse embryo. Development. 2008;135(15):2627–36.

29. Kan R, Yurttas P, Kim B, Jin M, Wo L, Lee B, et al. Regulation of mouse oocyte microtubule and organelle dynamics by PADI6 and the cytoplasmic lattices. Dev Biol. 2011;350(2):311–22.

30. Liu X, Morency E, Li T, Qin H, Zhang X, Zhang X, et al. Role for PADI6 in securing the mRNA-MSY2 complex to the oocyte cytoplasmic lattices. Cell Cycle. 2017;16(4):360–6.

31. Yu XJ, Yi Z, Gao Z, Qin D, Zhai Y, Chen X, et al. The subcortical maternal complex controls symmetric division of mouse zygotes by regulating F-actin dynamics. Nat Commun. 2014;5:4887.

32. Jentoft IMA, Bäuerlein FJB, Welp LM, Cooper BH, Petrovic A, So C, et al. Mammalian oocytes store proteins for the early embryo on cytoplasmic lattices. Cell. 2023 Nov;S0092867423010851.

33. Liu YL, Chiang YH, Liu GY, Hung HC. Functional Role of Dimerization of Human Peptidylarginine Deiminase 4 (PAD4). Proost P, editor. PLoS ONE. 2011 Jun 22;6(6):e21314.

34. Zhou X, Kong S, Maker A, Remesh SG, Leung KK, Verba KA, et al. Antibody discovery identifies regulatory mechanisms of protein arginine deiminase 4. Nat Chem Biol. 2024 Jun;20(6):742–50.

35. Krissinel E, Henrick K. Inference of Macromolecular Assemblies from Crystalline State. J Mol Biol. 2007 Sep;372(3):774–97.

36. Crooks GE, Hon G, Chandonia JM, Brenner SE. WebLogo: A Sequence Logo Generator. Genome Res. 2004 Jun;14(6):1188–90.

37. Schneider TD, Stephens RM. Sequence logos: a new way to display consensus sequences. Nucleic Acids Res. 1990;18(20):6097–100.

38. Pino DC, Badonyi M, Semple CA, Marsh JA. Protein structural context of cancer mutations reveals molecular mechanisms and identifies novel candidate driver genes [Internet]. 2024 [cited 2024 Jun 4]. Available from: http://biorxiv.org/lookup/doi/10.1101/2024.03.21.586131

39. Livesey BJ, Marsh JA. Variant effect predictor correlation with functional assays is reflective of clinical classification performance [Internet]. 2024 [cited 2024 Jun 4]. Available from: http://biorxiv.org/lookup/doi/10.1101/2024.05.12.593741

40. Rose R, Rose M, Ottmann C. Identification and structural characterization of two 14-3-3 binding sites in the human peptidylarginine deiminase type VI. J Struct Biol. 2012;180(1):65–72.

41. Snow AJ, Puri P, Acker-Palmer A, Bouwmeester T, Vijayaraghavan S, Kline D. Phosphorylation-dependent interaction of tyrosine 3-monooxygenase/ tryptophan 5-monooxygenase activation protein (YWHA) with PADI6 following oocyte maturation in mice. Biol Reprod. 2008 Aug 1;79(2):337–47.

42. Bertran MT, Walmsley R, Cummings T, Valle Aramburu I, Benton DJ, Assalaarachchi J, et al. A cyclic peptide toolkit reveals mechanistic principles of peptidylarginine deiminase IV (PADI4) regulation [Internet]. 2023 [cited 2024 Jun 6]. Available from: http://biorxiv.org/lookup/doi/10.1101/2023.12.12.571217

43. Winter G, Lobley CMC, Prince SM. Decision making in *xia* 2. Acta Crystallogr D Biol Crystallogr. 2013 Jul 1;69(7):1260–73.

44. Adams PD, Afonine PV, Bunkóczi G, Chen VB, Davis IW, Echols N, et al. *PHENIX* : a comprehensive Python-based system for macromolecular structure solution. Acta Crystallogr D Biol Crystallogr. 2010 Feb 1;66(2):213–21.

45. Emsley P, Lohkamp B, Scott WG, Cowtan K. Features and development of *Coot*. Acta Crystallogr D Biol Crystallogr. 2010 Apr 1;66(4):486–501.

46. Vaguine AA, Richelle J, Wodak SJ. *SFCHECK* : a unified set of procedures for evaluating the quality of macromolecular structure-factor data and their agreement with the atomic model. Acta Crystallogr D Biol Crystallogr. 1999 Jan 1;55(1):191–205.

47. Delgado J, Radusky LG, Cianferoni D, Serrano L. FoldX 5.0: working with RNA, small molecules and a new graphical interface. Valencia A, editor. Bioinformatics. 2019 Oct 15;35(20):4168–9.

48. Karczewski KJ, Francioli LC, Tiao G, Cummings BB, Alföldi J, Wang Q, et al. The mutational constraint spectrum quantified from variation in 141,456 humans. Nature. 2020 May 28;581(7809):434–43.

49. Gao H, Hamp T, Ede J, Schraiber JG, McRae J, Singer-Berk M, et al. The landscape of tolerated genetic variation in humans and primates. Science. 2023 Jun 2;380(6648):eabn8153.

50. Harrison PW, Amode MR, Austine-Orimoloye O, Azov AG, Barba M, Barnes I, et al. Ensembl 2024. Nucleic Acids Res. 2024 Jan 5;52(D1):D891–9.

51. Waterhouse AM, Procter JB, Martin DMA, Clamp M, Barton GJ. Jalview Version 2—a multiple sequence alignment editor and analysis workbench. Bioinformatics. 2009 May 1;25(9):1189–91.

52. Sievers F, Wilm A, Dineen D, Gibson TJ, Karplus K, Li W, et al. Fast, scalable generation of high-quality protein multiple sequence alignments using Clustal Omega. Mol Syst Biol. 2011 Jan;7(1):539.

