## Supplementary Information for "Structural Insight into the Function of Human Peptidyl Arginine Deiminase 6"

### Supporting information

**S1 File. PADI6 protein sequences used for homology analysis**

FASTA alignment of 80 PADI6 protein sequences used to generate sequence homology logo plots. Ensembl gene identifier and FASTA sequence provided for each gene after sequence alignment with sequence gaps shown by dashes.

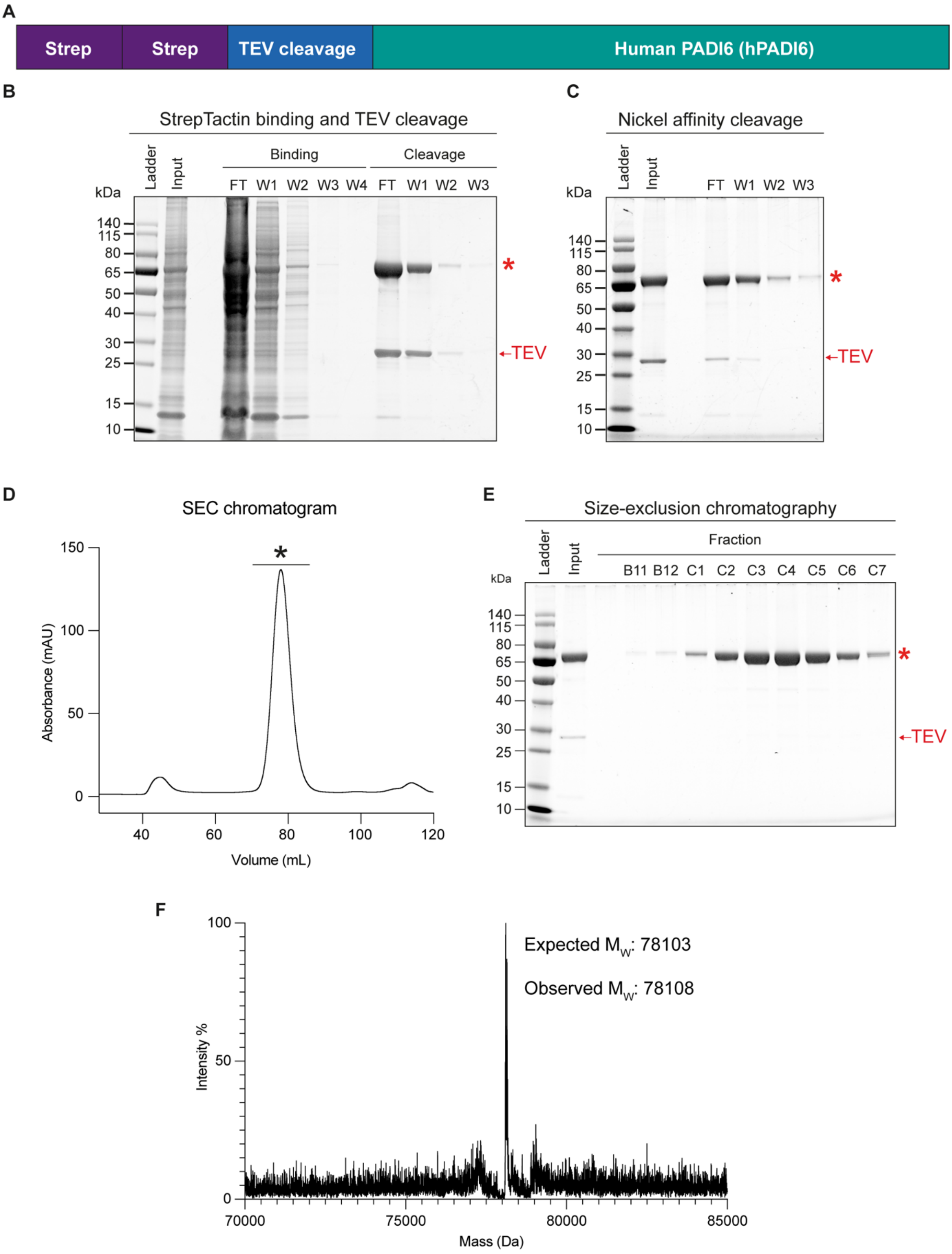


**S1 Fig. Recombinant expression of hPADI6**

(A) Expression construct schematic of hPADI6 (pcDNA3.1_Strep-Strep-TEV-hPADI6) with two Strep tags (sequence = WSHPQFEK), a TEV cleavage site (sequence = ENLYFQS) and the hPADI6 gene. (B) SDS-PAGE gel after binding of Expi293 lysate to StrepTactin resin, followed by on-resin cleavage of hPADI6 with

TEV protease. FT = Flow through, W1-W3/4 = washes. Red star indicates band at approximate hPADI6 $M_w$. (C) SDS-PAGE gel of sample before and after application to Nickel resin to remove TEV protease. FT = Flow through, W1-W3 = washes. (D) Chromatogram of SEC purification of hPADI6. Fractions taken for SDS-PAGE characterisation are labelled with a black star. SEC = size-exclusion chromatography. (E) SDS-PAGE gel of SEC purified fractions. Fractions C1 to C7 were combined and taken forward. (F) Intact-MS spectra of recombinant hPADI6.

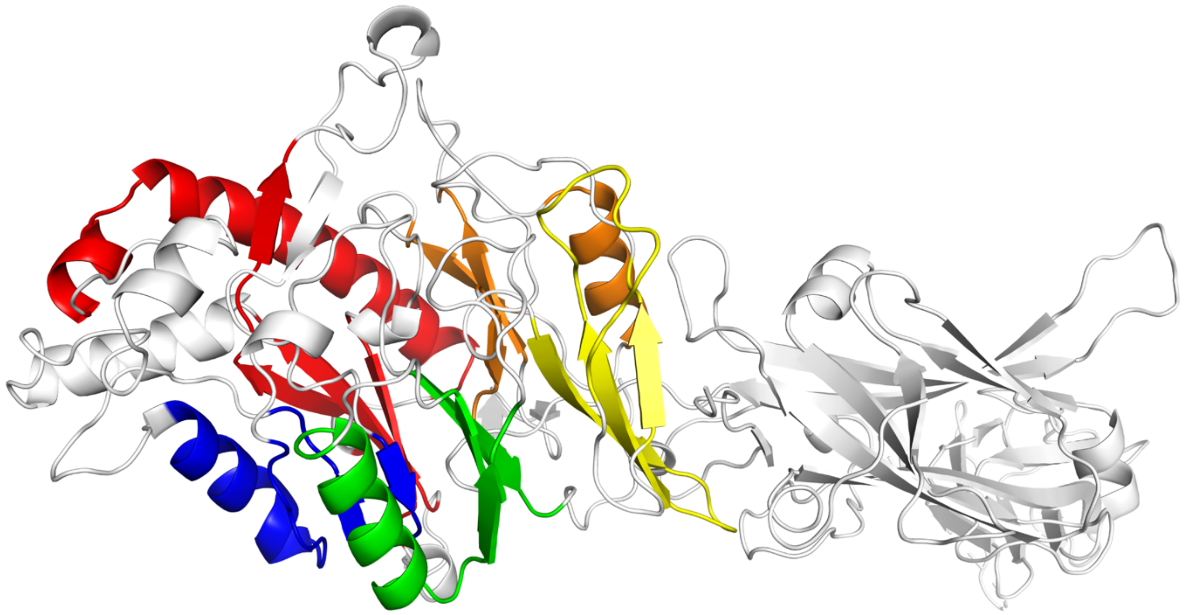

**S2 Fig. hPADI6 structure showing C-terminal ββαβ modules**

Ribbon representation of the hPADI6 structure, the five C-terminal ββαβ modules are coloured red, blue, green, yellow, and orange.

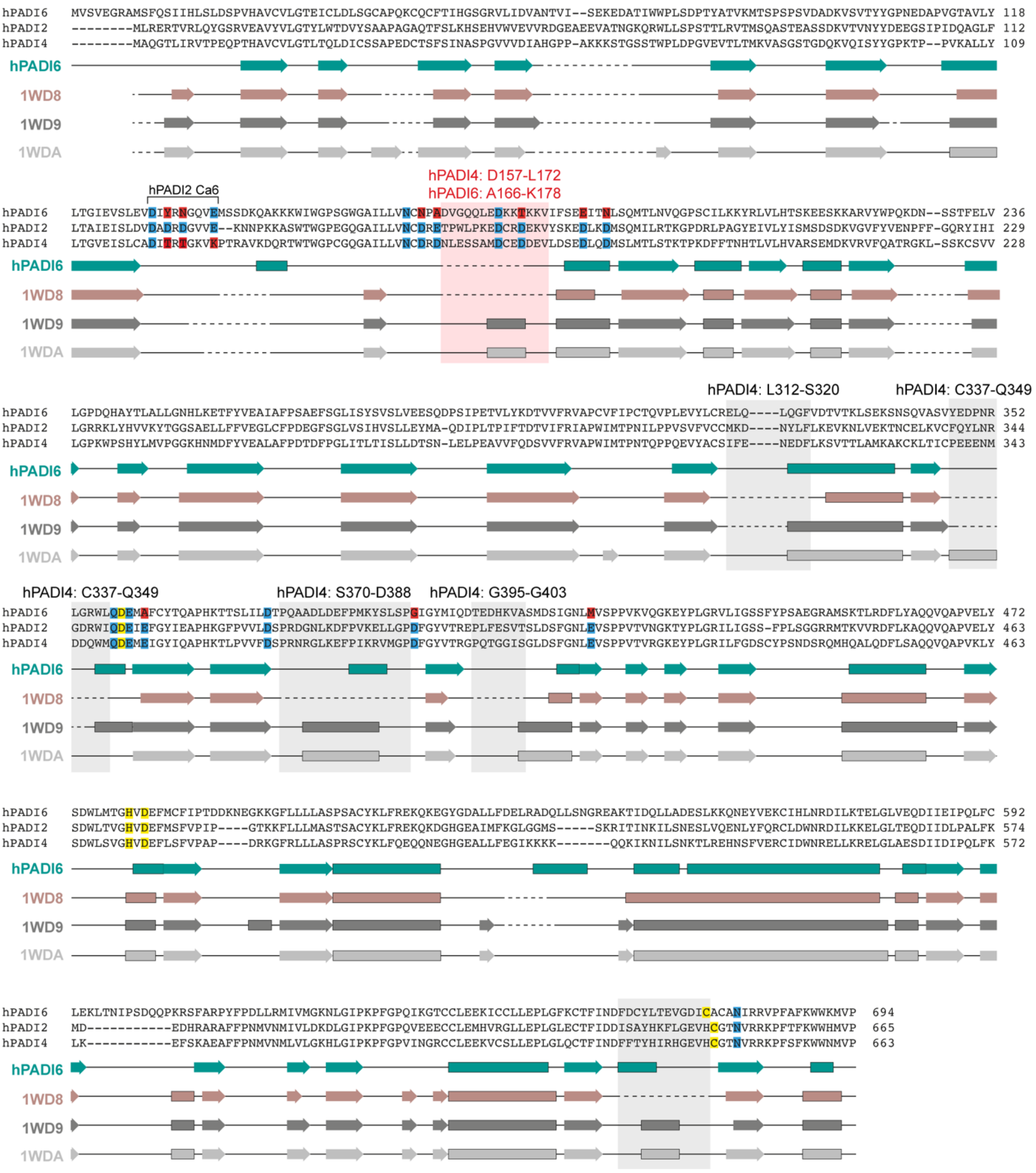


**S3 Fig. Multiple sequence alignment of hPADI2, hPADI4 and hPADI6**

Sequence alignment of hPADI6, hPADI2 and hPADI4 produced using Clustal Omega [52]. The secondary structure α-helices are displayed as bars and β-sheets as bars, for hPADI6, apo-hPADI4 (PDB: 1WD8), calcium bound hPADI4 (PDB: 1WD9), and calcium and substrate bound holo-hPADI4 (PDB: 1WDA) [26]. Disordered regions are shown by dashed lines. Conserved calcium binding residues are shaded blue, non-conserved calcium binding residues are shaded red. Catalytic tetrad amino acids are shaded yellow. The five loops that are disordered in apo-hPADI4, but structured in holo-hPADI4 and hPADI6 are highlighted by grey boxes with the corresponding loop residue numbers in hPADI4 highlighted. The one loop that is disordered in apo-hPADI4 and hPADI4, but structured in holo-hPADI4 is highlighted with a red box, with corresponding loop residue numbers in hPADI4 and hPADI6 highlighted.

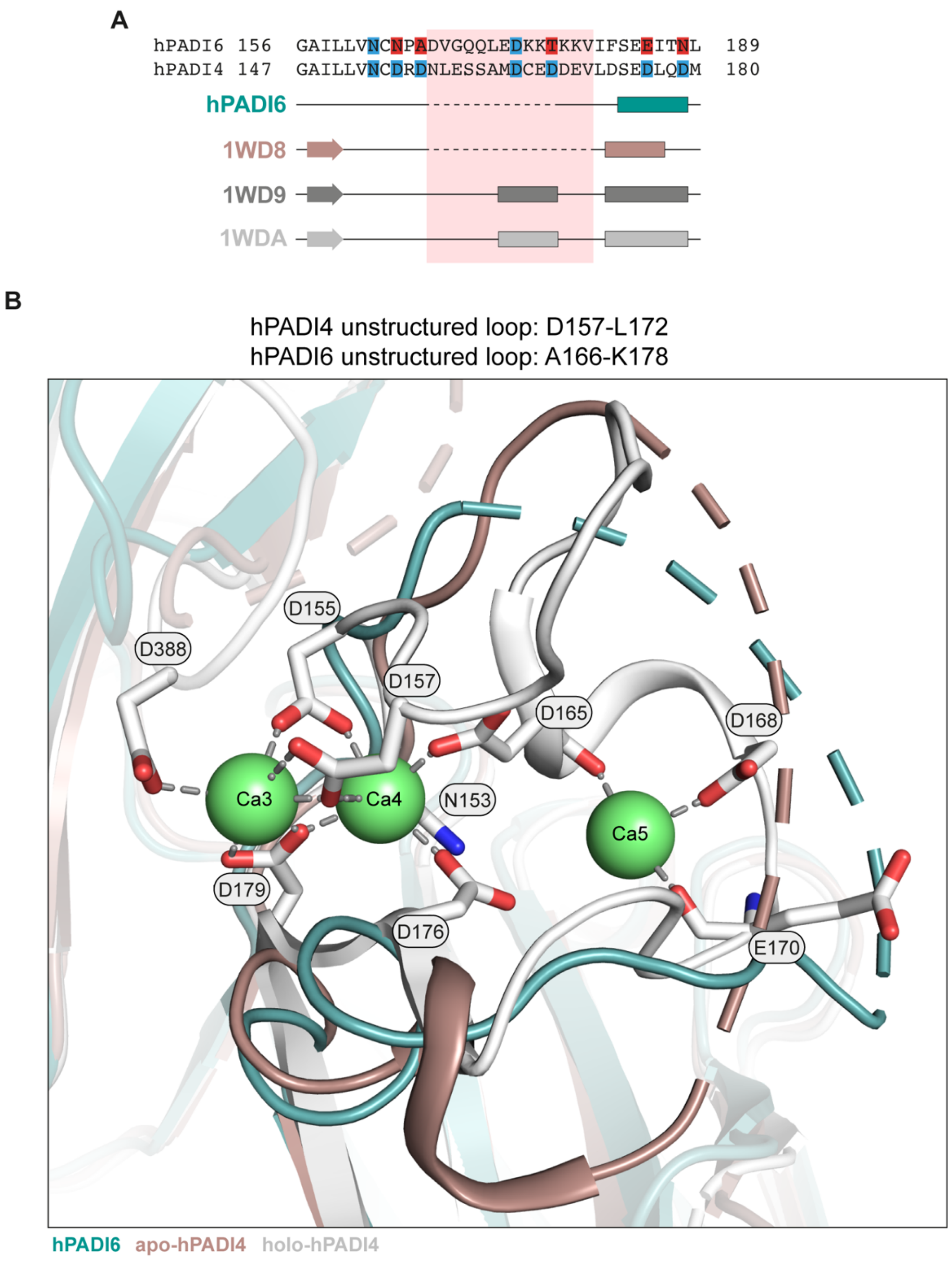


**S4 Fig. hPADI6 is disordered around the Ca3, 4, 5 binding site**

(A) Sequence alignment of hPADI6 and hPADI4 centred on the hPADI4 unstructured loop D157 to L172 and the hPADI6 unstructured loop A166 to K178 produced using Clustal Omega [52]. The secondary structure α-helices are displayed as bars and β-sheets as bars, for hPADI6, apo-hPADI4 (PDB: 1WD8), calcium bound hPADI4 (PDB: 1WD9), and calcium and substrate bound holo-hPADI4 (PDB: 1WDA) [26]. Disordered regions are shown by dashed lines. Conserved calcium binding residues are shaded blue, non-conserved calcium binding residues are shaded red. The unstructured loop D157 to L172 in the apo-hPADI4 and A166 to K178 in the hPADI6 structure is highlighted with a red box. (B) hPADI4 Ca3,4,5 site in the presence (holo-hPADI4, white, PDB:

1WDA) and absence (apo-hPADI4, brown, PDB: 1WD8) of $Ca^{2+}$, aligned with hPADI6 (teal). hPADI4 Ca3,4,5 coordinating residues are displayed in the holo-hPADI4 structure and interactions with $Ca^{2+}$ shown by grey dashed lines. Unstructured loop D157 to L172 in the apo-hPADI4 and A166 to K178 in the hPADI6 structure are shown by dashed lines.

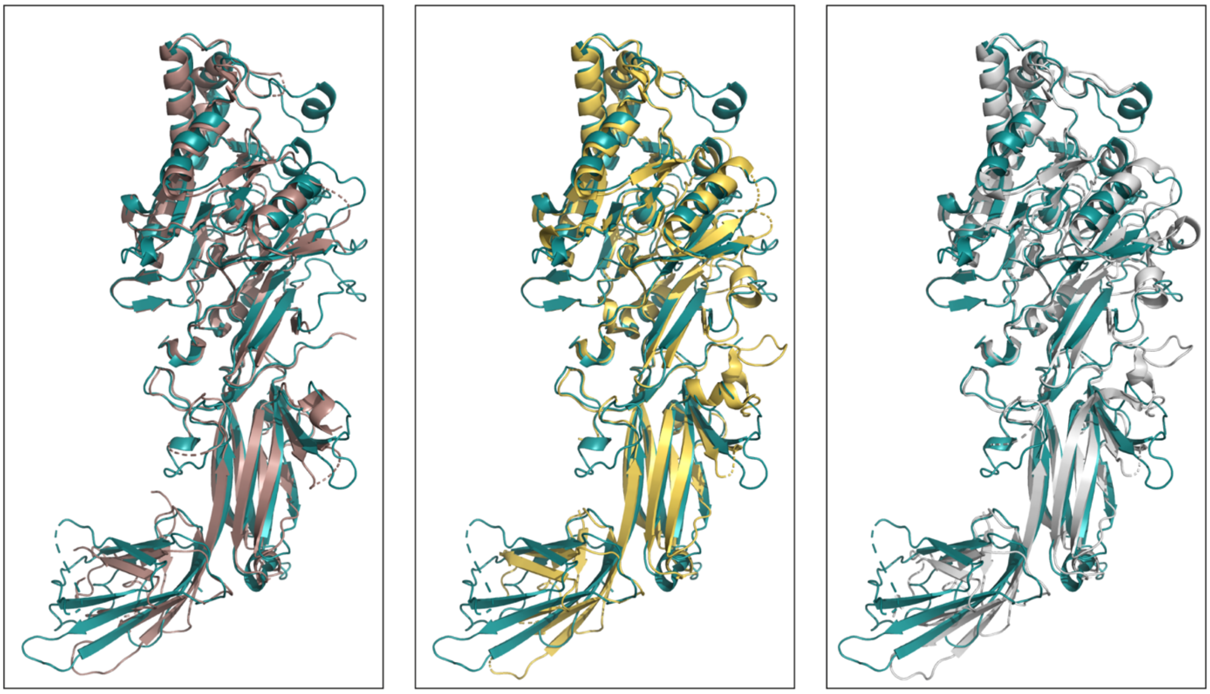

1WD8, WT 0 mM $CaCl_2$ (apo)
**PADI6 RMSD CA: 1.027 Å (416 CA)**

1WD9, C645A 5 mM $CaCl_2$
**PADI6 RMSD CA: 1.002 Å (430 CA)**

1WDA, C645A 5 mM $CaCl_2$ (holo)
**PADI6 RMSD CA: 1.060 Å (449 CA)**

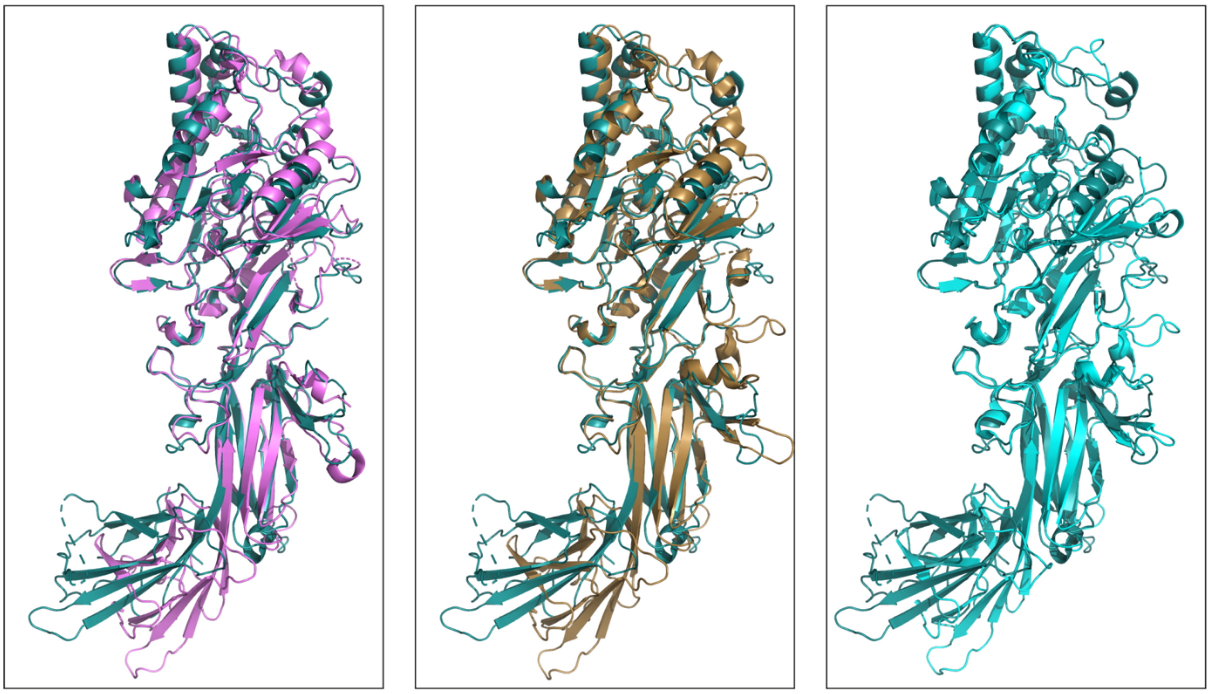

4N20, WT 0 mM $CaCl_2$ (apo)
**PADI6 RMSD CA: 1.134 Å (441 CA)**

4N2B, WT 10 mM $CaCl_2$
**PADI6 RMSD CA: 1.079 Å (434 CA)**

4N2C, F211/222A 10 mM $CaCl_2$ (holo)
**PADI6 RMSD CA: 1.018 Å (423 CA)**

**S5 Fig. Structural alignment of hPADI6 with hPADI2 and hPADI4 structures**

Top row: structural alignment of apo- (brown, PDB: 1WD8), calcium bound (yellow, PDB: 1WD9), and holo-hPADI4 (white, PDB: 1WDA) with hPADI6 (teal) [26]. Bottom row: structural alignment of apo-hPADI2 (pink, PDB: 4N20), calcium bound hPADI2 (bronze, PDB: 4N2B), and holo-hPADI2 (cyan, PDB: 4N2C) with hPADI6 (teal) [24]. Root mean square deviation (RMSD) values were calculated using only Cα positions.

RMSD values are reported in Å and the number of Cα used to calculated RMSD values noted for each alignment.

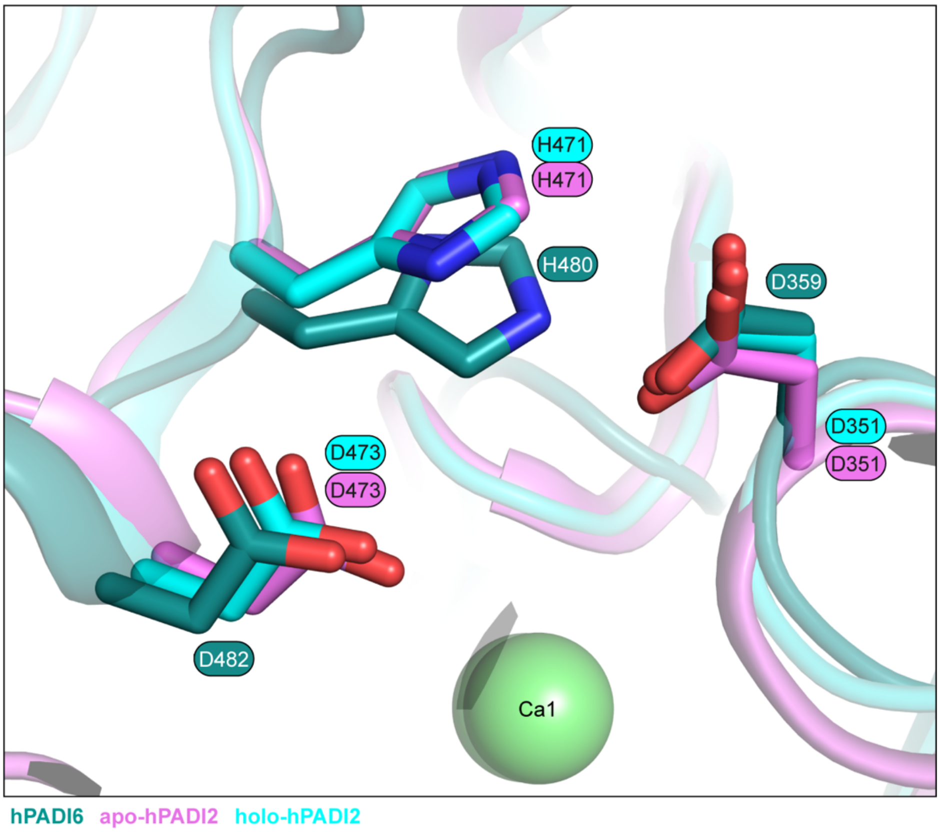


**S6 Fig. Structural alignment of the hPADI6 and hPADI2 catalytic residues**

Close up view of hPADI6 active site residues (teal), aligned with apo-hPADI2 (pink, PDB: 4N20), and holo-hPADI2 (pink, PDB: 4N2B) [24].

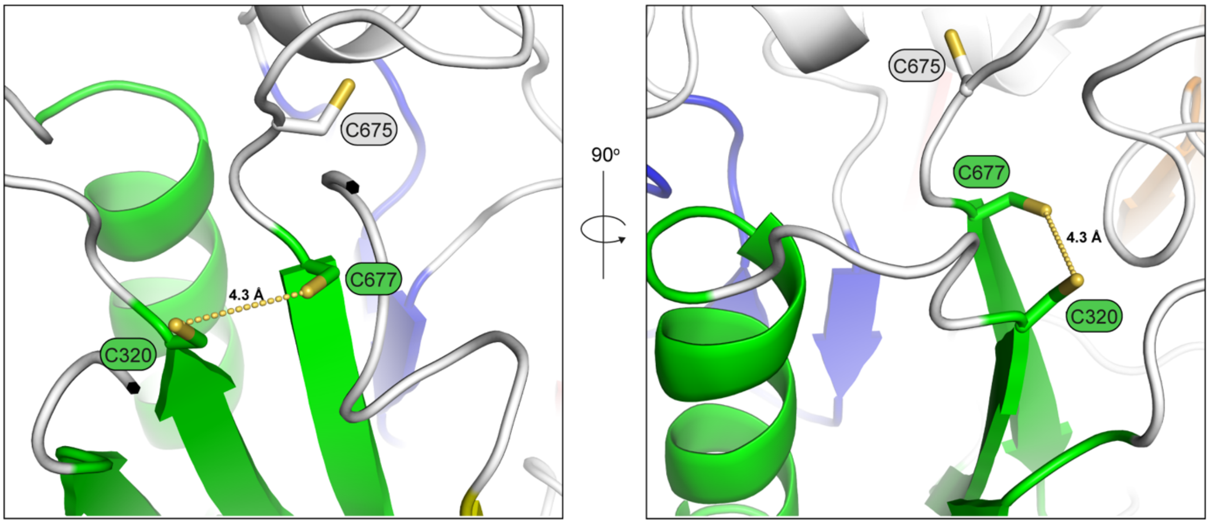


**S7 Fig. hPADI6 C777 and C320 position within ββαβ module**

Close up view of the positioning of hPADI6 C677 and C320 within a ββαβ module, separated by a distance of 4.3 Å.

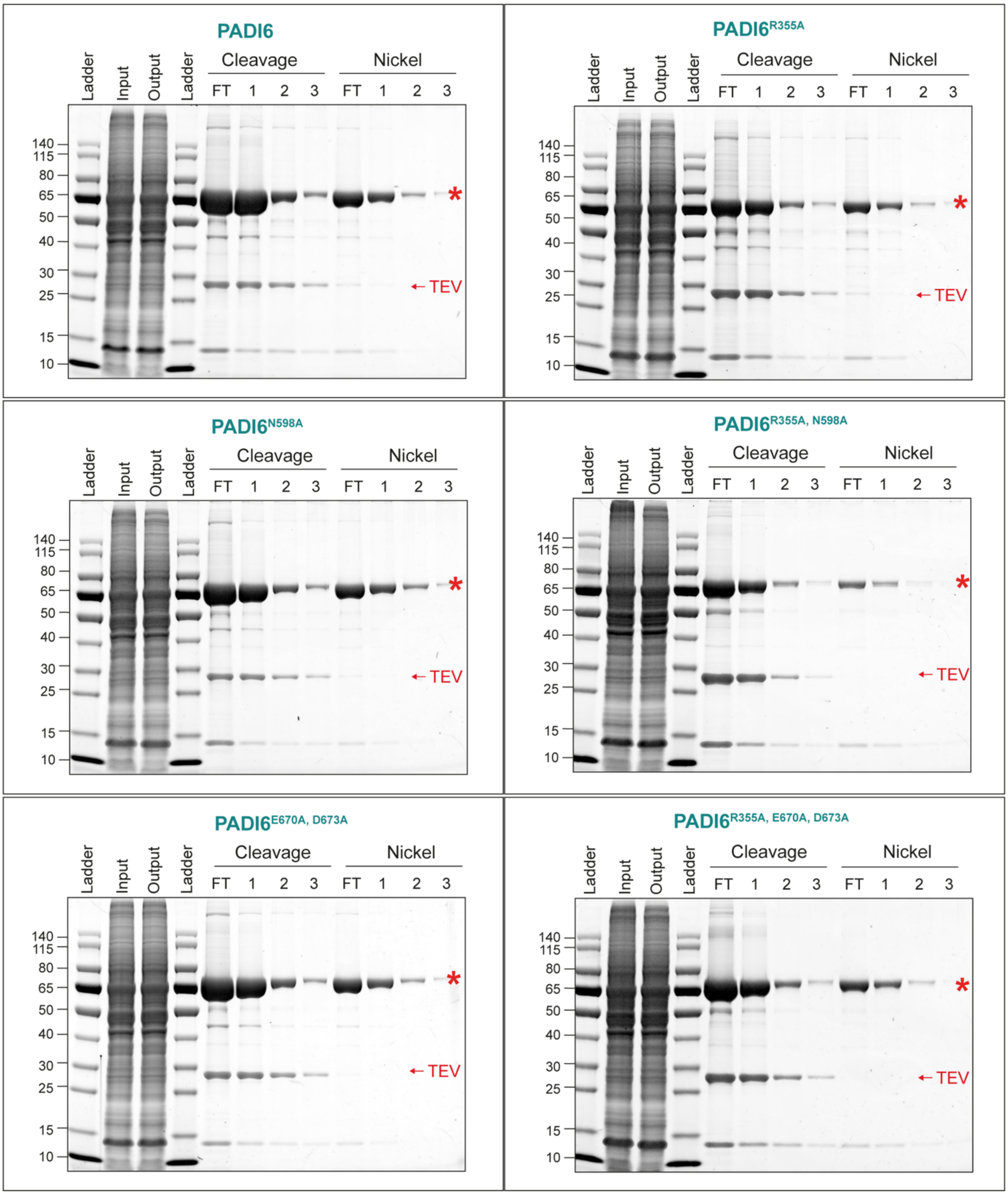


**S8 Fig. Expression of hPADI6 holding and blocking loop variants**

SDS-PAGE gel after binding of Expi293 lysate to StrepTactin resin, followed by on-resin cleavage of hPADI6 variants with TEV protease. FT = Flow through, W1-W3/4 = washes. Red star indicates band at approximate hPADI6 $M_w$.

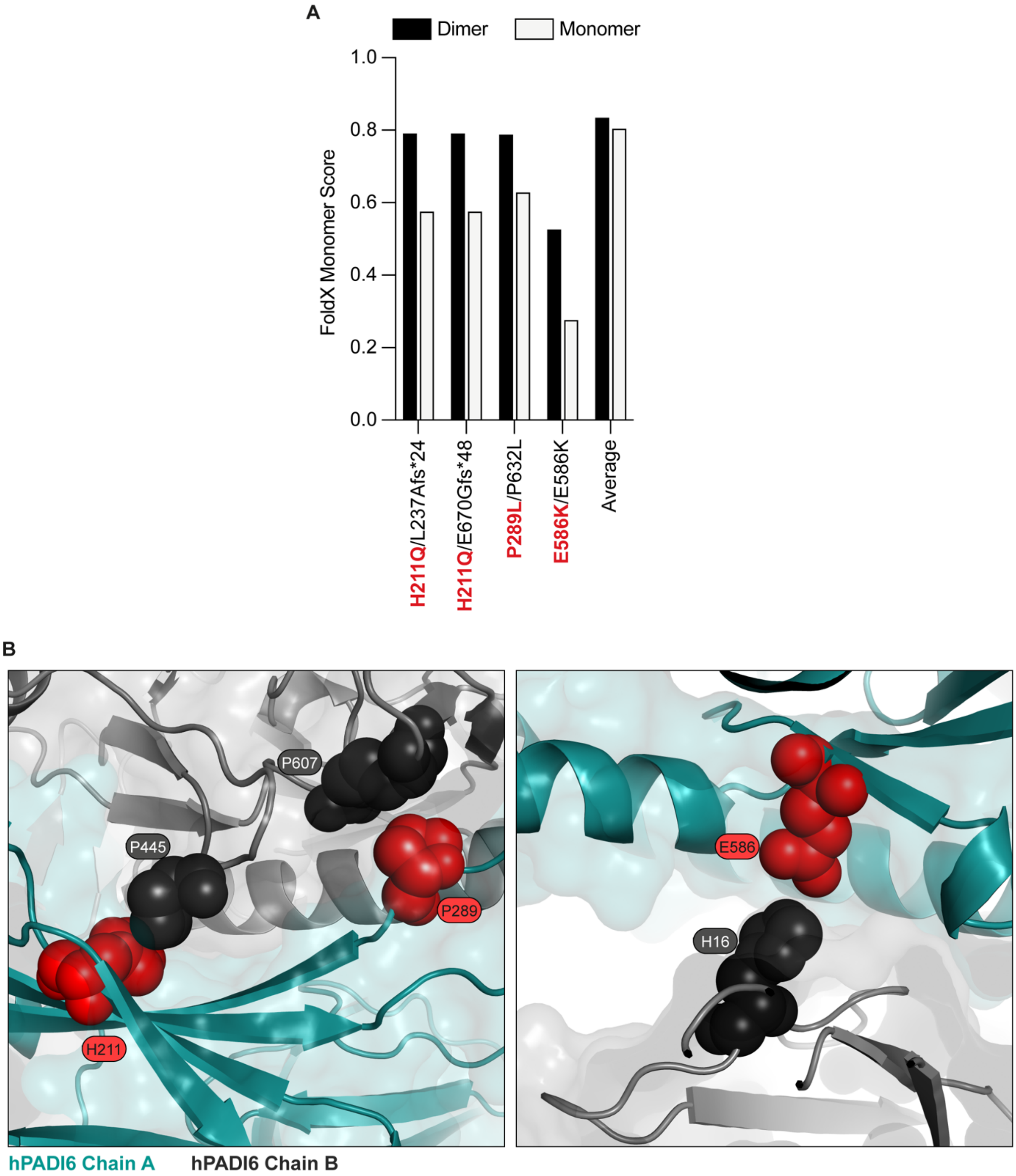


**S9 Fig. Three pathogenic *Padi6* variants possibly disrupt dimerisation**

(A) Biallelic structural damage scores, calculated from the full dimer, or monomer structure, for women with variants associated with infertility that result in a substitution at the dimer interface: H211Q, P289L, and E586K, as well as an average of the dimer and monomer structure damage scores for all pathogenic biallelic structural damage scores. (B) Close-up view of H211, P289 and E586 on chain A, and proximal amino acids on chain B, P445, P607 and H16 respectively.

**S1 Table. Oligonucleotides used in this work**

Oligonucleotides used for InFusion cloning and mutagenesis. Bb = Backbone, I = Insert, F = Forward, R = Reverse. All oligonucleotides were synthesised by Integrated DNA Technologies IDT.

| **InFusion cloning** | | |
|---|---|---|
| Plasmid | Primer | Sequence |
| pcDNA3.1_ Strep-Strep-TEV-hPADI6 | Bb-F | TAATAATCTAGAGGGCCCGTTTAAACCC |
| | Bb-R | GCTGCCGCTACCTGACT |
| | I-F | TCAGGTAGCGGCAGCATGGTCAGCGTGGAG |
| | I-R | CCCTCTAGATTATTATTAAGGTACCATCTTCCACCATTTGAAG |
| **Mutagenesis** | | |
| Variant | Primer | Sequence |
| hPADI6-N598A | F | CTGCTTGGAGAAGCTGACTGCGATCCCCTCTGACCAGCAGC |
| | R | GCTGCTGGTCAGAGGGGATCGCAGTCAGCTTCTCCAAGCAG |
| hPADI6-E760A, D673A | F | GACTTTGACTGTTACCTGACAGCGGTCGGAGCGATCTGTGCCTGTGCCA ACATC |
| | R | GATGTTGGCACAGGCACAGATCGCTCCGACCGCTGTCAGGTAACAGTC AAAGTC |
| hPADI6-R355A | F | GACCCCAACCGCCTGGGCGCGTGGCTCCAGGATGAGATG |
| | R | CATCTCATCCTGGAGCCACGCGCCCAGGCGGTTGGGGTC |
