## Supplementary material for "Structural Insight into the Function of Human Peptidyl Arginine Deiminase 6": S1 File

```
>ENSCGRG00001005116/1-684
--------MSFQSTVNLSLDSPTHAICVLGTEITLDISGCAPKNCESFTIRGSPRVLIHISSS--VITGKAD
AVVWQPMTQPTEALVRMVSPSPDVDEDKVLISFYSPDKEVPTATAVLYLTGIEISLEADIYRDGQLDMPSDK
QAKKRWVWGPNGWGAILLVNCNPAETEDL-DD-QSSDQEAPKDIQNLSQMTLTVEGPTSVFKNYMLILHTSK
EEASKARVYLSQKGTTCTYELVIGPGKPVHVLAP----FENRGKETFYIEAIEFPSASFAGLISFSISLVEK
SHDKSIPEMPLYKDTVMFRVAPYIFTPSTQMPLEVYLCREMQLQGFVDTVTRLSEKISVQVASVYEDPSRQG
RWLQDEMAFCYTQAPHKTVSLILDTPRVSKLEDFPMKYSLSPGVGYLTYKTEDHGVASLDSIGNMIVSPPVK
AQGKDYPLGRVLIGGSFYPSSEGRDMSKALRDFVCAQQVQAPVELFSDWLMTGHVDEFMCFVPVKDKNNDGK
GFRLLMASPSACYELFEQKQKEGFGDVTLFEDVRAEQLLSNGREARTISQILADQNLRKQNDYVEKCISLNR
TLLKKELGLMEKDIIAIPQLFCLELLTNVPSNQQTTKLFARPYFPDMLQMIVLDKNLGIPKPFGPQIKGTCC
LEEKVCNLLEPLGLKCTFIDDFDCYLTNMGDSCACVIIHRVPFAFKWWKMVP----
>ENSMAUG00000018048/1-642
----------------------------------------------SFAIRGSPRVLIHTSNS--VITGKTD
TVVWQPMTQPMDALVRMVSPSPTVDEDKVLVSYYCPENEVPKTTAVLYLTGIEISLEADIYRDGQLDMPSDK
QAKKKWVWGPNGWGAILLVNCNPV-TDEA-DD-EASDQEAPKEIQNMSQMTLTVDGPISVIKNYMMILHTSK
EEANKAKVYWPQKG---THEMVIGPGKPIHVLAP----FENGGKETFYIEAIEFPSASFTGLISFSISLVEE
SHDKLVPEMPLYKDTVMFRVAPYIFTPSTQPPLEVYLCRDLQLQGFVDTVTRLSEKINVQVASVYEDPSRQG
RWLQDEMAFCYTQAPHKTLSLVLDTPRVSKLDEFPMKYSLSPGVGYLTFETEDHGVASLDSIGNLIVSPPVK
AQGKDYPLGRVLIGGSFYPSNEGRDMSKVLRDFVCAQQVQAPVELFSDWLMTGRMDEFMCFVPVKDKNDDGK
GFRLLMASPSACYELFEEKQKEGFGDMTLFEDVGAEQLLSNGREARTISQILADQKLRKQNDYVEKCINLNR
TLLKKELGLTEKDIIAIPQLFCLELLTNVPSNQQTTKHYARPYFPNMLQMLVLDKHVGIPKPFGPQIKGTCC
LEEKVCDVLEPLGLKCTFIDDFDCYLTNMGDTCACAIVHRVPFAFKWWKMIP----
>ENSPEMG00000017319/1-692
EGQSSNLAMSFQSTLSLSLDSPTHAICVLGMEITLDISGCAPENCQSFTIRGSPRILIHISST--VITGKED
AVVWRPMNKPTEALVRMVSPSPTVDEDKVLVSYYCPDTEVPTATAVLYLTGVEVSLEADIYRDGQLDMPSDK
QAKKRWVWGPNGWGAILLVNCNPGERGES-EK-TRVWEPLYSEQENLSPMTLTVEGPQAVLKNYCLILHTSK
EEASKAKVYLLQKGTTSTYELVIGPGKPDHVLAP----FENRGKETFYIEAIEFPSANFTGLISFSVSLVEK
SYDQLIPEVPLYKDTVMFRVAPYIFTPSTQMPLELYLCRALQLQGFVDTVIRLSEKSNVQVASVYEDPNRQG
KWLQDEMAFCYTQAPHKTVSLIFDTPRVSNMEEFPIKYSLSPGVGYLTHHTEDHGVASLDSIGNLIVSPPVK
AQGKDYPLGRVLIGGSFYPSSEGRDMSKALRDFVYAQQVQAPVELFSDWLMTGHMDEFMCFVPVNDKYDEGK
GFRLLLASPSACYELFEQKQKEGYGDANLFDEVRPDQLISNGRESRTISQVLADENLRKQNDYVEKCISLNR
TLLKKELGLTDKDIIAIPQLFCLEMLTNIPSGQNFTKHFARPYFPDMVRMIVLGRNLGIPKPFGPLIKGTCC
LEERVCNLLEPLGLKCTFIDDFECYLTNMGDTCACVVIHRVPFAFKWWKMIP----
>ENSMSIG00000013004/1-682
--------MSFQNSLSLSLVNPTHALCMVGMEITLDISKCAPDKCKSFTIRGSPRILIHISSS--VIAGKED
TVVWRSMNHPTVALVRMVAPSPTVDEDKVLVSYFCPDQEVPTATAVLFLTGIEISLEADIYRDGQLDMPSDK
QAKKKWMWGMNGWGAILLVNCSPNAVGQP-DE-QS-FQEGPREIQNLSQMNVTVEGPTSILQNYQLILHTSE
EEAKKTRVYWSQRG-SSVYELVVGPNKPVYLLPT----FENRGKETFYVEATEFPSPSFSGLISLSLSLVEK
AHDECIPEIPLYKDTVMFRVAPYIFMPSTQMPLEVYLCRELQLQGFVDSVTKLSEKSKVQVVKVYEDPNRQS
KWLQDEMAFCYTQAPHKTVSLILDTPRVSKLEDFPMKYTLIPGFGYLIRQTEDHRVASLDSIGNLMVSPPVK
AQGKDYPLGRVLIGGSFYPSSEGRDMNKGLREFVYAQQVQAPVELFSDWLMTGHVDQFMCFVPTNDKNNDQK
DFRLLLASPSACFELFEQKQEEGYGNVTLFEDIGAEQLLSNGRESKTISQILADKSFREQNAYVEKCISLNR
TLLKTELGLEDKDIILIPQLFCLEQLTNVPSNQQSTKLFARPYFPDMLQIIVLGKNLGIPKPFGPKIKGTCC
LEEKVCGLLEPLGLKCTFIDDFDCYLANIGDVCASAIINRVPFAFKWWKMTP----
>ENSMUSG00000040935/1-682
--------MSFQNSLSLSLVNPTHALCMVGMEITLDISKCAPDKCKSFTIRGSPRILIHISSS--VIAGKED
TVVWRSMNHPTVALVRMVAPSPTVDEDKVLVSYFCPDQEVPTATAVLFLTGIEISLEADIYRDGQLDMPSDK
QAKKKWMWGMNGWGAILLVNCSPNAVGQP-DE-QS-FQEGPREIQNLSQMNVTVEGPTSILQNYQLILHTSE
EEAKKTRVYWSQRG-SSAYELVVGPNKPVYLLPT----FENRRKEAFYVEATEFPSPSFSGLISLSLSLVEK
AHDECIPEIPLYKDTVMFRVAPYIFMPSTQMPLEVYLCRELQLQGFVDSVTKLSEKSKVQVVKVYEDPNRQS
KWLQDEMAFCYTQAPHKTVSLILDTPRVSKLEDFPMKYTLTPGSGYLIRQTEDHRVASLDSIGNLMVSPPVK
AQGKDYPLGRVLIGGSFYPSSEGRDMNKGLREFVYAQQVQAPVELFSDWLMTGHMDQFMCFVPTNDKNNDQK
DFRLLLASPSACFELFEQKQKEGYGNVTLFEDIGAEQLLSNGRESKTISQILADKSFREQNTYVEKCISLNR
TLLKTELGLEDKDIILIPQLFCLEQLTNVPSNQQSTKLFARPYFPDMLQIIVLGKNLGIPKPFGPKINGTCC
LEEKVCGLLEPLGLKCTFIDDFDCYLANIGDVCASAIINRVPFAFKWWKMTP----
>MGP_SPRETEiJ_G0027703/1-682
--------MSFQNSLSLSLDNPTHALCMVGMEITLDISKCAPDKCKSFTIRGSPRILIHISSS--VIAGKED
TVVWRSMNHPTVALVRMVAPSPTVDEDKVLVSYFCPDQEVPTATAVLFLTGIEVSLEADIYRDGQLDMPSDK
QAKKKWMWGMNGWGAILLVNCSPNAVGQP-DE-QS-FQEGPREIQDLSPMSVTVEGPTSILQNYQLILHTSE
EEAKKTRVYWSQRG-SSAYELVVGPNKPVYLLPT----FENRRKETFYVEATEFPSPSFSGLISLSLSLVEK
AHDECIPEIPLYKDTVMFRVAPYIFMPSTQMPLEVYLCRELQLQGFVDSVTKLSKKSKVQVVKVYEDPNRQS
KWLQDEMAFCYTQAPHKTVSLILDTPRVSKLEDFPMKYTLIPGFGYLIRQTEDHRVASLDSIGNLMVSPPVK
AQGKDYPLGRVLIGGSFYPSSEGRDMNKGLREFVYAQQVQAPVELFSDWLMTGHMDQFMCFVPTNDKNNDQK
```

DFRLLLASPSACFELFEQKQKEGYGNVTLFEDIGAEQLLSNGRESKTISQILADKSFREQNAYVEKCISLNR
TLLKTELGLEDKDIILIPQLFCLEQLTNVPSNQQSTKLFARPYFPDMLQIIVLGKNLGIPKPFGPKIKGTCC
LEEKVCGLLEPLGLKCTFIDDFDCYLANIGDVCASAIINRVPFAFKWWKMIP----
>MGP_CAROLIEiJ_G0026723/1-682
--------MSFQSTLSLSLVNPTHALCMVGMEITLDISGCAPDKCKSFTIRGSPRILIHISSS--VIADKED
AIVWRPMNHPTEALVRMVAPSSTVDEDKVLVSYFCPDKDVPTATAVLFLTGIEVSLEADIYRDGQLDMPCDK
QAKKRWMWGLNGWGAILLVNCSPNAVGQP-DE-QS-FQEGPREIQNLSQMTLTVEGPTSILQNYQLILHTSE
EEAKRARVYWSQRG-SSAYELVVGPNKPVYLLPT----SENHGKEIFYVEATEFPSPSFSGLISFSLSLVEK
THNQCIPETPLYKDTVMFRVAPYIFMPSTQMPLEVYLCRELQLQGFVDSVTKLSEKSKVRVVTVYEDPNRQG
NWLQDEMAFCYTQAPHKTVSLILDTPRVFKLEDFPMKYSLSPGFGYLIRQTEDHRVASLDSIGNLMVSPPVK
AQGKDYPLGRVLIGGSFYPSSESRDMNKGLREFVYAQQVQAPVELFSDWLMTGHMDQFMCFVPTNDKNNDQK
DFRLLLASPSACFELFEQKQKEGYGNVTLFEDIGAEQLLSNGRESKTISQILADKSFREQNAYVEKCISLNR
TLLKTELGLEDKDIILIPQLFCLEQLTNVPSNLQSTKLFARPYFPDMLQIIVLGKNLGIPKPFGPQIKGTCC
LEERVCELLEPLGLKCTFIDDFDYYLANIGDVCASAIINRVPFAFKWWKMTP----
>ENSRNOG00000037079/1-683
--------MSFQSTLSLSLDSPTHAICMVGMEITLDISGCAPDKCESFTIRGSPRVLIHISSS--VITGKED
AVVWRPMKQPTEALVRMVSPSPTVDEDKVLVSYYCPDKEVPTATAVLFLTGIEVSLEADIYRDGQLDMPSDK
QAKKRWMWGLNGWGAILLVNCNPETPSQS-DE-QSGAQESPREIQNLSQMTLTVEGPTSILRDYQLLLHTSK
EEAKKTRVYWAHEG-ASAYELVVGPDKPFHILPT----FDTRGKETFYIEATEFPSPSFSGLISFSVSLIEQ
AHDQLIPETPLYKDTVMFRVAPYIFLPSTQLPLEVYLCRELQLQGFVDSVTKLSEKSNVQVASVYEDPNRQG
RWLQDEMAFCYTQAPHKTLSFILDTPRVAKLEDFPMKYSLSPGVGYLIQETEDHRVASLDSVGNLIVSPPVK
AQGKDYPLGRILIGGSFYPSFEGRDMSKALRDFVYAQQVQAPVELFSDWLMTGHVDQFLCFVPVDDKNSDRK
GFRLLLASPSACYELFEQKQKEGFGDVTLFEDVRAEQLISNGREVKTISQILADENLREQNAYVEKCISLNR
TLLKKELGLVDNDFIRIPQLFCLEPLTNVPSNQQTSKLFARPYFPDMLQMIVLGKNIGIPKPFGPQIKGTCC
LEETVCGLLEPLGLKCTFIDDFDCYLANIGDACASVIVNRVPFAFKWWKMNP----
>ENSNGAG00000014612/1-682
--------MSFQSTVRLSLDSPTRVVCVLGMEISLDISGCAPKKCESFTIRGSPRVLIHICSS--VITGKED
SIVWQPLSQSTEALVRMVSPSPAVDEDKVLVSYYCTDEETPAATAVLYLTGIEVSLEADIYRDGQLDMPSDK
QAKKKWMWGPSGWGAILLVNCSPTQAGQP-TG-QD-GKIFSEEIKNLSRMTLNVEGPNCILKNYRLVLHTSE
EEAAKTRVYWPQKDTSCDFELVMGPGKPSYALSP----LEDRGKEIFYVEATEFPSASFPGLISFSISLVEG
SQDLSIPDIPIYRDTVVFRVAPYIFTPSTQMPIEVYLCRELQLQGFVDTVIRLSERSNVQVASVYEDPSRLG
RWLQDEMAFCYTQAPHKTMSFILDTPRVTKLEDYPMKYSLSPGVGYLIRQTKHHQVTSLDTIGNLMVSPPVK
AQGKEYPLGRILVGGSFYPSSESRTMSKDLRDFVCAQQVQAPVEIFSDWLMTGHVDEFMCFVPA-DKSENKK
GFRLLLASPSACYELFEEKQKEGYGGATLFEDVRADQLLSNGREARTISQLLADKSLRKQNDYVEKCISLNR
ALLKKELGLVDKDIITIPQLFCLEQLTNVPSDQQTSKSFARPYFPDMLQMIVLGKNLGIPKPFGPQIKGTCC
LEEKICQLLEPLGLKCTFIDDFDCYLTNIGDSCACAIVHRVPFAFKWWKMIL----
>ENSJJAG00000019685/1-672
--------MSFQSTLRLSLDSPTHAVCVLGMETSLDISGCAPKKCEFFTIRGSPRVLIHVSSV--VITGKED
AVVWRSLYHPTEAVVRMMSPSPEVDDDKVLVSYYCPDEEVPMATAMLYLTGIEVSLEADIYRDGQLDMPSDK
QAKKKWMWGPSGWGAILTVNCNLVNMDEL-VN-KD------NNIKTLSRMTLSVKGPSCILKNYQLVLHTSK
EEASKTRVYCS-RG----LSLVLGPNKHFYALPP----LEDHGTETFYVEATEFPSASFSGLISFSVSLVER
SRDVSIPEIPIYKDTVVFRVAPCIFTPSTQMPLEVYLCSELQLQGFADTVSRLSERSDVHVVSVYEDPSRLG
RWLQDEMAFCYTQAPHKTVSLVLDTPRATELEDYPMKYSLSPGIGYMTQNTEDHQVASLDSIGNMIVSPPVK
AQGKEYPLGRVLLGSSFHPSPESRTMSKKLRDFLWAQQVQAPVELFSDWLMTGHVDEFMCFVPI-DDHEDRK
GFRLLLASPSACYNLLQEKQKEGYGDATLFGEVRTDQLHSNGREARTITQLLTDKSLRKQNDYVEKCICLNR
TILKKELGLVEKDIIAIPQLFCLEQLTNVPSDQQTPKLFARAYFPNMLRMIVLGQNLGIPKPYGPQIKGTCC
LEEKMCRLLEPLGLKCTFIDDFDCYLADIGDFCPCVSVRRAPFAFKWWKMMP----
>ENSSTOG00000019498/1-674
-------------VHLTLDTPAHAICVLGTEISLDISRCAPKNCQSFTVRASPRVLVDVAGT--VISGKED
AVICRSLLDTSHVLVRMVSPSASVDEDKVLVSYYHPNEEVPMATALLYLTGIEVNLEADIYRDGQLEVPSDK
QAKKKWVWGPNGWGAILLVNCSPANMSQI-TD-KS--KVFSDEIRKLSQMSLTVQGPSCILKNYKLILHTSK
EEAEKTRVYRPQKD-SCAYQLVLGLGWHCHALGP----LETLRKETFYVEATEFPSASFSGLVSFSVSLVEE
SQDLLIPEIPIYKDTVVFRVAPCIFTPSTQMPLEVYLCRELQLQGFVNTVTELSEKSDSQVAPVYEDPNRLG
RWLQDEMAFCYTQGPHRTVSFILDTPRAVHLEDFPMKYSLSPGIGYMIRRTEDHRVASLDSIGNLMVSPPVK
AHGKEYPLGRVLIGSSFYPSAQGRAMSRTLQDFLCAQQVQAPVELFSDWLMTGHVDEFMGFVPIDTKS-DKK
GFRLLIASPSSCYKLFQEKQEEGYGEATLFEDIRADQLLANGRKARTINQLLADESLRRQNDYVERCIDLNR
AILKRELGLGEKDIIDVPQLFCLERLTNIPSEQQTPKLLARAYFPNMLCMIVMDMNLGIPKPFGPQIKGICC
LQEKVRLLLEPLGLTCTFIDDFDCYLTEVGDFCPCANVLRVPFDFKWWKMVP----
>ENSUPAG00010017599/1-685
VVSMECRAVAFQSTVHLTLDTPAHAICVLGTEISLDISRCAPKNCQSFTVRASPRVLVDVAGT--VISGKED
AVICRSLLDTSHVLVRMVSXSASVDEDKVLVSYYHPNEEVPMATALLYLTGIGEFLWGS---PGPSPPPENS
WSQKKWVWGPNGWGAILLVNCSPANMSQI-TD-KS--KVFSDEIRKLSQMSLTVQGPSCILKNYKLILHTSK
EEAEKTRVYRPQKD-SCAYQLVLGLGWHCHTLGP----LENLQKETFYVEATEFPSASFSGLVSFSVSLVEE

SQDLLIPEIPIYKDTVVFRVAPCIFTPSTQMPLEVYLCRELQLQGFVNTVTELSEKSNSQVAPVYEDPNRLG
RWLQDEMAFCYTQGPHRTVSFVLDTPRAINLEDFPMKYSLSPGIGYMTRRTEDHRVASLDSIGNLMVSPPVK
AHGKEYPLGRVLIGSSFYPSAQGRAMSKTLQDFLCAQQVQAPVELFSDWLMTGHVDEFMGFVPIDTKN-DKK
GFRLLIASPSSCYKLFQEKQEEGYGEATLFEDIRADQLLANGRKARTINQLLADESLRRQNDYVERCIDLNR
AILKRELGLGEKDIIDVPQLFCLERLTNIPSDQQTPKLLARAYFPNMLCMIVMDMNLGIPKPFGPQIKGICC
LQEKVRLLLEPLGLACTFIDDFDCYLTEVGDFCPCANVLRVPFDFKWWKMVP----
>ENSMMMG00000020848/1-685
---MECRAIAFQSTVHLSLDIPAHAICVLGTEISLDISRCAPKNCESFTVRASPRVLVDVAGT--VISGKED
AVICRSLLDTSHVLVRMVSPSVSVDEDKVLVSYYHPNEEVPMATALLYLTGIEVNLEADIYRDGQLEVPSDK
QAKKKWVWGPNGWGAILLVNCGPADVSQI-TD-KS--KVFSDEIRKLSQMSLTVQGPSCILKNYKLVLHTSK
EEAEKTRVYRPQKD-SCAYQLVLGLGWHCHALGP----LENFQKETFYVEATEFPSASFSGLISFSVSLVEE
SQDLLIPEIPIYKDTVVFRVAPCIFTPSTQMPLEVYLCRELQLQGFVNTVTELSEKSDSQVAPVYEDPNRLG
RWLQDEMAFCYTQGPHRTVSFILDTPRAINLEDFPMKYSLSPGIGYMIRRTEDHRVASLDSIGNLMVSPPVK
AHGKEYPLGRVLIGSSFYPSAQGRAMSKTLQDFLCAQQVQAPVELFSDWLMTGHVDEFMGFVPIDTKS-DTK
SFRLLIASPSSCYKLFQEKQEEGYGEATLFEDIRADQLLANGRKARTINQLLADKSLRRQNDYVERCINLNR
AILKRELGLGEDDIIDVPQLFCLERLTNIPSDQQTPKLLARAYFPNMLCMIVMDMNLGIPKPFGPQINGICC
LQEKVRQLLEPLGLACTFIDDFDCYLTEVGDFCPCANVRRVPLDFKWWKMVP----
>ENSSVLG00005001183/1-688
--VSGGRAMSFQSIVHLSLDSPAHAICVLGMEISLDISRCAPKKCESFTVRASPRVLIDISGT--VITGKED
AVICRTLSSTMHVLVRMASPSLSVDEDKVLVSYYRPNEEVPTATAVLYLTGIEVSLEADIYRDGQFEVPSDK
QAKKKWVWGPSGWGAILLVNCSPTDVSQL-RD-KT-NKEFLDEMKNLSQMTLTVQGPSCILKNYRLVLHTSE
EEAEKARVYWAQKD-AFAFELVLGPGRHCHALGP----LENHQKETFYVEATEFPSASFSGLISFSASLVEE
SRDLLIPEIPTYKDTVVFRVAPCIFTPSTQMPLEVYLCRELQLQGFVSSVTELSEKSNSQVASVYEDPSRLG
RWLQDEMAFCYTQGPHKTVSFVLDTPRALELEDFPMKYSLSPGVGYVTRRTEDHRVASLDSIGNVMVSPPVK
AQGKEYPLGRVLIGSSFYPSAEGRTMSRSLRDFLCAQQVQVPVELFSDWLMTGHVDEFMCFVPVDAKSEDKK
GFRLLMASPSSCYELFQEKQEEGYGEAALFEEVRADQLLSNGREARTIDQLLADESLRKQNNYVERCLGLNR
AVLKRELGLVEEDIIDVPQLFCLEQLTNVPSDQQTPKLLARAYFPNMLRMIVMGTNLGIPKPFGPQIKGICC
LEGKIRQLLQPLGLMCTFIDDFDCYLTDVGDFCPCANVRRVPFAFKWWKMVP----
>ENSHGLG00000019075/1-680
--------MSLQSVVQLSLDSPVHAVCVLGSCLVVDISGCAPKKCDSFMLTCSPKVLVVLKSQ--VTSGRDK
TSIWHPLSEPMSALVMMVLPSSVVDEDKVLVSYYCPYEEAPLATAVLYLTGIEVSLKTDIYRDGQQEMPLNK
QMKEKWLWGPTGWGAILLVNCSPAEVNQD------TKTVLSEDIKDLSQMTLTVQGPTSALKNYQLVLHTSE
EEAEKARVYLPHGDTSGAFQIVLGPGCNSHTFNP----LVNLGKETLYVEAIEFPSANFAGLISFSVSLIED
SPVPSIPETPVYKDTVVFRVAPCIFTPSTQMPLEVYLYRDVKIQGFVDMVAALSEKTDSQVVSVYEDPNRLG
RWLQDEMAFCYTEAPQKMTSFVLDTPRATKLDEFPMKYALSPGVGYVIQHTKDHTVASLDTIGNLMVSPPVR
AQGKEYPLGRVLFGGSFYPSAESRMMSKSLCDFLHAQQVQAPVELFSDWLMTGHLDEFMCFIPVEEKKESSK
GFRLLLASPRACYKLLQEKQEEGYGNATLFEGVSPSLLRSNGREAKTINQFLADKSLRNQNDYVERCIRLNR
AILREKLGLLEDDIIDIPQLFCLEQLTNVPSDQQPCRFFARPYFPDVLRMIVMGKDLGIPKPFGPQIKGTCC
LEEKICQLLEPLGLKCTFIDDFDCYLTDVGDFCACTNIRRVPFAFKWWKMVP----
>ENSODEG00000016131/1-673
--------MSHLSVIHLSLDVPVHGVCVLGTSLVVDISGCAPKDCTSFTITCSPRVLVALKSK--VTSGKDK
ATVWHSLAEPVCALVKMVLPSSDVDEDKVLVSYYRANEEVPLVTAVLYLTGIEVSLQTDVYRDG-HEMPVNK
QTKETWVWGPAGWGAILLVNSSPADVNPD------TQMVQRQPQKLLTQMTLSIQGPTCILKNYRLVFHTSE
EEAKKARVFWPHG------ELLLGPDRNSYIFNP----LVKLGKETIYVEAIEFPSANFSGLISFSVSLIED
PPISCLPEIPVYKDTVVFRVAPCIFTPSTQMPLEVYLYRDVKVQGFVDAVVALSEKTDSQMASVYEDSNRSG
KWLQDEMAFCYTEMPHKMSSFVLDTPRAIALDDFPMKYALTPGIGYVTQDTKDHSVASLDTIGNLMVSPPVK
AQGKDYPLGRVLFGSSFYPSIDGRDMSKSLRDFLYAQQVQAPVELFSDWLMTGRLEEFMCFVPVENQSGDAK
DFRLLLASPCACYSLLREKQEEGYGDATLFEGVSPDLLHANGREAKTINQFLSDKSLKNQNDYAEKCIHLNR
AILKEKLGLQDEDIIDVPQLFCLEKLTNVPSDQQPHRLFARPYFPDMLRMMVMGKELGIPKPFGPQIKGTCC
LEEKICHLLEPLGLKCTFIDDFDCYMTDVGDACACTNIRRVPLAFKWWKMVP----
>ENSCLAG00000003328/1-672
--------MSHQSIVHLCLDVPVHGVCVLGTSLIVDISGCAPKNCESFTIICSPRVLVVLNSK--VTSGKDK
AIVWHSLSKPVSALVRMVLPSSNVDEDKVLVSYYRANEEVPLATAVLYLTGIEVSLQADVYRDGQHEMPVNK
QTKEKWVWGPSGWGAILLVNCSPADVSQD------TKTVISEDIKDLSQMILSVQGPTCVLKNYQLVLHTSV
EEADKARVYWPHG------EIVLGPGHNSYTFSP----LVNLGKETFYVEGIEFPSADFSGLISFSISLIED
PP--SIPEAPVYKDTVVFRVAPCIFTPSTQMPLEVYLYRDVKVQGFVDTVVALSEKTDCQVASVYEDPSRQG
RWLQDEMAFCYTETPHKMTSFVLDTPRAAKLDEFPMKYALSPGVGYIIQDTKDHTVASLDTIGNLMVSPPVK
AQGKEYPLGRVLFGSSFYPSTEGRAMSKSLRDFLYAQQVQAPVELFSDWLMTGHVDEFMCFVPVEDKGEDAK
GFRLLLASPCACYKLFREKQEEGHGDATLFEGVSTDLLHSNGREAKTINQFLADESLRNQNNYVEKCIHLNR
AILKERLGLQEEDIIDIPQLFCLEQLTNVPSDQQPRKLFARPYFPDVLQMIVMGKDLGIPKPFGPQIKGTCC
LEEKICHLLEPLGLSCTFIDDFDCYLTDVGDSCACTNIRRVPFAFKWWKMLP----
>ENSCPOG00000024187/1-667
--------MSNQSVVHLSLDDPAHGVCVLGTSLVVDVSGCAPMNCESFTITCSSRVLVVLNSK--VTSGKDK

GIIWHSISDRRHALVRMVLPSSELDEDKVLVSYYRANEEAPLATAVLYLTGIEVSLQTDVYRDGQQEMPINK
QIKEKWVWGPSGWGAVLLVNCSPADLRQE------SKMVVSEHIKDLSQMTLNVNGPNCALKNYHLVLHTSA
EEAEKTRVYWPQG------EVVLGPDRSSYTFTP----LENL-EETFYVEALEFPSASFSGLISFSISLIED
PP------GPVYKDTVVFRVAPCIFTPSTQMPLEMYLYRDVKIQGFVDAVTALSEKSDSQVASVYEDFNRMG
RWLQDEMAFCYTETPNKMMSFVLDTPRTTTPDEFPMKYALSPGVGYLIQNITDHTVASLDTIGNLMVSPPVK
AQGKEYPLGRVLFGSSFYPSSEGRAMSKNLRDFLYAQQVQAPVELFSDWLMTGHVDEFMCFVPVEDKSNDSK
GFRLLLASPCACYNLFREKQEEGFGGSVLFGGVNPDTLHANGREAKTINQLLADQTMRNQNNYVEKCIHLNR
AILKEKLGLQEEDIIDVPQLFCLEELTNVPSDQQPCKLFARPYFPDVLRMIVMGKDLGIPKPFGPQIKGTCC
LEEKICHLLEPLGLKCTFIDDFECYLTCVGDPCACTNIRRMPFAFKWWNMVP----
>ENSDORG00000005081/1-658
--------MPVQHLIHLSLDSPVHAVCVLGMEVSVDINGCAPTECTSFTIHGSPGVLIGSSST--AFTRKED
TTVSRPLPSPAHVPLQMLSLSFSEDEDKVEVSYFCPNEDSPAATAVLYLTSIYVSLDADIYRDGQLEMASDK
LAKKKWVWGPNGWGAILLVNCNSVDLSQV-TD----KAIFSKEIKTLAEVTLTVQGPSSTLKNYGLVLHTSE
EEAKKIRIYWPQ-DAQSTFKLVVGPSQHFHTLSP----LES-MEQTLYAEAIDFPSSSFSGLISFSLSLVEQ
PQDSAIPETPIYKDTVVFRVAPCIFMPSTQMPLEVYLCREVQFTGSSEVLLRLS----------------G
SWLTDEMAFCYSQAPHGTISMVIDSPRTAKLEDFPMKYSLSPGISYMTHSTEDHQVASLDSVGNMMVSPPVK
AQGKDYPLGRILIGSSFYP---SRTMGKSLQAFLYAQQVQAPLELFSDWLMTGHVDEFMCFVPVEN-SRDTK
DFRLLLASPSACYELFQKKQREGYGNATLFEEVRTDQLISNGRKAKTIDQLLADEKLRKENDYVEKCIYLNR
ILLKRELGLVEKDIIDIPQLFYLEKLTNIPSDQQTTKLFAKPYFPDLLQMIVMGNNLGIPKPFGPQIKGICC
LEEKVCHLLEPLGLQCTFIDDFDCYLTDVGDFCSCANIRRVPFAFKWWKVMP----
>ENSOCUG00000025853/1-693
GVSLEGRAMSFQNTLCLSPDSPVHAICVLGTEINLDLSRAAPDNCEFFTVSGSQRVLVNVCDV--VITEKEA
ATIWQPLSDSTHVLVQMVSPSPAVDEDKVLVSYYRPKEEEPAATAVLYLTGIEVSLQVDVYRNGHMQVPTEK
QEKEKWVWGPKGRGPILLVNTGPANVDQL-IDKETNKVFLSDEIKNLSRMILNLQGPSCVLKKYRLVLHTSR
EEAEKARVYWPQKDGSSAFELVLGPGRHSYTFTS----IENPLKETFYLEATEFPSASFSGLISYSASLVEE
SEDPSIPETPVYKDTVVFRVAPCIFTPSTQMPLEVYLCRELQLQGFVNMVTELSEKSNSQLASVYEDPNRLG
RWLQDEMAFCYTQAPHKTVSLILDTPRATKLEDFPMKYSLSPGVGYVVQHTEDHRVASMDSIGNLMVSPPVQ
VQEKEYPLGRVLIGSSFYPSVEGRAMSKSLRDFLYAQQVQAPVELFSDWLMSGHVDEFMCFVPTDDKSGSKK
GFLLLLASPSSCYNLFKEKQKEGYGDAVLFEGVRGDQLLSNGREARTINQLLADESLRKQNDYVEKCLRLNR
TILKSELGLAEEDIVEVPQLFCLEQLSNVPSEQQSRKPLARPYFPDLLQMIVMDKNLGVPKPFGPQIKGSCC
LEQELCRLLEPLGFKCSFIDDFDCYLTQVGDFCACANIRRVPFAFKWWNMTP----
>ENSPTRG00000052623/1-693
MVSVEGRAMSFQSIVHLSLDSPVHAVCVLGTEICLDLSGCAPQKCQCFTIHGSGRVLIDVANT--VISEKED
ATIWWPLSDPTYATVKMTSPSPSMDADKVSVTYYGPNEDAPVGTAVLYLTGIEVSLEVDIYRNGQVEMSSDK
QAKKKWIWGPSGWGAILLVNCNPADVGQQLEDKKTKKVIFSEEITNLSQMTLNVQGPSCILTKYRLVLHTSK
EESKKARVYWPQKDNSSTFELVLGPDQHAYTLA-----LGNHLKETFYVEAIAFPSAEFSGLISYSVSLVEE
SQDPSIPETVLYKDTVVFRVAPCVFIPCTQVPLEVYLCRELQLQGFVDTVTKLSEKSNSQVASVYEDPNRLG
RWLQDEMAFCYTQAPHKTTSLILDTPQAADLDEFPMKYSLSPGIGYMIQDTEDHKVASMDSIGNLMVSPPVK
VEGKEYPLGRVLIGSSFYPSAEGRAMSKTLRDFLYAQQVQAPVELYSDWLMTGHVDEFMCFIPTDDKNEGKK
GFLLLLASPSACYKLFREKQKEGYGDALLFDELRADQLLSNGREAKTIDQLLADESLKKQNEYVEKCIHLNR
DILKTELGLVEQDIIEIPQLFCLEKLTNIPSDQQPKRSFARPYFPDLLRMIVMGKNLGIPKPFGPQIKGTCC
LEEKICCLLEPLGFKCTFINDFDCYLTEVGDICACANIRRVPFAFKWWKMVP----
>ENSPPAG00000040946/1-694
MVSVEGRAMSFQSIVHLSLDSPVHAVCVLGTEICLDLSGCAPQKCQCFTIHGSGRVLIDVANT--VISEKED
ATIWWPLSDPTYATVKMTSPSPSMDADKVSVTYYGPNEDAPVGTAVLYLTGIEVSLEVDIYRNGQVEMSSDK
QAKKKWIWGPSGWGAILLVNCNPADVGQQLEDKKTKKVIFSEEITNLSQMTLNVQGPSCILTKYRLVLHTSK
EESKKARVYWPQKDNSSTFELVLGPDQHAYTLAL----LGNHLKETFYIEAIAFPSAEFSGLISYSVSLVEE
SQDPSIPETVLYKDTVVFRVAPCVFIPCTQVPLEVYLCRELQLQGFVDTVTKLSEKSNSQVASVYEDPNRLG
RWLQDEMAFCYTQAPHKTTSLILDTPQAADLDEFPMKYSLSPGIGYMIQDTEDHKVASMDSIGNLMVSPPVK
VEGKEYPLGRVLIGSSFYPSAEGRAMSKTLRDFLYAQQVQAPVELYSDWLMTGHVDEFMCFIPTDDKNEGKK
GFLLLLASPSACYKLFREKQKEGYGDALLFDELRADQLLSNGREAKTIDQLLADESLKKQNEYVEKCIHLNR
DILKTELGLVEQDIIEIPQLFCLEKLTNIPSDQQPKRSFARPYFPDLLRMIVMGKNLGIPKPFGPQIKGTCC
LEEKICCLLEPLGFKCTFINDFDCYLTEVGDICACANIRRVPFAFKWWKMVP----
>ENSG00000276747/1-694
MVSVEGRAMSFQSIIHLSLDSPVHAVCVLGTEICLDLSGCAPQKCQCFTIHGSGRVLIDVANT--VISEKED
ATIWWPLSDPTYATVKMTSPSPSVDADKVSVTYYGPNEDAPVGTAVLYLTGIEVSLEVDIYRNGQVEMSSDK
QAKKKWIWGPSGWGAILLVNCNPADVGQQLEDKKTKKVIFSEEITNLSQMTLNVQGPSCILKKYRLVLHTSK
EESKKARVYWPQKDNSSTFELVLGPDQHAYTLAL----LGNHLKETFYVEAIAFPSAEFSGLISYSVSLVEE
SQDPSIPETVLYKDTVVFRVAPCVFIPCTQVPLEVYLCRELQLQGFVDTVTKLSEKSNSQVASVYEDPNRLG
RWLQDEMAFCYTQAPHKTTSLILDTPQAADLDEFPMKYSLSPGIGYMIQDTEDHKVASMDSIGNLMVSPPVK
VQGKEYPLGRVLIGSSFYPSAEGRAMSKTLRDFLYAQQVQAPVELYSDWLMTGHVDEFMCFIPTDDKNEGKK
GFLLLLASPSACYKLFREKQKEGYGDALLFDELRADQLLSNGREAKTIDQLLADESLKKQNEYVEKCIHLNR
DILKTELGLVEQDIIEIPQLFCLEKLTNIPSDQQPKRSFARPYFPDLLRMIVMGKNLGIPKPFGPQIKGTCC

LEEKICCLLEPLGFKCTFINDFDCYLTEVGDICACANIRRVPFAFKWWKMVP----
>ENSGGOG00000015257/1-694
MVSVEGRVMSFQSIVHLSLDSPVYAVCVLGTEICLDLSGCAPQKCQCFTVHGSGRVLIDVANT--VISEEED
ATIWWPLSDPTYATVKMTSPSPSVDADKVSVTYYGPNEDAPVGTAVLYLTGIEVSLEVDIYRNGQVEMSSDK
QAKKKWIWGPSGWGAILLVNCNPADVGQQLEDKKTKKVIFSEEITNLSQMTLNVQGPSCILKKYRLVLHTSK
EESKKARVYWPQKDDSSTFELVLGPDQHVYTLAL----LGNHLKETFYVEAIAFPSAEFSGLISYSVSLVEE
SQDPSIPETVLYKDTVVFRVAPCVFIPCTQVPLEVYLCRELQLQGFVDTVTKLSEKSNSQVASVYEDPNRLG
RWLQDEMAFCYTQAPHKTTSLILDTPQAADLDEFPMKYSLSPGIGYMIQDTEDHKVASMDSIGNLMVSPPVK
VQGKEYPLGRVLIGSSFYPSAEGRAMSKTLRDFLYAQQVQAPVELYSDWLMTGHVDEFMCFIPTDDKNEGKK
GFLLLLASPSACYKLFREKQKEGYGDALLFDELRADQLLSNGREAKTIDQLLADESLKKQNEYVEKCIHLNR
DILKTELGLVEQDIIEIPQLFCLEKLTNIPSDQQPKRPFARPYFPDLLRMIVMGKNLGIPKPFGPQIKGTCC
LEEKICCLLEPLGFKCTFINDFDCYLTEVGDICACANIRRVPFAFKWWKMVP----
>ENSPPYG00000001806/1-694
VVSVEGRAMSFQSIVHLSLDSPAHAVCVLGTEICLDLSGCAPQKCQCFTIHGSGRVLINMANT--VISEKED
ATIWWPLSDPTYATVKMISPSPSVDGDKVSVTYYGPNEDAPVGTAVLYLTGIEVSLEVDIYRNGQVEMSSDK
QAKKKWIWGPSGWGAILLVNCNPADVGQQLEDKKTKKVIFSEEITNLSQMTLSVQGPTCILKKYRLVLHTSK
EESKKARVYWPQKDNSSTFELVLGPGQHAYTLAL----LGSHLKETFYVEAIAFPSAEFSGLISYSVSLVEE
SQDPSIPETLLYKDTVVFRVAPCVFIPCTQVPLEVYLCRELQLQGFVDTVTELSEKSNSQVASVYEDPNRLG
RWLQDEMAFCYTQAPHKTTSLILDTPQATDLDEFPMKYSLSPGVGYMIQDTEDHNVASMDSIGNLMVSPPVK
VQGKEYPLGRVLIGSSFYPSAEGRAMSKTLRDFLYAQQVQAPVELYSDWLMSGHVDEFMCFIPTDDKNEGEK
GFQLLLASPSACYKLFREKQKEGYGDALLFDELRADQLLSNGREAKTIDQLLADESLKKQNEYVEKCIHLNR
DILKKELGLVEQDIIEIPQLFCLEKLTNIPSDQQPKRPFARPYFPDLLRMIVMGKNLGIPKPFGPQIKGTCC
LEEKICCLLEPLGFKCTFINDFDCYLTEVGDICACANIRRVPFAFKWWKMVP----
>ENSNLEG00000007782/1-642
VVSVEGRAMSFQSIVHLSLDSPAHAVCVLGTEI--------------YSAHGVG-----------LCSR---
-----PLPSNTAYGFRMLSP-------QVSVTYYGPNEDAPMGTAVLYLTGIEVSLEVDIYRNGQVEMSSDK
QAKKKWIWGPSGWGAILLVNCNPADVGQQLEDKKTKKVIFSEEITNLSQMTLNVQGPSCILKKYRLVLHTSK
EESKKARVYWPQKDNSSTFELVLGPGQHAYTLAL----LGSHLKETFYVEAIAFPSAEFSGLISYSVSLVEE
SQDPSIPETLLYKDTVVFRVAPCVFIPCTQVPLEVYLCRELQLQGFVDTVTELSEKSNSQVASVYEDPNRLG
RWLQDEMAFCYTQAPHKTTSLILDTPQAADLDEFPMKYSLSPGIGYMIQDTEDHKVASMDSIGNLMVSPPVK
VQGREYPLGRVLIGSSFYPSAEGRAMSKTLRDFLYAQQVQAPVELYSDWLMTGHVDEFMCFIPTDDKNEGKK
GFQLLLASPSACYKLFREKQKEGYGDALLFDELRADQLLSNGKGTPF-----------AQN---RKCIHLNR
DILKTELGLVEQDIIEIPQLFCLEKLTNIPSDQQPKRPFARPYFPDLLRMIVMGKNLGIPKPFGPQIKGTCC
LEEKICCLLEPLGFKCTFINDFDCYLTEVGDICACANVRRVPFAFKWWKMVP----
>ENSMLEG00000029519/1-694
MVSVEARAMSFQSIVRLSLDSPAHAVCVLGTEICLDLSGCAPQKCQCFTIHGSGRVLIDVANT--VISEEED
ATIWWPLSDPTYATVKMISPSPSVDADKVSITYYGPNEDAPVGTALVYLTGIEVSLEVDIYRSGQVEMSSDK
QAKKKWIWGPSGWGAILLVNCNPADVGQQPEDKKTKKVIFSEEIKNLSQMTLSVQGPSCILKKYRLVLHTSK
EESKKARVYWPQKDNSSTFELVLGPGQHAYTLAL----LGNHLKETFYVEAIAFPSAEFSGLISYSASLVEE
SQDPSIPETLLYKDTVVFRVAPCVFVPCTQVPLEVYLCRELQLQGFVDTVTELSEKSNSQVASVYEDPNRLG
RWLQDEMAFCYTQAPHKTTSLILDTPQATDLDEFPMKYSLSPGIGYMIQDIEDHKVASMDSIGNLMVSPPVK
VQEKEYPLGRVLIGSSFYPSAEGRAMSKTLRDFLYAQQVQAPVELYSDWLMTGHVDEFMCFIPTDDKNEGKK
GFQLLLASPSACYKLFREKQKEGYGDALLFDELRADQLLSNGREARTIDQLLADESLKKQNEYVEKCIHLNR
DILKTELGLVEQDIIEIPQLFCLEKMTNIPSDQQPKRPFARPYFPDLLRMIVMGKNLGIPKPFGPQIKGTCC
LEEKICRLLEPLGFKCTFINDFDCYLTEVGDICACANVRRVPFAFKWWKMVP----
>ENSPANG00000031439/1-694
MVSVEGRAMSFQSIVRLSLDSPAHAVCVLGTEICLDLSGCAPQKCQCFTIHGSGRVLIDVANT--VISEEED
ATIWWPLSDPTYATVKMISPSPSVDADKVSITYYGPNEDAPVGTALVYLTGIEVSLEVDIYRSGQVEMSSDK
QAKKKWIWGPSGWGAILLVNCNPADVGQQPEDKKTKKVIFSEEIKNLSQMTLSVQGPSCILKKYRLVLHTSK
EESKKARVYWPQKDNFSTFELVLGPGQHAYTLAL----LGNHLKETFYVEAIAFPSAGFSGLISYSASLVEE
SQDPSIPETLLYKDTVVFRVAPCVFVPCTQVPLEVYLCRELQLQGFVDTVTELSEKSNSQVASVYEDPNRLG
RWLQDEMAFCYTQAPHKTTSLILDTPQATDLDEFPMKYSLSPGIGYMIQDIEDHKVASMDSIGNLMVSPPVK
VQEKEYPLGRVLIGSSFYPSAEGRAMSKTLRDFLYAQQVQAPVELYSDWLMTGHVDEFMCFIPTDDKNEGKK
GFQLLLASPSACYKLFREKQKEGYGDALLFDELRADQLLSNGREARTIDQLLADESLKKQNEYVEKCIHLNR
DILKTELGLVEQDIIEIPQLFCLEKMTNIPSDQQPKRPFARPYFPDLLRMIVMGKNLGIPKPFGPQIKGTCC
LEEKICHLLEPLGFKCTFINDFDCYLTEVGDICACANVRRVPFAFKWWKMVP----
>ENSCATG00000040335/1-694
MVSVEARAMSFQSIVRLSLDSPAHAVCVLGTEICLDLSGCAPQKCQCFTIHGSGRVLIDVANT--VISEEED
ATIWWPLSDPTYATVKMISPSPSVDADKVSITYYGPNEDAPVGTALVYLTGIEVSLEVDIYRSGQVEMSSDK
QAKKKWIWGPSGWGAILLVNCNPADVGQQPEDKKTKKVIFSEEIKNLSQMTLSVQGPSCILKKYRLVLHTSK
EESKKARVYWPQKDNSSTFELVLGPGQHAYTLAL----LGNHLKETFYVEAIAFPSAEFSGLISYSASLVEE
SQDTSIPETLLYKDTVVFRVAPCVFVPCTQVPLEVYLCRELQLQGFVDTVTELSEKSNSQVASVYEDPNRLG
RWLQDEMAFCYTQAPHKTTSLILDTPQATDLDEFPMKYSLSPGIGYMIQDIEDHKVASMDSIGNLMVSPPVK

VQEKEYPLGRVLIGSSFYPSAEGRAMSKTLRDFLYAQQVQAPVELYSDWLMTGHVDEFMCFIPTDDKNEGKK
GFQLLLASPSACYKLFREKQKEGYGDALLFDELRADQLLSNGREARTIDQLLADESLKKQNEYVEKCIHLNR
DILKTELGLVEQDIIEIPQLFCLEKMTNIPSDQQPKRPFARPYFPDLLRMIVMGKNLGIPKPFGPQIKGTCC
LEEKICRLLEPLGFKCTFINDFDCYLTEVGDICACANVRRVPFAFKWWKMVP----
>ENSMMUG00000022437/1-694
MVSVEGRAMSFQSIVRLSLDSPAHAVCVLGTEICLDLSGCAPQKCQCFTIHGSGRVLIDVANT--VISEEED
ATIWWPLSDPTYATVKMISPSPSVDADKVSITYYGPNEDAPVGTALVYLTGIEVSLEVDIYRSGQVEMSSDK
QAKKKWIWGPSGWGAILLVNCNPADVGQQPEDKKTKKVIFSEEIKNLSQMTLSVQGPSCILKKYRLVLHTSK
EESKKARVYWPQKDNSSTFELVLGPGQHAYTLAL----LGNHLKETFYVEAIAFPSAEFSGLISYSASLVEE
SQDPSIPETLLYKDTVVFRVAPCVFVPCTQVPLEVYLCRELQLQGFVDAVTELSEKSNSQVASVYEDPNRLG
RWLQDEMAFCYTQAPHKTTSLILDTPQATDLDEFPMKYSLSPGIGYMIQDIEDHKVASMDSIGNLMVSPPVK
VQEKEYPLGRVLIGSSFYPSAEGRAMSKTLRGFLYAQQVQAPVELYSDWLMTGHVDEFMCFIPTDDKNEGKK
GFQLLLASPSACYKLFREKQKEGYGDALLFDELRADQLLSNGREARTIDQLLADESLKKQNEYVEKCIHLNR
DILKTELGLVEQDIIEIPQLFCLEKMTNIPSDQQPKRPFARPYFPDLLRMIVMGKNLGIPKPFGPQIKGTCC
LEEKICRLLEPLGFKCTFINDFDCYLTEVGDICACANVRRVPFAFKWWKMVP----
>ENSMFAG00000002560/1-694
MVSVEGRAMSFQSIVRLSLDSPAHAVCVLGTEICLDLSGCAPQKCQCFTIHGSGRVLIDVANT--VISEEED
ATIWWPLSDPTYATVKMISPSPSVDADKVSITYYGPNEDAPVGTALVYLTGIEVSLEVDIYRSGQVEMSSDK
QAKKKWIWGPSGWGAILLVNCNPADVGQQPEDKKTKKVIFSEEIKNLSQMTLSVQGPSCILKKYRLVLHTSK
EESKKARVYWPQKDNSSTFELVLGPGQHAYTLAL----LGNHLKETFYVEAIAFPSAEFSGLISYSASLVEE
SQDPSIPETLLYKDTVVFRVAPCVFVPCTQVPLEVYLCRELQLQGFVDAVTELSEKSNSQVASVYEDPNRLG
RWLQDEMAFCYTQAPHKTTSLILDTPQATDLDEFPMKYSLSPGIGYMIQDIEDHKVASMDSIGNLMVSPPVK
VQEKEYPLGRVLIGSSFYPSAEGRAMSKTLRGFLYAQQVQAPVELYSDWLMTGHVDEFMCFIPTDDKNEGKK
GFQLLLASPSACYKLFREKQKEGYGDALLFDELRADQLLSNGREARTIDQLLADESLKKQNEYVEKCIHLNR
DILKTELGLVEQDIIEIPQLFCLEKMTNIPSDQQPKRPFARPYFPDLLRMIVMGKNLGIPKPFGPQIKGTCC
LEEKICRLLEPLGFKCTFINDFDCYLTEVGDICACANVRRVPFAFKWWKMVP----
>ENSMNEG00000039952/1-694
MVSVEGRAMSFQSIVRLSLDSPAHAVCVLGTEICLDLSGCAPQKCQCFTIHGSGRVLIDVANT--VISEEED
ATIWWPLSDPTYATVKMISPSPSVDADKVSITYYGPNEDAPVGTALVYLTGIEVSLEVDIYRSGQVEMSSDK
QAKKKWIWGPSGWGAILLVNCNPADVGQQPEDKKTKKVIFSEEIKNLSQMTLSVQGPSCILKKYRLVLHTSK
EESKKARVYWPQKDNSSTFELVLGPGQHAYTLAL----LGNHLKETFYVEAIAFPSAEFSGLISYSASLVEE
SQDPSIPETLLYKDTVVFRVAPCVFVPCTQVPLEVYLCRELQLQGFVDTVTELSEKSNSQVASVYEDPNRLG
RWLQDEMAFCYTQAPHKTTSLILDTPQATDLDEFPMKYSLSPGIGYMIQDIEDHKVASMDSIGNLMVSPPVK
VQEKEYPLGRVLIGSSFYPSAEGRAMSKTLRGFLYAQQVQAPVELYSDWLMTGHVDEFMCFIPTDDKNEGKK
GFQLLLASPSACYKLFREKQKEGYGDALLFDELRADQLLSNGREARTIDQLLADESLKKQNEYVEKCIHLNR
DILKTELGLVEQDIIEIPQLFCLEKMTNIPSDQQPKRPFARPYFPDLLRMIVMGKNLGIPKPFGPQIKGTCC
LEEKICRLLEPLGFKCTFINDFDCYLTEVGDICACANVRRVPFAFKWWKMVP----
>ENSCSAG00000000865/1-694
MVSVEGRAMSFQSIVRLSLESPAHAVCVLGTEICLDLSGCAPQKCQCFTIHGSGRVLIDVANT--VISEEED
ATIWWPLSDPTYATVKMISPSPSVDTDKVSITYYGPNEDAPVGTAVVYLTGIEVSLEVDIYRSGQVEMSSDK
QAKKKWIWGPSGWGAILLVNCNPADVGQQPEDKKTKKVIFSEEIKNLSQMTLSVQGPSCILKKYRLVLHTSK
EESKKARVYWPQKDNSSTFELVLGPGQHAYTLAL----LGNHLKETFYIEAIAFPSAEFSGLISYSASLVEE
SQDPSIPETVLYKDTVVFRVAPCVFVPCTQVPLEVYLCRELQLQGFVDTVTELSEKSNSQVASVYEDPNRLG
RWLQDEMAFCYTQAPHKTTSLILDTPQATDLDEFPMKYSLSPGIGYMIQDIEDHKVASMDSIGNLMVSPPVK
VQEKEYPLGRVLIGSSFYPSAEGRAMSKTLRDFLYAQQVQAPVELYSDWLMTGHVDEFMCFIPTDDKNEGKK
GFQLLLASPSACYKLFREKQKEGYGDALLFDELRADQLLSNGREAKTIDQLLADESLKKQNEYVEKCIHLNR
DILKTELGLVEQDIIEIPQLFCLEKMTNIPSDQQPKRPFARPYFPDLLRMIVMGKNLGIPKPFGPQIKGTCC
LEEKICRLLEPLGFKCTFINDFDCYLTEVGDICACANVRRVPFAFKWWKMVP----
>ENSRROG00000039820/1-694
MVSVAGRAMSFQSIVRLSLDSPAHAVCVLGTEICLDLSGCAPQKCECFTIHGSGRVLIDVANT--VISEEED
ATIWWPLSNPTYATVKMISPSPSVDGDKVSIAYYGPNEDAPVGTAVVYLTGIEVSLEVDIYRSGQVEMSSDK
QAKKKWIWGPSGWGAILLVNCNPADVGQQPEDKKTKKVIFSEEIKNLSQMTLSVQGPSCILKKYRLVLHTSK
EESKKARVYWPQKDNSSTFELVLGPGQHAYTLAL----LGNHLKETFYVEAIAFPSAEFSGLISYSASLVEE
SQDPSIPETLLYKDTVVFRVAPCVFIPCTQVPLEVYLCRELQLQGFVDTVTELSEKSNSQVASVYEDPNRLG
RWLQDEMAFCYTQAPHKTTSLILDTPQATDLDEFPMKYSLSPGIGYMIQDTEDHKVASMDSIGNLMVSPPVK
VQEKEYPLGRVLIGSSFYPSTEGRAMSKTLRDFLYAQQVQAPVELYSDWLMSGHVDEFMCFIPTDDKNEGKK
GFQLLLASPSACYKLFREKQKEGYGDALLFDELRADQLLSNGREAKTIDQLLADESLKKQNEYVEKCIHLNR
DILKTELGLVEQDIIEIPQLFCLEKLTNIPSDQQPKRPFARPYFPDLLRMIVMGKNLGIPKPFGPQIKGTCC
LEEKICRLLEPLGFKCTFINDFDCYLTEVGDICACANVRRVPFAFKWWKMVP----
>ENSRBIG00000035034/1-694
MVSVAGRAMSFQSIVRLSLDSPAHAVCVLGTEICLDLSGCAPQKCECFTIHGSGRVLIDVANT--VISEEED
ATIWWPLSNPTYATVKMISPSPSVDGDKVSIAYYGPNEDAPVGTAVVYLTGIEVSLEVDIYRSGQVEMSSDK
QAKKKWIWGPSGWGAILLVNCNPADVGQQPEDKKTKKVIFSEEIKNLSQMTLSVQGPSCILKKYRLVLHTSK

EESKKARVYWPQKDNSSTFELVLGPGQHAYTLAL----LGNHLKETFYVEAIAFPSAEFSGLISYSASLVEE
SQDPSIPETLLYKDTVVFRVAPCVFIPCTQVPLEVYLCRELQLQGFVDTVTELSEKSNSQVASVYEDPNRLG
RWLQDEMAFCYTQAPHKTTSLILDTPQATDLDEFPMKYSLSPGIGYMIQDTEDHKVASMDSIGNLMVSPPVK
VQEKEYPLGRVLIGSSFYPSTEGRAMSKTLRDFLYAQQVQAPVELYSDWLMSGHVDEFMCFIPTDDKNEGRK
GFQLLLASPSACYKLFREKQKEGYGDALLFDELRADQLLSNGREAKTIDQLLADESLKKQNEYVEKCIHLNR
DILKTELGLVEQDIIEIPQLFCLEKLTNIPSDQQPKRPFARPYFPDLLRMIVMGKNLGIPKPFGPQIKGTCC
LEEKICRLLEPLGFKCTFINDFDCYLTEVGDICACANVRRVPFAFKWWKMVP----
>ENSCCAG00000036970/1-693
KVSVEGQAMSFQNIIRLSLDSPAHAVCVMGTEICLDLSGCAPEKCQCFTIRGSGRVMIDVANT--VISEKES
ATIWWPLSDATYATVKMISPSPSMDGDKVSVKYYGLNEDAPVGKAVLYLTGIEVSLEADIYRNGQVEVSRDK
QAKKKWTWGPSGWGAILLVNCNLADAGQQLED-KTNRVIFSEEIKNLSQMTLKVQGPSCLLKKYRLVLHTSE
EESKKARVYWPRKDNSSTFELVLGPSQHAYTLAL----LGDHLKETFYLEAMVFPSAEFSGLITYSVSLVEE
SQDPLIPETPLYKDTVVFRVAPCIFIPCTQVPLEVYLCRELQLQGFVDTVTELSEKSNSQVASVYEDPSRLG
RWLQDEMAFCYTQAPHKTTSLILDTPRAPNLDEFPMKYSLSPGIGYMTHNTEDHKVASMDSIGNVMVSPPVK
VQGKEYPLGRILMGSSFYPSVEGRAMSKTLRDFLYAQQVQAPVELYSDWLMTGHVDEFMCFIPTDEKSEGKK
GFQLLLASPGACYKLFQEKQKEGYGDAFLFDGVRADQLLSNRREAKTIDQLLADESLRKQNEYVEKCIHLNH
DILKTELGLVEQDIIEIPQLFCLEKLTNVPSDQQPKRLFARPYFPNLLQMIVMGKNLGIPKPFGPQIKGTCC
LEEKIRFLLEPLGFKCTFISDFDCYLADVGDICACATVRRVPFAFKWWKMVP----
>ENSCJAG00000007733/1-693
KVSVEGQAMSFQNVIRLSLDSPAHAVCVIGTEICLDISRCAPEKCQCFSIRGSERVMIDVANT--VFSEKES
ATIWWPLSDTPYATVKMISPSPSMDGDKVSVKYYGLNEDAPVGKAVLYLTGIEVSLETDIYRNGQVEVSRDK
QAKKKWTWGPSGWGAILLVNCNPADAGQQLED-ETKRVIFSEEIKNLSQMTLNVQGPSCLLKKFRLVLHTSE
EESKKARVYWPRKDNSSTFELVLGPSQHAYTLAL----LGDHLKETFYLEAMEFPSAEFSGLITYSLSLVEE
SQDPLIPETPLYKDTVVFRVAPCIFIPCTQVPLEVYLCRELQLQGFVDTVMELSKKSNSQVASVYEDPSRLG
RWLQDEMAFCYTQAPHKTTSLILDTPRAPNLDEFPMKYSLSPGIGYMIHDTEDHKVASMDSIGNVMVSPPVK
VQGKEYPLGRVLIGSSFYPSMEGRAMSKTLRDFLYAQQVQAPVELYSDWLMTGHVDEFMCFIPTDDKSEGKK
GFQLLLASPGACYKLFQEKRKEGYGDAFLFDGVRADQLLSNRREAKTIDQLLADESLRKQNEYVEKCIHLNH
DILKTELGLVEQDIIEIPQLFCLEKLTNVPSDQQPKRSFARPYFPNLLRMIVMGKNLGIPKPFGPQIKGTCC
LEEKIRCLLEPLGFKCTFVSDFDCYLADVGDICACATVRRVPFAFKWWKMVP----
>ENSCCAG00000026784/1-571
------------------MDSVTKVPHTASSQVALNVLPPPPKRLQ--------------------------
-------SEPTYQWTS-VSPSCEKH-----------------------------TL---------TEMSSHL
QATKKWTWGPSGWGAILLVNCNPADAGQQLED-KTNRVIFSEEIKNLSQMTLKVQGPSCLLKKYRLVLHTSE
EESKKARVYWPRKGNSSTFELVLGPSQHAYTLAL----LRDHLKETFYFKAMVFPSAEFSGLITYSVSLVEE
SQDPLIPETLLYKDTVVFRVAPCIFIPCTQVPL----CRELQLQGFVDTVTELSEKSNSQVAS---------
----DEMVFCYTQARHKTTSLILDTPRAPKLDEFPMKYSLSPGIGYMTHDTEDHKVASMDSIGNVMMSPPVK
VQGKEYPLGRVLIGSSFYPSMEGRAMSKTLRDFLYDQQVQAPVELYSDWLMTSHLDEFMCFIPTDEKSEGKK
GFQLLLASPSACYKLFQEKQKEGYGDAFLFDGVRADQLLSNRR---------------------EKCIHLNH
DILKTELGLVEQDIIEIPQLFCLEKLTNVPSDQQPKRLFARPYFPNLLQMIVMGKNLGIPKAFGPQIMGTCC
LEEKMRCLLEPLGFKCTFISDFDCYLADVGDICACATVHRVPFAFKWWSAPPGGRA
>ENSSBOG00000030271/1-692
-VSVEGQAMSFQNIIRLSLDSPAHAVCVIGTEICLDLSRCAPEKCQCFTIRGSGRVVIDVANT--VISEKES
ATIWWPLSDTTYATVKMVSPSPSLDGDKVSVKYYGLNEDAPVGKAVLYLTGIEVSLEADIYRNGQVEVSRDK
QAKKKWTWGPSGWGAILLVNCNPADAGQQLED-KTKRVIFSEEIKNLSQMTLKVQGPSCLLKKYRLVLHTSE
EESKKARVYWPRKDDSSTFELVLGPSQHAYTLAL----LGDHLKETFYLEAMVFPSAEFSGLITYSVSLVEE
SEDPLIPETPLYKDTVVFRVAPCIFIPCTQVPLEVYLCRELQLQGFVDTVTELSDKSNSQVASVYEDPSRLG
RWLQDEMAFCYTQAPHKTTSLILDTPRAPNLDEFPMKYSLSPGIGYMTHDTEDHKVASMDSIGNVMVSPPVK
VQGKEYPLGRVLIGSSFYPSVEGRAMSKTLRDFLYAQQVQAPVELYSDWLMTGHVDEFMCFIPTDEKSEGKK
GFQLLLASPSACYKLFQEKQKEGYGDAFLFDGVRADQLLSNRREAKTIDQLLADESLRKQNEYVEKCIHLNH
DILKTELGLVEPDIIEIPQLFCLEKLTNVPSDQQPKRLFARPYFPNLLQMIVMGKNLGIPKPFGPQIKGTCC
LEERMRCLLEPLGFKCTFISDFDCYLADVGDICACATVRRAPFAFKWWKMVP----
>ENSANAG00000033689/1-633
------QAMSFQNIIRLSLDSPAHAVCVIGTEICLDLSSRGP--CR-------------------ETSEAIT
GNQGYPPRLGTWPRALLLAPSP-----QVSVKYYGLNEDAPVGKAVLYLTGIEVSLEADIYRNGQVEVSRDK
QAKKKWTWGPSGRGAILLVNCNPADASQQLED-KTKRVIFSEEIKNLSQMTLNVQGPSCLLKKYRLVLHTSE
EESKKARVYWPRKDNSSTFELVLGPSQHAYTLAL----LGDHLKETFYLEAMVFPSAEFSGLIAYSVSLVEE
SQDPLIPETPLYKDTVVFRVAPCIFIPCTQVPLEVYLCRELQLQGFVDTVTELSEKSNSQVASVYEDPSRLG
RWLQDEMAFCYTQAPHKTTSLILDTPRAPSLDEFPMKYSLSPGIGYMTHDTEDHKVASMDSIGNVMVSPPVK
VQGKEYPLGRVLIGSSFYPSATGVSHHAQLR------------KLASD-------SSFL-----------SQ
GFQLLLASPGACYKLFQEKQKEGYGDAFLFDGVRADQLLSNRREAKTIDQLLADESLRKQNEYVEKCIHLNH
DILKTELGLVEQDIIEIPQLFCLEKLTNVPSDQQPKRLFARPYFPNLLRMIVMGKNLGIPKPFGPQIKGTCC
LEEKMRCLLEPLGFKCTFISDFDCYLADVGDICACATVRRVPFAFKWWKMVP----
>ENSPSMG00000006237/1-673

```
--------MSFQNIIRLSLDNPAHAVCVLGSEIYVDVSGCAPQMCECFTIRGSGRVLIDIADT--MITDKED
ATIWWPLTEPTYAIVKMTSPSFMVDGDKVSL---GQRDEGCLG---------EVSLEVDIYRNGQVGLPSDR
KAKKKWTWGPNGWGAILLVNCSSADTGQP-EDGKTKTVLLSEEIKNLSQMTLNVQGPSYTLKKYQLILHTSE
EESKKARVYWPQKDSSHTFELVLGPGQHTYTLAL----LGNQLKETLYVEATEFPSASFSGLISYSASLVEE
SQDPAIPETPVYKDTVVFRVAPCIFLPSTQMPLEVYLCRELQLQGFVETVTELSEKSNFQVATVYEDPNRMG
RWLQDEMAFCYTQAPHKTISFVLDTPRAISLEEISMKYSLSPGVGYMIQHTKDQRVASMDSIGNLMVSPPVK
VQGKEYPLGRVLIGSSFYPSTEGRAMSDTLRDFLYAQQVQAPVEVYSDWLMTGHMDEFMCFVPIDDKSDGKK
SFQLLLASPSACYKLFQEKQKQGYGNALLFDEVRGDQLLSNGREAKTIDQFLADENLRKQNEYAEKCILLNR
DILKTELGLVEKDIIDIPQLFCLARLTNVPSDQQTQRPFVRPYFPDLLQMIVIGKTLGIPKPFGPQIKGTCC
LEERVCQLLEPLGFNCTFIDDFDCYLTEVGDLCACANIRRVPFAFKWWKMVP----
>ENSPCOG00000005699/1-657
--------MSFQNIVRLSLDNPTHAVCVLGSEIYVDLSGCAPQLCECFTIRGSGRVLIDIADT--MITDKED
ATIWWPLTEPTYAIVKMTSPSFMVDGDKVLVTYYGTNEEMSLGTAVLFLTG-------------IGECWSH
RGPKKWTWGPNGWGAILLVNCSPADAGQL-EDR-TKKVLLSEEIKNLSQMTLNVQGPSYTLKKYQLVLHTSE
EESKKARVYWPQKDSSHTFELVLGPGQHTYTLAL----LGNHLKETLYVEAMEFPSASFSGLISYSASLVEE
SQDPSIPEIPVYKDTVVFRVAPCIFLPSTQMPLEVYLCRELQLQGFVNTVTELSEKNNFQVASVYEDPNRLG
RWLQDEMAFCYTQAPHKTVSFVLDTPRAISLDEISMKYSLSPGVGYMIQYTKDHRVASMDSIGNLMVSPPVK
VQGKEYPLGRVLIGSSFYPSAEGRDMSDTLRDFLYAQQVQAPVELYSDWLMSGHMDEFMCFVPIDDKSDGKK
VCGGLGAGPS----LPSSGQHRPTGLP---------ALLPTGREAKTIDQFLADENLRKQNEYAEKCILLNR
DILKTELGLVEKDIIDIPQLFCLARLTNVPSDQQPQRPFARPYFPDLLQMIVMGKTLGIPKPFGPQIKGTCC
LEEKVCHLLEPLGFKCTFIDDFDCYLTDVGDLCACANIHRVPFAFKWWKMVP----
>ENSMICG00000014075/1-677
--------MSFQNIIRLSLDDPAHAVCVLGSEIYVDVSGCAPQMCECFTIHGSGRVLIDIADT--MITDKED
ATIWWPLTEPTYAIVKMTSPSFMVDGDKVLVTYYGTNEEMSLGTAVLYLTGIDVSLEVDIYRTGQVGLPSDK
KAKKKWTWGPNGWGAILLVNCSP-DPGQL-EDQ-TKKVLLSDEIKNLSQMMLNVQGPSYTLKKYQLVLHTSE
EESKKARVYWPQKDSAHTFELVLGPGQHTYTLPL----LGNHLKETLYVEATEFPSASFSGLISYSASLVEE
SQDPLIPEVPVYKDTVVFRVAPCIFLPSTQMPLEVYLCRELQLQGFVNTVTELSEENHFQVASVYEDPNRLG
KWLQDEMAFCYTQAPHKTVSFVLDTPRAILLEEISMKYSLSPGVGYMIQETKDHTVASMDSIGNLMVSPPVK
VQGKEYPLGRVLIGSRFYPSAEGRAMSEALRDFLYAQQVQAPVELYADWLMSGHMDEFMCFVPIDDKSDGKK
SFQLLLASPRACYTLLQEKQRQGYGNALLFDEVRGDQLLSNGREAASL--FLTSPSLPA----PQKCILLNR
EILKTELGLVEKDIIDIPQLFCLARLTNVPSEQQPQRPLARPYFPDLVRMIVMGKTLGIPKPFGPQIKGTCC
LEEKVCQLLEPLGFKCTFIDDFDCYLTDVGDLCACANIRRVPFAFKWWKTVP----
>ENSOGAG00000014538/1-683
--------MLFQNIVHLSLDNPAHAVCVMGSEICVDISEGAPRECASFTISGSAGILIDIVDS--MITEKED
TTIWWPLSDPTYAVMKMTSPSPAVDGDKVLVTYYGPHEN-TLGTAVLYLTGIAVSLEVDIYRNGQIVTPEDR
KSKKKWIWGPNGCGAILLVNCSPAEIGQL-EN-KTKKVFFPEEMKNLSLMTVTIQGPSSLLRRYQLVLHTSE
EESKKARVYWPHKDGSGTFDMVLGPGQHTYTLAL----LGNNLKETFYVEATEFPSASFSGLISYSASLVEE
SQDLSIPETPVYKDTVVFRVAPCIFIPSTQMPLEIYLCRELQLQGFVDTVVELSERSNFQMASVYEDPNRLG
KWLQNEMAFCYTQAPHKTTSLVLDTPRATDLEDFPMKYSLSPGVGYVIYYTQDQRVASMDSIGNLTVSPPVK
VQGKEYPLGRILIGGSFYPSEEGRAMSNTLRDFLCAQQVQAPVELYSDWLMSGHMDEFMCFVPTDDKSEGRK
GFQLLLASPRACYKLFQEKQRQGYGNALLFNEVRQDQLLSNGREFKTIDQLLADENLRKQNEYVEKCIQLNR
DILKTELGLVEKDIIDIPQLFCLAQLANVPPEQQIRKHFAKPYFPDLLQMVVLGKNLGIPKPFGPQIKGTCC
LEEHVRHLLEPLGLHCTFIDDFDRYLTDIGDLCACANIRRVPFAFKWWKMVP----
>ENSUMAG00000021304/1-683
--------MSFQSLVHLSLDSPVHALCVLGMEICLDLNGCAPEKCKSFTVSGSPGVLVDILRTVPAVSREEA
APTRWPLSHPIDVLVKMHSPSSAIDSDKVLVSYYLPDEDVPVATAVLYVTGVMVSLDVDIYRSGQVEMASDK
QAKKKWVWGPSGWGAILLVNCSPADKGQI-VDKKTTKVFFPEEIKSLSQMTLNVQGPGDTLKKYRLVLHTSK
EEAEKARVYRPQRNSSSTFEMLLGPGRHAYTFAP----LEDPLKETFYVEAVEFPSADFSGLISYSVSLVEE
SQDPSIPETLVYKDTVVFRVAPCIFTPSTQMPLEVYLCRELQVQGFVNTIMELSEKSNSQVASVYEDPNRLG
RWLQDEMAFCYTQAPHKTLSLVLDTPRVLMLDDIPMKYSLSPGVGYMIQGTKDHRVASMDSIGNLMVSPPVQ
VED--------IIGSCFYPSKEGRDMGKALRDFLYAQRVQAPVELFSDWLMVGHVDEFMCFVPTQDKSQGGK
GFRLLLASPSSCYRLFEEKQKEGYGDMTLFEEVQEDQLLSTGREANSISQLLADENMRKQNNYVEKCVNLNR
AILKKELGLVERDIIDLPQLFCLEQLTNVPSNEQMAKLFARPYFPNLLQMIVMDKHLGIPKPFGPQVKGTCC
LEEKICQLLEPLGFQCTFIKDFDCYLTEIGDFCACANVRRVPFAFKWWRMVPEPQA
>ENSUAMG00000014926/1-691
--------MSFQSLVHLSLDSPVHALCVLGMEICLDLNGCAPEKCKSFTVSGSPGVLVDILRTVPAVSREEA
APTRWPLQHPIDVLVKMTSPSSAIDSDKVLVSYYLPDEDVPVATAVLYLTGVMVSLDVDIYRSGQVEMASDK
QAKKKWVWGPSGWGAILLVNCSPADKGQI-VDKKTTKVFFPEEIKSLSQMTLNVQGPGNTLKKYRLVLHTSK
EEAEKARVYRPQRNSSSTFEMLLGPGRHAYTFAP----LEDPLKETFYVEAVEFPSADFSGLISYSVSLVEE
SQDPSIPETLVYKDTVVFRVAPCIFTPSTQMPLEVYLCRELQVQGFVNTIMELSEKSNSQVASVYEDPNRLG
RWLQDEMAFCYTQAPHKTLSLVLDTPRVLMLDDIPMKYSLSPGVGYMIQGTKDHRVASMDSIGNLMVSPPVQ
VEGKEYPLGRILIGSCFYPSKEGRDMGKALRDFLYAQRVQAPVELFSDWLMVGHVDEFMCFVPTQDKSQGGK
GFRLLLASPSSCYRLFEEKQKEGYGDMTLFEEVQEDQLLSTGREANSISQLLADENMRKQNNYVEKCINLNR
```

AILKKELGLVERDIIDLPQLFCLEQLTNVPSNEQMAKLLARPYFPNLLQMIVMDKHLGIPKPFGPQVKGTCC
LEEKICQLLEPLGFQCTFIEDFDCYLTEIGDFCACANVRRVPFAFKWWRMVPEPQA
>ENSAMEG00000000749/1-698
VVSLEGRVMSFQSLVHLSLDSPVHALCVLGMETCLDLNGCAPEKCKSFTVSGSPGVLVDILDTVPAVSREEA
APTRWPLSHPIDVLVKMTSPSSAIDSDKVLVSYYLPDEDVPVATAVLHLTGVMVSLDVDIYRSGQVEMASDK
QAKKKWVWGPSGWGAILLVNCSPADKGQI-VD-KTTKVFFPEEIKSLSQMTLNVQGPGDTLKKYRLVLHTSK
EEAEKARVYWPQRNSSSTFEMLLGPGRHAYTFAP----LEDPLKETFYVEAVEFPSADFSGLISYSVSLVEE
SQDPSVPETLVYKDTVVFRVAPCIFTPSTQMPLEVYLCRELQVQGFVNTVMELSEKSNSQVASVYEDPNRLG
RWLQDEMAFCYTQAPHKTLSLVLDTPRVLMLDDIPMKYSLSPGVGYMIQGTKDHRVASMDSIGNLMVSPPVQ
VEGKEYPLGRILTGSCFYPSKEGRDMGKALRDFLYAQRVQAPVELFSDWLMVGHVNEFMCFVPTQDKSQGDK
GFRLLLASPSACYRLFEKKQKEGYGDMTLFEEVQEDQLLSTGREANSISQLLADENMRKQNDYVEKCINLNR
AILKKELGLVERDIIDLPQLFCLEQLTNVPSNEQMAKLFARPYFPNLLQMIVMGKHLGIPKPFGPQVKGTCC
LEEEICQLLEPLGFQCSFIEDFDCYLTEIGDFCACANVRRVPFAFKWWRMVPEPQA
>ENSNVIG00000014111/1-696
VDGLEGRAMSFQSIVRLSLDSPVHALCVLGTEICLDINGCAPEKCKSFTVSGSPGVLVDVPNTVQAGSGEEA
APTRWPLSHPIDVLVKMTAPSSAIDGDKVSVSYYLSDEDVPVATAVLCLTGIVVSLDVDVYRTGQVQMTSDK
QAKKNWVWGPSGWGAILLVNCSPADKGQI-VDKKTTKVFLPEEIQSLSQMTLNVQGPSCALKKYRLLLHTSK
EEAEKARVYRPQRDSSSTFELLLGPGRSTYSLTP----LEKPLKETFYVEAVEFPSASFSGLISYSVSLLEE
SQDPSIPETLAYKDTVVFRVAPCVFTPSTQMPLEVYLCRELQVEGFVNTVMELSEKSNIQVASVYEDPNRLG
RWLQDEMAFCYTQAPHKTVSLVLDTPRVLTLDDFPMKYSLSPGVGYLIQRTKDHRVASMDSIGNLMVSPPVK
VEGKEYPLGRILIGSSFYPSKEGRDMGKALRDFLYAQRVQAPVELFSDWLMAGHVDEFLCFIPTRDKSEGDK
GFRLLLASPSSCYRLFEEKKKEGYGDMTLLEEVREDQLLSNGREASTINQLLMDGNLRKQNNYVEKCINLNR
DLLKKELGLVEKDIIDIPQLFCLEQLTNVPSNEQTAKLFARPYFPNLLQMIVMDKNLGIPKPFGPQIKGICC
LEEKIRQRLEPLGFQCTFIDDFDCYLTEIGDFCACANIRRVPFAFKWWNMVPE---
>ENSMPUG00000016211/1-695
VDGLEGRAMSFQSIVRLSLDSPVHALCVLGTDICLDINGCAPEKCKSFTVSGSPGVLVDVPNTVQAGSGEEA
APTRWPLSHPIDVLVKMTAPSSAINGDKVSVSYYLSDEDVPVATAVLCLTGIVVSLDVDIYRTGQVQMTSDK
QAKKNWVWGPSGWGAILLVNCSPADKGQI-VDEKTTKVFLPEEIQSLSQMTLNVQGPSCALKKYRLLLHTSE
EEAEKARVYRPQRDSSSTFELLLGPGRSTYTLTP----LESPLKETLYVEAVEFPSASFSGLVSYSVSLLEE
SQDPSIPETLAYKDTVVFRVAPCVFTPSTQMPLEVYLCRELQVEGFANTVMELSEKSNIQVASVYEDPNRLG
RWLQDEMAFCYTQAPHKTVSLVLDTPRVLTLDDFPMKYSLSPGVGYVIQHTKDHRVASMDSIGNLMVSPPVK
AEGKEYPLGRILIGSSFYPSKEGRDMGKALRDFLYAQRVQAPVELFSDWLMAGHVDEFLCFIPTQDKSEGDK
GFRLLLASPSSCYRLFEEKKKEGYGDVTLLEEVREDQLLSNGREASTINQLLTDENLRKQNNYVEKCINLNR
DILKEELGLGEEDIVHIPQLFCLEQLTNVPSNEQTAKLFARPYFPNLLQMIVMDKNLGIPKPFGPQIKGICC
LEEKIRQLLEPLGFQCTFIEDFDCYLTEIGDFCACANIRRVPFAFKWWNMVP----
>ENSCAFG00845003990/1-691
--------MSFQSIIHLSLDSPVRALCVLGMEICLDVNGCTPERCEAFTISSSPGVSVDLHGTLPGTGQEGT
AVTRWPLSNPTDVLVKMTCPSSASDGDQVLVSYYLPDEDAPVATAVLRLTGIVVSLDVDIYRSGQVEIASDK
QAKKNWVWGPSGWGAILLVNCSPADKGQI-IDQKTTKVFFPEEIKSLSQMTLNVQGPGATLKKYRLVLHTSE
EEARKARVYRPQRDSSSTFEVLLGPGQCTYTFAP----LENNLKETFYVEAIEFPSADFSGLISYSVSLVEE
PQDPSIPETLVYKDTVVFRVAPCVFIPSTQMPLEVYLCRELQVQGFVNTVMELSEKSNSQVASVYEDPNRLG
RWLQDEMAFCYTQAPHKTLSLVLDTPRVLTLEDFPMKYSLSPGVGYVIQCTKDHRVASMDSIGNLMVSPPVK
VEGKEYPLGRILIGSCFYPSKEGRDMSKALRDFLYAQRVQAPVELFSDWLMVGHMDEFMCFIPTQDKSEGEK
GFRLLLASPSSCYRLFEEKQKEGYGDMALFEEVREDQLLSNGRQANTINQLLADKNMRKQNDYVEKCINLNR
DILKKELGLVERDIIDIPQLFCLEQLTNVPSSEQTGKFFARPYFPDLLQMIVMGKNLGIPKPFGPQIKGTCC
LEEKICQLLEPLGFKCTFIDDFDCYLTEIGDFCACANIRRVPFAFKWWRMVPEPQA
>ENSCAFG00020005177/1-691
--------MSFQSIIHLSLDSPVRALCVLGMEICLDVNGCTPERCEAFTISSSPGVSVDLHGTLPGTGQEGT
AVTRWPLSNPTDVLVKMTCPSSASDGDQVLVSYYLPDEDAPVATAVLRLTGIVVSLDVDIYRSGQVEIASDK
QAKKNWVWGPSGWGAILLVNCSPADKGQI-IDQKTTKVFFPEEIKSLSQMTLNVQGPGATLKKYRLVLHTSE
EEARKARVYRPQRDSSSTFEVLLGPGQCTYTFAP----LENNLKETFYVEAIEFPSADFSGLISYSVSLVEE
PQDPSIPETLVYKDTVVFRVAPCVFIPSTQMPLEVYLCRELQVQGFVNTVMELSEKSNSQVASVYEDPNRLG
RWLQDEMAFCYTQAPHKTLSLVLDTPRVLTLEDFPMKYSLSPGVGYVIQCTKDHRVASMDSIGNLMVSPPVK
VEGKEYPLGRILIGSCFYPSKEGRDMSKALRDFLYAQRVQAPVELFSDWLMVGHMDEFMCFIPTQDKSEGEK
GFRLLLASPSSCYRLFEEKQKEGYGDMALFEEVREDQLLSNGRQANTINQLLADKNMRKQNDYVEKCINLNR
DILKKELGLVERDIIDIPQLFCLEQLTNVPSSEQTGKFFARPYFPDLLQMIVMGKNLGIPKPFGPQIKGTCC
LEEKICQLLEPLGFKCTFIDDFDCYLTEIGDFCACANIRRVPFAFKWWRMVPEPQA
>ENSVVUG00000029038/1-681
--------MSFQSIIHLSLDSPVRALCVLGMEICLDVNGCAPEKCEAFTISSSPGVSVDLHSTLPGPGREGT
TMTRWPLSNPTDVLVKMTCPSSANDGDQV----YLPSDDLPIEGLWPTLLA-VVSLDVDIYRSGQVEIASDK
QAKKNWVWGPSGWGAILLVNCSPADKGQI-IDQKTTKVFFPEEIKSLSQMTLNVQGPGATLKKYRLVLHTSE
EEARKARVYRPQRDSSSTFEVLLGPGQCTYTFAP----LENNLKETFYAEAIEFPSADFSGLISYSVSLVEE
PQDPSIPETLVYKDTVVFRVAPCVFIPSTQMPLEVYLCRELQVQGFVNTVMELSEKSNSQVASVYEDPNRLG

RWLQDEMAFCYTQAPHKTLSLVLDTPRVLTLEDFPMKYSLVWG-----SCTKDHRVASMDSIGNLMVSPPVK
VEGKEYPLGRILIGSCFYPSKEGRDMSKALRDFLYAQRVQAPVELFSDWLMVGHMDEFMCFIPTQDKSEGEK
GFRLLLASPSSCYRLFEEKQKEGYGDMALFEEVREDQLLSNGREANTINQLLADENMRKQNDYVEKCINLNR
DILKKELGLVERDIIDIPQLFCLEQLTNVPSSEQTGKFFARPYFPDLLQMIVMGKNLGIPKPFGPQIKGTCC
LEEKICQLLEPLGFKCTFIDDFDCYLTEIGDFCACANIRRVPFAFKWWRMVPEPQA
>ENSPPRG00000008467/1-691
--------MSFQSLVHLSLDGPVHALCVLGVDICLDLSGCAPEKCKSFTISGSPGVLVDIHNTSPLMTKEET
ATTRWPLSDPMDVLVNMISPSSAPDGDKVLVSYYLPDEEVPVATAVLCLTGIVVSLDVDIYRSGQVEVASDK
QAKKNWVWGPSGWGAILLVNCSPTDKGQS-MDKKTTRVFFPEEIKSLSQMTLNVQGPSCTLKKYRLVLHTSK
EEAEKARVYRPQKDSSSTFEVLLGPGQHTYAFAP----LENHLKETFYVEAVEFPSADFSGLISYSVSLVQE
SPDPSIPETLVYKDTVVFRVAPCVFIPSTQMPLEVYLCRELQVEGFVNTMMELSEKSNSQVASVYEDPNRLG
RWLQDEMAFCYTQAPHKTLSFVLDTPRALTLEDFPMKYSLSPGVGYVIQCTQDHRVASMDSIGNLMVSPPVK
VGGKEYPLGRILIGSCFYPSKEGRDMSKALRDFLYAQRVQAPVELFSDWLMVGHIDEFMRFIPTQDKSEGEK
GFRLLLASPSSCYRLFEEKQREGYGDVTLFEEVRGDQLLSNGREANTIHQLLADENMRKQNEYVEKCINLNR
DIVKKELGLAERDIIDIPQLFCLEQLTNVPSDQQTGKFYARPYFPDLLQMMVMGKNLGIPKPFGPQIKGTCC
LEKKICQLLEPLGFQCTFIDDFDCYLTEVGDFCACANIRRVPFAFKWWRMVPEPQA
>ENSPLOG00000016269/1-691
--------MSFQSLVHLSLDGPVHALCVLGVDICLDLSGCAPEKCKSFTIRGSPGVLVDIHNTSPLMTKEET
ATTRWPLSDPMDVLVNMISPSSAPDGDKVLVSYYLPDEEVPVATAVLCLTGIVVSLDVDIYRSGQVEVASDK
QAKKNWVWGPSGWGAILLVNCSPTDKGQS-MDKKTTRVFFPEEIKSLSQMTLNVQGPSCTLKKYRLVLHTSK
EEAEKARVYRPQKDSSSTFEVLLGPGQHTYAFAP----LENHLKETFYVEAVEFPSADFSGLISYSVSLVQE
SPDPSIPETLVYKDTVVFRVAPCVFIPSTQMPLEVYLCRELQVEGFVNTMMELSEKSNSQVASVYEDPNRLG
RWLQDEMAFCYTQAPHKTLSFVLDTPRALTLEDFPMKYSLSPGVGYVIQCTQDHRVASMDSIGNLMVSPPVK
VGGKEYPLGRILIGSCFYPSKEGRDMSKALRDFLYAQRVQAPVELFSDWLMVGHVDEFMCFIPTQDKSEGEK
GFRLLLASPSSCYRLFEEKQREGYGDVTLFEEVRGDQLLSNGREANTIHQLLADENMRKQNEYVEKCINLNR
DIVKKELGLAERDIIDIPQLFCLEQLTNVPSDQQTGKFYARPYFPDLLQMMVMGKNLGIPKPFGPQIKGTCC
LEKKICQLLEPLGFRCTFIDDFDCYLTEVGDFCACANIRRVPFAFKWWRMVPEPQA
>ENSPTIG00000014335/1-691
--------MSFQSLVHLSLDGPVHALCVLGVDICLDLSGCAPEKCKSFTISGSPGVLVDIHNTSPLMTKEET
ATTRWPLSDPMDVLVNMISPSSAPDGDKVLVSYYLPDEEVPVATAVLCLTGIVVSLDVDIYRSGQVEVASDK
QAKKNWVWGPSGWGAILLVNCSPTDKGQS-MDKKTTRVFFPEEIKSLSQMTLNVQGPSCTLKKYRLVLHTSK
EEAEKARVYRPQKDSSSTFEVLLGPGQHTYTFAP----LENYLKETFYVEAAEFPSADFSGLISYSVSLVQE
SPDPSIPETLVYKDTVVFRVAPCVFIPSTQMPLEVYLCRELQVEGFVNTVMELSEKSNSQVASVYEDPNRLG
RWLQDEMAFCYTQAPHKTLSFVLDTPRALTLEDFPMKYSLSPGVGYVIQCTQDHRVASMDSIGNLMVSPPVK
VGGKEYPLGRILIGSCFYPSKEGRDMSKALRDFLYAQRVQAPVELFSDWLMVGHIDEFMCFIPTQDKSEGEK
GFRLLLASPSSCYRLFEEKQREGYGDVTLFEEVREDQLLSNGREANTIHQLLADENMRKQNEYVEKCINLNR
DIVKKELGLAERDIIDIPQLFCLEQLMNVPSDQQTGKFYARPYFPDLLQMMVMGKNLGIPKPFGPQIKGTCC
LEKKICQLLEPLGFQCTFIDDFDCYLTEVGDFCACANIRRVPFAFKWWRMVPEPQA
>ENSFCAG00000010186/1-691
--------MSFQSLVHLSLDGPVHALCVLGVDICLDLSGCAPEKCKSFTISGSPGVLVDIHNTSPLMTKEEM
ATTRWPLSDPMDVLVNMVSPSSAPDGDKVLVSYYLPDEEVPVATAVLCLTGIVVSLDVDIYRSGQVEVASDK
QAKKNWVWGPSGWGAILLVNCSPTDKGQS-MDKKTTKVFFPEEIKSLSQMTLNVQGPSCTLKKYRLVLHTSK
EEAEKARVYRPQKDSSSTFEVLLGPGQHTYTFAP----LENHLKETFYVEAVEFPSADFSGLISYSVSLVQE
SPDPSIPETLVYRDTVVFRVAPCVFVPSTQMPLEVYLCRELQVQGFVNTVMELSEKSNSQVASVYEDPNRLG
RWLQDEMAFCYTQAPHKTLSFVLDTPRALTLEDFPMKYSLSPGVGYVIQCTQDHRVASMDSIGNLMVSPPVK
VGGKEYPLGRVLIGSCFYPSKEGRDMSKALRDFLYAQRVQAPVELFSDWLMVGHIDEFMCFIPTQDKSEGEK
GFRLLLASPSACYRLFEEKQREGYGDVTLFEEVREDQLLSNGREANTIHQLLADENMRKQNEYVEKCINLNR
DIVKKELGLAERDIIDIPQLFCLEQLTNVPSDQQTGKFFARPYFPDLLQMMVMGKNLGIPKPFGPQIKGTCC
LEKKVCQLLEPLGFKCTFIDDFDCYLTEVGDFCACANIRRVPFAFKWWRMVPEPQA
>ENSEASG00005015759/1-687
--------MSFQSIIHLSLDSPVHALCVLGMEIYLDLRSCAPQKCKSFTVSSSLGVLVDVHNSVPVMTREEV
ATTLWPLSDSMDVLVKMLSPSSAIDGDKVLVSYHQPDEEVPVATATLYLTGIAVSLDVDVYRRGQVEMASDN
RAKRNWVWGPSGWGAILLVNCSPADMGQL-TDRNTTKAFFPEEMKNLSQMTLSVQGPNCILKKHRLVLHTSE
DEAQRARVYQPQKDGSSTLELVVGPGRHTYTLPP----LESPLKETFYVEAVEFPSASFSGLIAYSVCLVEE
PPDPSIPETLVYKDTVVFRVAPCIFIPSTQMPLEVYLCRELQVQGFVNTVMELSEKSNSQVASVYEDPSRLG
RWLQDEMAFCYTQAPHKTISLVLDTPRVLKLEDFPMKYSLSPAVGYRIQRTEDHRVASMDSIGNLTVSPPVT
VGGKEYPLGRILIGSSFYPSKEGRDMGKALRDFLYAQQVQAPVELFSDWLMVGHIDEFMCFIPTQDKREGDK
GFRLLLASPSSCYQLFQEKQKEGYGDMHLFEGVRADQLLSNGREANTIHQLLADENMRRQNDYVQKCINLNR
DVLKAELGLAERDIIDIPQLFCLEQLTNAPSDQQTGKLFARPYFPNLLQTIVMGKNLGIPKPFGPQIKGTCC
LEEKICQLLEPLGFQCTFIDDFDCYLTEIGDFRACANVRRVPFAFKWWRMVP----
>ENSCDRG00005005429/1-695
VVSLEGCAMSFRSLVHLSLDSPAHAICVVGMEICLDLSGCAPPSCESFTISGSLGVLVDIHNTVPGKAKEEA
AMARWPLSHPVDVLVKMTSPSSAINEEKILVSYYQCNEKVPVATALLYLTGIEISLDVDIYRSGHVEMASDK

QAKKNWVWGPSGWGAILLVNCSPADMGQL-VDKKSGNVLFMEEIKGLSQMTLKVEGPSCVLKKYRLVLHTSK
EESEKARVYRPKKDSSSSFELVLGPDWHTYTFAP----LEKNLMETFYIEAVEFPSASFSGLISYSVSLVEE
SQDLLIPETLLYKDTVVFRVAPCIFIPTTQMPLEVYLCRELQVQGFVNTVTLLSERSNSQVASVYEDPNRLG
RWLQDEMAFCYTQAPHKTVSLVLDTPRALKLEDFPMKYSLSPGIGYIIQRTKDHRVASMDSIGNLMVSPPVK
VEGKEYPLGRILIGGSFYPSKEGRDMSKALRDFLYAQQVQAPVELFSDWLMAGHICEFMCFIPAQYKQEGRK
DFRLLLASPSSCYKLFKEKQKEGYGDMPLFEEVRKDQLISNGREASTINQLLADENMRKQNDYVENCINLNR
DILKRELGLVEKDIIDIPQLFCLEQLTNVPSSEQTQKLFARPYFPDLLQIIVMGTNLGIPKPFGPRINGTCC
LEEKVCQLLEPLGFQCTFIDDFDCYLTEIGDFCSCANIRRVPFAFKWWRMVP----
>ENSRFEG00010015258/1-693
-VSLEGRAMSFQSLIHLSLDNPERAHCIVGTEICLDLSGCAPQKCKFFTISGSPGVFVNIHNTGPVITSEEA
AVNRWSLSDPMDVLVKMTSPSTVIDEEKVLVSYYQPDEEVPVATAELYLTGVAVSLDVDIYRNGQVEMANNK
QAKKNWVWGPKGWGAILLVNCSPTDMGQL-IDKKTTRVIFSEEIKSLSQMTLNVQGPSCILKNHQLVLYTSE
EESEKTRVYRPQEDGSSTFELVLGPGRHTYTFAP----LESHLKETFYVEALEFPSADFFGLISYSVSLVEE
SQDPSIPETLVYKDTVVFRVAPCIFTPSTQMPLEVYLCRELQVQGFVSAVVELSEKSNSQVASVYEDPSRLG
RWLQDEMAFCYTQAPHKTISLVLDTPRVLKLEDFPMKYSLSPGVGYMIQCTKDHRVASMDSIGNLMVSPPVK
VEGKEYPLGRILFGSSFYPSKEGRAMSKALRDFLYAQQVQAPVELFSDWLMTGHVDEFMCFIPTHDKSEGKK
-FRLLLASPSACYKLFQEKQKEGYGDMPLFEEVREDQLLSNGREANTINQLLADENMRKQNAYVEKCVNLNR
DILKRELGLVEKDIIDVPQLFCLEQLTNVPSSQQTEKLYARPYFPNLLQVIVMGKNLGIPKPFGPQIKGTCC
LEEKICQLLEPLGFKCTFINDFDCYLTEVGDFCACANIRRVPFAFKWWRMVP----
>ENSDLEG00000017509/1-676
MGDLEGRAMSFQSTVHLSLDSPAHAICVQGMEIYLDVNGCAPQQCESFTVDSSPGVVVQIYGTDPVKIGEEA
AMNWWPLSHPTNVLVGMTSPSSANDKGKVSVSYYQHNEDVPMATAVLYLTSIGEAL----------------
--PKNWVWGPKGWGAILLVNCSPADAHQL-VD-RKSKVFSTEETKSLSQMILRVQGPSYILKKCRVVLHTSK
EESEKARVYRPQKDSSSTFELVLGPDQHTYNLAP----VEDDLEETFYVEALEFPSASFSGLISYSASLVEE
SQDLSIPETVVYKDTVVFRVAPCVFVSSTQMPLEVYLCRELQVQGFVNTVMELSERSNIQVASVYEDPNRLG
RWLQDEMAFCYTQAPHKTTSLVLDTPRLPKLDDFPMKYSLSPGVGYMTQRTQDHTVASIDSIGNLMVSPPVK
AQGKEYPLGRILIGSSFYPSKDCRNLSKTLRDFLYAQQVQAPVELFSDWLMIGHAYEFMCFIPAQYKVEDKK
DFWLLLASPSSCYKLFKEKQKEGYGDARLFEGIRKDQLLSNGREANTINQLLADENMRKQNAYVEKCINLNR
SILKRELGLEEQDIIDIPQLFCLEHIANIPSSEQTEKLYARPYFPDLLQMVVMGQNLGIPKPFGPQINGACC
LEEKIRQLLEPLGFQCTFVNDFDCYLTEIGDFCSCANIRRVPFAFKWWRMVP----
>ENSDLEG00000017924/1-676
MGDLEGRAMSFQSTVHLSLDSPAHAICVQGMEIYLDVNGCAPQQCESFTVDSSPGVVVQIYGTDPVKIGEEA
AMNWWPLSHPTNVLVGMTSPSSANDKGKVSVSYYQHNEDVPMATAVLYLTSIGEAL----------------
--PKNWVWGPKGWGAILLVNCSPADAHQL-VD-RKSKVFSTEETKSLSQMILRVQGPSYILKKCRVVLHTSK
EESEKARVYRPQKDSSSTFELVLGPDQHTYNLAP----VEDDLEETFYVEALEFPSASFSGLISYSASLVEE
SQDLSIPETVVYKDTVVFRVAPCVFVSSTQMPLEVYLCRELQVQGFVNTVMELSERSNIQVASVYEDPNRLG
RWLQDEMAFCYTQAPHKTTSLVLDTPRLPKLDDFPMKYSLSPGVGYMTQRTQDHTVASIDSIGNLMVSPPVK
AQGKEYPLGRILIGSSFYPSKDCRNLSKTLRDFLYAQQVQAPVELFSDWLMIGHAYEFMCFIPAQYKVEDKK
DFWLLLASPSSCYKLFKEKQKEGYGDARLFEGIRKDQLLSNGREANTINQLLADENMRKQNAYVEKCINLNR
SILKRELGLEEQDIIDIPQLFCLEHIANIPSSEQTEKLYARPYFPDLLQMVVMGQNLGIPKPFGPQINGACC
LEEKIRQLLEPLGFQCTFVNDFDCYLTEIGDFCSCANIRRVPFAFKWWRMVP----
>ENSDLEG00000009237/1-676
MGDLEGRAMSFQSTVHLSLDSPAHAICVQGMEIYLDVNGCAPQQCESFTVDSSPGVVVQIYGTDPVKIGEEA
AMNWWPLSHPTNVLVGMTSPSSANDKGKVSVSYYQHNEDVPMATAVLYLTSIGEAL----------------
--PKNWVWGPKGWGAILLVNCSPADAHQL-VD-RKSKVFSTEETKSLSQMILRVQGPSYILKKCRVVLHTSK
EESEKARVYRPQKDSSSTFELVLGPDQHTYNLAP----VEDDLEETFYVEALEFPSASFSGLISYSASLVEE
SQDLSIPETVVYKDTVVFRVAPCVFVSSTQMPLEVYLCRELQVQGFVNTVMELSERSNIQVASVYEDPNRLG
RWLQDEMAFCYTQAPHKTTSLVLDTPRLPKLDDFPMKYSLSPGVGYMTQRTQDHTVASIDSIGNLMVSPPVK
AQGKEYPLGRILIGSSFYPSKDCRNLSKTLRDFLYAQQVQAPVELFSDWLMIGHAYEFMCFIPAQYKVEDKK
DFWLLLASPSSCYKLFKEKQKEGYGDARLFEGIRKDQLLSNGREANTINQLLADENMRKQNAYVEKCINLNR
SILKRELGLEEQDIIDIPQLFCLEHIANIPSSEQTEKLYARPYFPDLLQMVVMGQNLGIPKPFGPQINGACC
LEEKIRQLLEPLGFQCTFVNDFDCYLTEIGDFCSCANIRRVPFAFKWWRMVP----
>ENSMMNG00015008836/1-680
MGDLEGRAMSFQSTVHLSLDSPAHAICVQGVEIYLDVNGCAPQQCESFTVDSSPGVVVQIYGTDPVKIGEEA
AMNWWPLSHPMNVLVGMTSPSSDPD-----TSLRQ-----PW---VLTYSSKAASLDVDVYRSGHFE-TNDK
QAKKNWVWGPKGWGAILLVNCSPADAHQL-VD-RKSKVFSTEETKSLSQMILRVQGPSCILKKCRVVLHTSK
EESEKARVYRPQKDSSSTFELVLGPDQHTYNLAP----VEDDLEETFYVEALEFPSASFSGLISYSASLVEE
SQDLSIPETVVYKDTVVFRVAPCVFVSSTQMPLEVYLCRELQVQGFVNTVMELSERSNIQVASVYEDPNRLG
RWLQDEMAFCYTQAPHKTTSLVLDTPRLPKLDDFPMKYSLSPGVGYMTQRTQDHTVASIDSIGNLMVSPPVK
AQGKEYPLGRILIGSSFYPSKDCRNLSKTLRDFLYAQQVQAPVELFSDWLMIGHAYEFMCFIPAQYKVEDKK
DFWLLLASPSSCYKLFKEKQKEGYGDARLFEGIRKDQLLSNGREANTINQLLADENMRKQNAYVEKCIDLNR
SILKRELGLEEQDIIDIPQLFCLEHIANIPSSEQTEKLYARPYFPDLLQMVVMGQNLGIPKPFGPQINGACC
LEEKIRQLLEPLGFQCTFINDFDCYLTEIGDFCSCANIRRVPFAFKWWRMVP----

>ENSPSNG00000000724/1-676
MGDLEGRAMSFQSTVHLSLESPAHAICVQGMEIYLDVNGCAPQQCESFTVDSSPGVVVQIYGTDPVKIGEEA
AMNWWPLSRPTNVLVGMTSPSSANDKGKVSVSYYQHNEDVPMATAVLYLTSIGESL----------------
--QKNWVWGPKGWGAILLVNCCPADAHQL-VD-RKSKVFSTEETKSLSQMILRVQGPSCILKKCRVVLHTSK
EESEKARVYRPQKDCSSTFELVLGPDQHTYNLAP----VEDDLEETFYVEALEFPSASFSGLISYSASLVEE
SQDLSIPETVVYKDTVVFRVAPCVFVSSTQMPLEVYLCRELQVQGFVNTVMELSERSNIQVASVYEDPNRLG
RWLQDEMAFCYTQAPHKTISLVLDTPRLPKLDDFPMKYSLSPGVGYMTQRTQDHTVASIDSIGNLMVSPPVK
AQGKEYPLGRILIGSSFYPSKDCRNLSKTLRDFLYAQQVQAPVELFSDWLMIGHAYEFMCFIPAQYKVEDKK
DFWLLLASPSSCYKLFKEKQKEGYGDARLFEGIRKDQLLSNGREANTINQLLADENMRKQNAYVEKCIDLNR
SILKRELGLEEQDIIDIPQLFCLEHIANIPSSEQTEKLYARPYFPDLLQMVVMGQNLGIPKPFGPQINGACC
LEEKIRQLLEPLGFQCTFINDFDCYLTEIGDFCSCANIRRVPFAFKWWRMVP----
>ENSPCTG00005022042/1-643
MGDLEGRAMSFQSIVHLSLDSPAHAICVQGVEIYLDVNGCAPQQCESFTVDSSPGVVIQIYGTDPVKIGEEA
AMNWWPLSHPTNVLVGMTSPSSANDKGKVSVSYYQHNEDVPMATAVLYLTSIGESL----------------
--QKNWVWGPKGWGAILLVNCSPADTHQL-VD-RKSKVFSTEETKSLSQMMLRVQGPSCILKKCRVVLHTSK
EESEKARVYRPQKDSSSTFELVLGPDQHTYNLAP----VGDDLEETFYVEALEFPSASFSGLISYSASLVEE
SQDLSIPETVVYKDTVVFRVAPCVFVPSTQ--------------------------------ILPGPGRFC
NWLQDEMAFCYTQAPHKTISLVLDTPRLPKLDDFPMKYSLSPGVGYMTQRTQDHAVASIDSIGNLMVSPPVK
AQGKEYPLGRILIGSSFYPSKDCRNLSKTLRDFLYAQQVQAPVELFSDWLMIGHAYEFMCFIPAQYKVEDEK
GFRLLLASPSSCYKLFKEKQKEGYGDARLFEGIRKDQLLSNGREANTINQLLADENMRKQNSYVEKCIDLNR
NILKRELGLEEQDIIDIPQLFCLEHIANIPSSEQTEKLYARPYFPDLLQMIVMGQNLGIPKPFGPQINGTCC
LEEKIRQLLEPLGVQCTFINDFDCYLTEIGDFCSCANIRRVPFAFKWWRMVP----
>ENSBMSG00010001172/1-676
MGSPGSRAMSFQSIVHLSLDSPAHAICVQDMEIYLDVNGCAPQQCKSFTVDSSPGVVVQIYGTDPVKIGEEA
AMNRWPLSHPTNVLVGMASPSSANDKGKVSVSYYQHSEDVPMATAVLYLTSIGESL----------------
--QKNWVWGPKGWGAILLVNCSPADAHQL-VG-RKSKVFSTEETKSLSQMILRVQGPSCILKKCRVVLHTSK
EESEKARVYRPQKDSSSTFELVLGPDQHTYNLAP----VEDDLEETFYVEALEFPSASFSGLISYSASLVGE
SPDPSIPETVVYKDTVVFRVAPCVFVPSTQMPLEVYLCRELQVQGFVNTVMELSERSNIQVASVYEDPNRLG
RWLQDEMAFCYTQAPHKTISLVLDTPRLPKLDDFPMKYSLSPGVGYMTQRTQDHTVASIDSIGNLMVSPPVK
AQGKEYPLGRILIGSSFYPSKDCRNLSKTLRDFLYAQQVQAPVELFSDWLMIGHAYEFMCFIPAQYKVEDKK
DFRLLLASPSSCYKLFKEKQKEGYGDAHLFEGIRKDQLLSNGREANTINQLLADENMRKQNDYAEKCIDLNR
SILKRELGLEEQDIIDIPQLFCLERIANIPSSEQTEKLYARPYFPDLLQMIVMGQNLGIPKPFGPQINGTCC
LEEKIRQLLEPLGFQCTFINDFDCYLTEIGDFCSCANIRRVPFAFKWWRMVP----
>ENSOARG00020012854/1-688
--------MAFRNIVPLSLDSPTHAVCVLGVELFLDVSGCAPPACESFSVDASLGVVVRVCGADPVESREGP
AAPRWPLSRPTDVLVAMTSPNFAEDEGKVLVFYHQPKEDTPVATAVLHLTGVELSLDVDIYRSGHFE-ATDK
QAKRTWVWGPGGWGAILLVSCSPSAAGQP-GD--KNKVFSSEEVKSLSQMVLKVQGPSCITKNYRVVLHTSK
EESEKARVYRPQNDGSTAFELVLGPDQHTYTLAPQWDDRKGTLKETFYVEALEFPSASFSGLISFSASLVEE
SQDPLVPETVLYKDTVLFRVAPCIFAPTTQMPLEVYLCRELQVQGFVSAVTELSERSNSQVASVYEDPNRLG
RWLQDEMAFCYTQAPHKTVSLVLDTPRVAKPDDFPMKYSLSPGVGYLTLRTQDHTVASIDIIGKLMVSPPVK
AQGKEYPLGRVLIGSSFYPSKDSRNMSQSLQDFLRAQQVQAPVELFSDWLMTGHACEFMCFIPTQYKVEGKK
DFRLLLASPSSCYKLFKERQKEGYGDAMLFEGLRKDQLISNGREAVTINQLLADEKMRKHNDYAEKCIHLNR
SILKRELGLQEEDIIPIPQLFCLEHIANAPPSEQTKRLYARPYFPDLLQMVVMGLNLGIPKPFGPRVSGACC
LEERVCQLLEPLGFQCTFIDDFDCYLTEIGDFCSCASIRRVPFAFKWWGMVP----
>ENSCHIG00000014830/1-688
--------MAFRNIVPLSLDSPTHAVCVLGVELFLDISGCAPPACESFSVDASLGVVVRVCGADPVESREGP
AAPRWPLSCPTDVLVSMTSPNFAEDEGKVLVFYHQPKEDTPVATAVLHLTGVELSLDVDVYRSGRFE-ATDK
QAKKTWVWGPGGRGAILLVNCSPSVAGQP-VD--KNKVFSSEEVKSLSQMVLKVQGPSCITKNYRVVLHTSK
EESEKARVYRPQNDGSTAFELVLGPDQHTYTLAPQWDDRKGTLKETFYVEALEFPSASFSGLISFSASLVEE
SQDPLVPETVLYKDTVLFRVAPCIFAPTTQMPLEVYLCRELQVQGFVSAVTELSERSNSQVASVYEDPNRLG
RWLQDEMAFCYTQAPHKTVSLVLDTPRVAKPDDFPMKYSLSPGVGYLTLRTQDHTVASIDIIGKLMVSPPVK
AQGKEYPLGRVLIGSSFYPSKDSRNMSQSLQDFLRAQQVQAPVELFSDWLMTGHACEFMCFIPMQYKVEGKK
DFRLLLASPSSCYKLLKERQKEGYGDAMLFEGLRKDQLISNGREAVTINQLLADEKMRKQNDYAEKCIHLNR
SILKRELGLQEEDIIPIPQLFCLEHIANAPPSEQTKRLYARPYFPDLLQMVVMGLNLGIPKPFGPRVSGACC
LEERVCQLLEPLGFQCTFIDDFDCYLTEIGDFCSCASIRRVPFAFKWWGMVP----
>ENSBIXG00005026457/1-695
-VSPEGCAMAFRNIVPLSLDSPTHAVCVLGMEIFLDISGCAPRACESFSVDASLGVVVRVCGADPVESREKS
AAPRWPLSRPTDVLVAMTSPNFAEDEGKVMIFYHQPDEDAPMATAVLHLTGIELSLDVDIYRSGRFE-AADK
QAKKTWVWGPGGWGAILLVNCSPSVSGQP-VN--KSKVFSSEEVKSLSQMVLKVQGPSCITKNYRVVLHTSK
EESEKARVYRPQNDGSMTFELILGPEQHAYTLAPQWDEVKGTLRETFYVEALEFPSASFSGLISFSASLVEE
SQDPLVPEAVLCKDTVLFRVAPCIFVPSTQMPLEVYLCRELQVQGFVSTVTELSERSNSQVASVYEDPNRLG
RWLQDEMAFCYTQAPHKTVSLVLDTPRVAKLDDFPMKYSLSPGVGYLTLHAQDHTVASIDVIGKLMVSPPVK
AQGREYPLGRVLIGSSFYPSKDSRNMSRSLQDFLRAQQVQAPVELFSDWLMTGHACEFMRFIPAQYKVEGKK

DFRLLLASPSSCYKLFKERQKEGYGDAMLFEGLRKDQLLSNGREAITINQLLADEKMRKQNDYAEKCIHLNR
GILKRELGLQEEDIIPVPQLFCLEHIANAPPSEQTKKLYARPYFPDLLEMVVMGLNLGIPKPFGPRINGTCC
LEERICQLLEPLGFQCTFIDDFDCYLTEIGDFCSCANIRRVPFAFKWWGMVP----
>ENSBTAG00000038945/1-688
--------MAFRNIVPLSLDSPTHAVCVLGMEIFLDISGCAPRACESFSVDASLGVVVRVCGADPVESREKS
AAPRWPLSRPTDVLVAMTSPNFAEDEGKVMIFYHQPDEDAPMATAVLHLTGIELSLDVDIYRSGRFE-AADK
QAKKTWVWGPGGWGAILLVNCSPSVSGQP-VD--KSKVFSSEEVKSLSQMVLKVQGPSCITKNYRVVLHTSK
EESEKARVYRPQNDGSMTFELILGPEQHAYTLAPQWDEVKGTLRETFYVEALEFPSASFSGLISFSASLVEE
SQDPLVPEAVLCKDTVLFRVAPCIFVPSTQMPLEVYLCRELQVQGFVSTVTELSERSNSQVASVYEDPNRLG
RWLQDEMAFCYTQAPHKTVSLVLDTPRVAKLDDFPMKYSLSPGVGYLTLHTQDHTVASIDVVGKLMVSPPVK
AQGREYPLGRVLIGSSFYPSKDSRNVSRSLQDFLRAQQVQAPVELFSDWLMTGHACEFMRFIPAQYKVEGKK
DFRLLLASPSSCYKLFKERQKEGYGDAMLFEGLRKDQLLSNGREAITINQLLADEKMRKQNDYAEKCIHLNR
GILKRELGLQEEDIIPVPQLFCLEHIANAPPSEQTKKLYARPYFPDLLEMVVMGLNLGIPKPFGPRINGTCC
LEERICQLLEPLGFQCTFIDDFDCYLTEIGDFCSCANIRRVPFAFKWWGMVP----
>ENSBMUG00000003419/1-695
-VSPEGCAMAFRNIVPLSLDSPTHAVCVLGMEIFLDISGCAPRACESFSVDASLGVVVRVCGADPVESREKS
AAPRWPLLRPTDVLVAMTSPNFAEDEGKVMIFYHQPDEDAPMATAVLHLTGIELSLDVDIYRSGRFE-AADK
QAKKTWVWGPGGWGAILLVNCSPSVSGQP-MD--KSKVFSSEEVKSLSQMVLKVQGPSYITKNYRVVLHTSK
EESEKARVYRPQNDGSMTFDLILGPEQHAYTLAPQWDDVKGTLRETFYVEALEFPSASFSGLISFSASLVEE
PQDPLVPEAMLCKDTVLFRVAPCIFVPSTQMPLEVYLCRELQVQGFVSTVTELSERSNSQVASVYEDPNRLG
RWLQDEMAFCYTQAPHKTVSLVLDTPRVAKLDDFPMKYSLSPGVGYLTLHTQDHTVASIDVIGKLMVSPPVK
AQGREYPLGRVLIGSSFYPSKDSRNMSRSLQDFLRAQQVQAPVELFSDWLMTGHACEFMRFIPAQYKVEGKK
DFRLLLASPSSCYKLFKERQKEGYGDAMLFEGLRKDQLLSNGREAITINQLLADEKMRKQNDYAEKCIHLNR
GILKRELGLQEEDIIPVPQLFCLEHIANAPPSEQTKKLYARPYFPDLLEMVVMGLNLGIPKPFGPRINGTCC
LEERICQLLEPLGFQCTFIDDFDCYLTEIGDFCSCANIRRVPFAFKWWGMVP----
>ENSBGRG00000003075/1-635
--------MSLESLMP-----SNHLIL------------CRP-----------LLLLPSIFSSIRVFSNESA
LHIRWPKFWSFNFSISLSNESFRMDTLKSLLQHHSSKASIQIRTGV--------------FLS---------
SLQKTWVWGPGGWGAILLVNCSPSVSGQP-VD--KSKVFSSE---SLSQMVLKVQGPSCITKNYRVVLHTSK
EESEKARVYRPQNDGSMTFDLILGPEQHAYTLAPQWDDVKGTLRETFYVEALEFPSASFSGLISFSASLVEE
PQDPLVPEAVLCKDTVLFRVAPCIFVPSTQMPLEVYLCRELQVQGFVSTVTELSERSNSQVASVYEDPNRLG
RWLQDEMAFCYTQAPHKTVSLVLDTPRVAKLDDFPMKYSLSPGVGYLTLHTQDHTVASIDVIGKLMVSPPVK
AQGREYPLGRVLIGSSFYPSKDSRNMSRSLQDFLRAQQVQAPVELFSDWLMTGHACEFMRFIPAQYKVEGKK
DFRLLLASPSSCYKLFKERQKEGYGDAMLFEGLRKDQLLSNGREAITINQLLADEKMRKQNDYAEKCIHLNR
GILKRELGLQEEDIIPVPQLFCLEHIANAPPSEQTKKLYARPYFPDLLEMVVMGLNLGIPKPFGPRINGTCC
LEERICQLLEPLGFQCTFIDDFDCYLTEIGDFCSCANIRRVPFAFKWWGMVP----
>ENSBGRG00000020369/1-672
--------MYYKDL---------HA--VLGMEIFLDISGCAPRACESFSVDASLGVVVRVCGADPVESREKS
AAPRWPLSRPTDVLVAMTSPNFAEDEGKVLIFYHQPDEDAPMATAVLHLTGIELSLDVDIYRSGRFE-AADK
QAKKTWVWGPGGWGAILLVNCSPSVSGQP-VD--KSKVFSSEEVKSLSQMVLKVQGPSYITKNYRVVLHTSK
EESEKARVYRPQNDGPMTFELILGPEQHAYTLAPQWDDVKGTLRETFYVEALEFPSASFSGLTSFSASLVEE
SQDPLVPEAMLCKDTVLFRVAPCIFVPSTQMPLEVYLCRELQVQGFVSTVTELSERSNSQVASVYEDPNRLG
RWLQDEMAFCYTQAPHKTVSLVLDTPRVAKLDDFPMKYSLSPGVGYLTLHTQDHTVASIDVVGKLMVSPAVK
AQGREYPLGRVLIGSSFYPSKDSRNMSLSLQDFLRAQQVQAPVELFSDWLMTGHACEFMRFIPAQY-----K
DFRLLLASPSSCYKLFKERQKEGYGDAMLFEGLRKDQLLSNGRKAITINQLLADEKMRKQNDYAEKCIHLNR
GILKRELGLQEEDIIPVPQLFCLEHIANAPRSEQTKKLYAQPYFPDLLEMVVMGLNLGIPKPFGPRINGTCC
LEERICQLLEPLGFQCTFIDDFDCYLTEIGDFCSCANIRRVPFAFKWWGMVP----
>ENSMMSG00000007431/1-687
VVSLEVRAMAFRNIVPLSLDSPTHAVCVLGMEIFLDVSGCAPRTCESFSVDASLGVEVRICGAAPVESREEP
AATRWPLSRPTNVLVAMTSPNFAKDEGKVLVFYHQPDEDAPMATAVLHLTGIGECLPEPVFLS---------
SLQRTWVWGPGGWGAILLVNCSPSVAGQP-MD--QSKVFSS-EVKSLSQMVLKVQGPSCVIRNCRVTLHTSK
EESEKARVYRPQNDSSTTFELVLGPDRHTYTLAPQWDDVKDTLKETFYVEALEFPSASFSGLISFSASLVEE
SQDPLVPETVLYKDTVLFRVAPCIFVPSTQMPLEVYLCRELQVQGFVSTLVELSERSNSQVASVYEDPNRLG
RWLQDEMAFCYTQAPHKTISLVLDTPRAAKLDDFPMKYSLSPGVGYLTLRTRDHTVASIDTIGKLMVSPPVK
AQGREYPLGRVLIGSSFYPSKVSRDMSQSLRDFLRAQQVQAPVELFSDWLMTGHACEFMCFVPTQYKVEGKK
DFRLLLASPSSCYKLFEEKQKEGYGDVMLFEGLREDQLLSNAREAVTINQLLADTKMRRQNEYAEKCIHLNR
SILKKELGLQEEDIIPVPQLFCLEHVANAPSSEQTEKLYARPYFPDLVRMVVMGLNLGIPKPFGPRINGTCC
LEERICQLLEPLGFQCTFIDDFDCYLTEIGDFCSCANIRRVPFAFKWWGMVP----
>ENSCHYG00000003136/1-695
-ASLEGRAMPFRNVVPLSLDRPAHAVCVLGMEIFLDVSGCAPRSCESFSVDASLGVVVRVCSAAPVESKEEP
AATRWPLSSPKNVLVAMTSPNFAEDEGKVLVFYHQPDKDAPVATAVLHLTGIELSLDVDVYRKGRFE-TSDR
QAKKTWVWGPSGWGAILLVNCSPSVAGQP-VD--KSKVFSSEEVKSLSQMVLKVQGPSYIIKNYRVVLHTSK
EEAEKARVYRPQNNSSTTFELVLGPDQHTYTLAPQWDDVKDALKETFYVEALEFPSASFSGLISFSASLVEE

SQDPLVPETVLYKDSVLFRVAPCIFVPSTQMPLEVYLCRELQVQGFVSTVTELSERSNSQVASVYEDPNRLG
RWLQDEMAFCYTQAPHKTVSLVLDTPRVAKLDDFPMTYSLSPGVGYLTLRTQDHTVASIDAAGKLMVSPPVK
AQGKEYPLGRVLFGSSFYPSKDSRNMSRNLRDFLRAQQVQAPVELFSDWLMTGHACEFMCFIPTQYKVEGKK
DFRLLLASPSSCYKLFKEKQKEGYGDATLFEGLRKDQLLSNGREPVTINQLLADEKMRKQNDYVEKCIHLNR
SILKRELGLQEEDVIPIPQLFCLERVTNAPSSEQTEKLYARPYFPDLLQMVVMGLNLGIPKPFGPRVNGTCC
LEERVCRLLEPLGFQCTFIDDFDCYLTEIGDFCSCANIRRVPFAFKWWGMVP----
>ENSCWAG00000005012/1-695
AVSIEALAMSFQSVIHLSLDSPVHTVCVLGMEICLDLSGCAPQKCESFTISGSPGVLVDIYNIAPGKARQEA
ALTRWPLSHPMDVLVKMTSPSSTVNGDKVLVSYFLPDEEVPVATAILYLTGIEVSLDVDIYRNGDVEMASDK
HAKKNWVWGPDGWGAILLVNCTPADPGQL-KDGKSTKVFSTEEMKNLSQMVLRVQGPSSILKKHRLVLHTSK
DESDKARVYWPQKGSSSTFKLVLGPGQHSYSLGP----LENELKETFYVEALEFPSGSFSGLISFSASLVEE
PQDLAIPETLAYKDTVVFRVAPCIFIPSTQMPLEVYVCRELQVQGFVNAVTELSERSNSQVASVYEDPNRLG
RWLQDEMAFCYTQAPHKTVALVLDTPRVTKLDDFPMRYSLSPGVGYATQHTLDHTVASIDSVGNLVVSPPVQ
VEGKEYPLGRILIGSSCYPSKDGRDMSKALRDFLYAQQVQAPVELFSDWLMIGHPYEFMCFVPTQFKVEGKK
DFRLLLASPSTCYKLFMEKQKQGYGDAALFGEVRKEQLLANGREAFTINRLLADENMKKQNDYVERCISLNR
AILKRELGLAEKDIIDIPQLFCLEQLVNVPSNELTGKLYARPYFPDLLQIIVMGQNLGIPKPFGPQINGTCC
LEEKIYELLEPLGFQCTFINDFDCYLTEIGDFCSCANIRRVPFAFKWWKMVP----
>ENSSSCG00000003484/1-687
--------MSFQSIIHLSLDSPVHSTCMLGMEICLDLSGCAPQKCESFTIASSPGVLVDIYNVASGKVTQEA
ALTQWPLSHPMDVLVKMTSPSSDVNGDKVLVSYFLPHEDVPVATAILYLTGIEVSLDVDIYRNGDVEMANDK
QAKKNWVWGPDGWGAILLVNCALTNRGQI-KDGKITKVFSAEDIRNLSPMVLRVQGPSCILKKHRLLLHTSK
EESDKARVYRPQKDSSSTFELVLGPGQHAYALGP----LESDLKETFYVEALQFPSGSFPGLISYSVSLVEE
SKDGIGPQILVYKDTVVFRVAPCIFTPSTQMPLEMYVCRELQVQGFVNAVTELSERSNSQVASVYEDPNRLG
RWLQDEMAFCYTQAPHKTTALVLDTPRVTKLEDFPMRYSLSPGIGYVTLNTLDHTVASIDSIGNLMVSPPVK
VKGKEYPLGRILIGSSFYPSKDGRDMSKALRDFLYAQQVQAPVELFSDWLMIGHIYEFMCFIPTQFKVEGKK
DFRLLLASPSTCYKLFTEKQKQGYGDAALFGEIRKEQLLSNGREALTINQLLADENMKKQNDYVERCIDLNR
AILKRELGLAEKDIIDIPQLFCLEQIVNVPSNELTGKLYARPYFPNLLQIIVMGQNLGIPKPFGPQINGACC
LEEKIYELLEPLGFQCTFINDFDCYLTEIGDFCSCANIRRVPFAFKWWKMVP----
>ENSMLUG00000014390/1-656
-----NR--------------------------------CAPSKCTSFTISSSPRILLNLSHLVPAVTNKEG
ASTRLLLCEPMDVLVMVKFPSSDISEDKVSVSYYQPDVKVPVATAELFLTGIEVSLDVDIYRSGQVQVASDK
WAKKNWAWGPSGWGAILLVNCTPDGKAQD-GD--KTKVFLPEEINNLSPMMLSVQGPTCFLKTCRLVLYTSK
EEAKKARVYRPQDHSDTTFDLVLGPGCHTYTFPP----LESSLKETFYVEATEFPSASFSGLISYSVALVEE
SENTLIPETLMHKDMVVFRVAPCIFTPSTQMPLEIYLCRELQVQGFVKTVTELSEKSKSQVASVYEDPNRLG
RWLQDEMAFCYTQAPYKTISLVLDTPRVSNLEEFPMKYSLSPGIGYVTRPTEDGKVITMDSIGNLMVSPPVE
VKGKEYPLGRILIGGSFYPSKEARDMRKDLRDFLYAQQVQAPVELYSDWLMIGHMDQFMCFVPTQVKGEGDK
GFRLLLASPRSCYQLFQEKQEEGYGDMALFEEVREDQLLSNGREATTINQLLDDDDMRKQNDYVEKCINLNR
DILKRELGLLERDIIHIPQLFCLEQLSNVPATEQTEKLYARPYFPNLLQMIVMGKHLGIPKPFGPQIKGTCC
LEKRICQLLEPLGFKCTFINDFDCYMTEIGDFCACANVRRVPFAYKWWKMMP----
>ENSDNOG00000016263/1-692
---LEGGAMVFHDVICLSPDTPAHAVCMLGTEICLDLSGCAPQGCQSFTIVGSLGVLIHVYNTFPVKTSKEV
STTRWPLSDHMDVLVKMLSPSTAPDGDKVLVSYYQPDEEDPMATAVLYLTGIEVSLDVDIYRRGQVEMPSDP
HAKRKWTWGSSGWGAILLVNCNPANVGQP-PDTDTTKVFSSEEIKHLSQMTLSVQGPSCFLQKHRLVLHTSK
EESLKARVYWPQKDNPSVFELVLGPGQHNYALPC----LENRLKETFYVEAMEFPSADFSGLISYSVSLVEE
SQDLSIPEVLVYRDTVVFRVAPCIFTPSTQMPLEVYVCRELQLQGFVNTVMELSKKSNSQVASVYEDPNRLG
RWLQDEMAFCYTQAPHQTTSLVLDTPRVAKLEEFPMKYSLSPGTGYVTRRVEDCSAASMDSIGNLMVSPPVK
FQGKDYPLGRLLLGSSFYPSAEGRDMGTRLRDFLYAQQVQAPVELFSDWLTTGHVDEFMCFVPVDDKSEGAK
GFRLLLASPSSCYDLFQEKQKEGYGDAPLFEEVRAAQLLSNGREVKTIDQLLADDHLRKQNDYVERCIRLNR
DILRRELGLVEADIVHIPQLFCLEPLANVPSSQQPGKSFARPYFPDMLRMIVMGKNLGIPKPFGPQIKGACC
LEEKVCHLLEPLGFQCTFINDFDCYLTDVGDFCACANIRRVPFAFKWWKMLP----
>ENSLAFG00000010718/1-682
--------MSFQSIVHLSLDSPIQAICVLGTEICLDLHGCAPRECDSFTITGSLGVLTNVHNAVPIKTTEEA
ATTRWPLSGPMDVLVKMVSPSTAPSEDKVLVSYYEPDKEVPVSIAVLYLTGIEISLDVDIYRSGQVEMRSDK
QAKKKWVWGSSGWGAILLVNCNPASVGQL-TDKKTTK----PDIKSLSQMTLSVQGPSCTLKKHRLVLHTSR
EESEKARVYW-PQNSSSIFELVLGPGQHTYTLAP----LEDHLKKTFYIEAMEFPSADFSGLISYSVSLVDE
SQDPLIPEALVYKDTVVFRVAPCIFTPSTQMPLEVYICRELQLRGFVNTVMELSKKSNSQVASVYEDPNRLG
RWLQDEMAFCYTQAPHKTIPLVLDTPRVTKIEDLPMKYSLSSDVGYIIQGIKDYRVASMDSIGNLMVSPPVT
VQGKEYPLGRILIGSSFYPSEEGRNMSKTLQDFLHAQQVQAPVELFSDWLMVGHIDEFMCFVPIDERSEGEK
GFRLLLASPSSCYKLFEEKKKAGYGHMLLFEEVKVDQLLSNGREARTIDQLLADENMRNQNDYVEKCINLNR
DILKRELGLVEKDIIHIPQLFCLEQLTHISSNQQTEKRFARPYFPNMLQMIVMGKNLGIPKPFGPHIKGNCC
LEEKVCHLLEPLGFKCTFINDFDCYLTDVGDFRACANVRRVPFAFKWWRMMP----
